# Interaction between long-range chromatin regulators *Nipbl* & *Isl1* synergistically drives heart defects in mice

**DOI:** 10.1101/2025.05.15.654123

**Authors:** Stephenson Chea, Rosaysela Santos, Martha E. Lopez-Burks, Arthur D. Lander, Anne L. Calof

**Author notes:** These authors contributed equally to this work (Stephenson Chea, Rosaysela Santos).

## Abstract

Congenital heart defects (CHDs) are frequently observed in the most common form of Cornelia de Lange Syndrome (CdLS), which is caused by haploinsufficiency for *NIPBL*, a gene involved in chromatin looping and cis-regulatory control of gene expression. Here, we surveyed cardiac defects in mice made *Nipbl*-haploinsufficient in the second heart field using two Cre drivers: *Mef2c-Cre* and *Isl1-Cre*. Only *Isl1-Cre*-driven *Nipbl*-haploinsufficiency resulted in CHDs – a finding we traced to the additional contribution of *Isl1*-haploinsufficiency caused by the *Isl1-Cre* allele. To test whether combined reduction of *Nipbl* and *Isl1* cause CHDs, we made mice globally haploinsufficient for both genes. Indeed, *Nipbl^+/-^; Isl1^+/-^* mice exhibited a substantially higher frequency and severity of CHDs than mice haploinsufficient for either gene alone. As a member of the LIM-homeodomain transcription factor family, *Isl1* is involved in chromatin looping and enhancer-promoter communication via a mechanism distinct from that of *Nipbl*. Nevertheless, when we performed RNA sequencing on E10.5 hearts from wildtype, *Nipbl^+/-^*, *Isl1^+/-^*, and *Nipbl^+/-^; Isl1^+/-^* embryos, we observed that combined haploinsufficiency resulted in largely additive gene expression changes, including dysregulation of known cardiac regulators (*Irx4, Tbx1, Foxo6, Heyl, Bnc1, Sox17*) and novel candidates (*Gbx1, Csdc2, Myrf, Pou6f1, Zfp579, ad Zfp763*). A subset of additive changes arose from *opposing* regulatory influences in single mutants that restored gene expression to WT levels in *Nipbl⁺^/^⁻; Isl1⁺^/^⁻* hearts. For example, *Hoxc4, Pitx2, Isl1* itself, and *Pax6* (a known target of *Isl1*), were upregulated in *Nipbl^+/-^*hearts, downregulated in *Isl1^+/-^* hearts, but expressed at WT levels in *Nipbl⁺^/^⁻; Isl1⁺^/^⁻* hearts. Since loss of *Isl1* upregulation from *Nipbl^+/-^* to *Nipbl^+/-^; Isl1^+/-^* hearts coincided with a marked increase in CHDs, we propose that *Isl1* upregulation compensates for the loss of cis-regulatory interactions due to *Nipbl*-haploinsufficiency, and protects hearts from severe CHD risk. Supporting this model, other LIM-homeodomain transcription factors (*Lhx2, Lhx3, Lhx9*) were also upregulated in *Nipbl^+/-^* hearts, with *Lhx3* and *Lhx9* showing even greater upregulation in *Nipbl^+/-^; Isl1^+/-^* hearts. Despite this, CHDs resulting from the combined loss of *Nipbl* and *Isl1* were particularly severe. These findings suggest that heart development is exquisitely sensitive to small changes in gene expression, leading to synergistic phenotypic interactions when relatively modest gene expression changes are combined.

## INTRODUCTION

Congenital heart defects (CHDs) affect nearly 1% of all newborns in the United States and are the leading cause of birth defect-associated infant mortality and morbidity (Hoffman and Kaplan, 2002, Oster et al., 2013, Mai et al., 2019). The causes of CHDs are largely unknown, with mutations in single genes accounting for less than 20% of cases (Richards and Garg, 2010). The majority of CHDs are thought to arise from multifactorial and/or polygenic origins (Priest et al., 2016, Ware and Jefferies, 2012), which has complicated the creation of animal models.

Studies of Cornelia de Lange Syndrome (CdLS) have shed some light on causal factors for CHDs (Santos et al., 2016). CdLS is a multisystem birth defects disorder that occurs in approximately 1 in 10,000 to 1 in 30,000 live births (Mannini et al., 2013, Boyle et al., 2015). About 30% of individuals with CdLS have CHDs, with ventricular septum defects (VSD) and atrial septum defects (ASD) being the most common (Chatfield et al., 2012).

Most CdLS cases are caused by haploinsufficiency for the gene *Nipped-B-like* (*NIPBL*) (Gillis et al., 2004). *NIPBL* encodes a universally-conserved protein that plays a role in loading the cohesin complex onto chromosomes (Ciosk et al., 2000, Gillespie and Hirano, 2004, Takahashi et al., 2004, Watrin et al., 2006). While cohesin is widely recognized for its role in maintaining sister chromatid cohesion during mitosis (Nasmyth and Haering, 2009), sister chromatid cohesion is unaffected in people with CdLS and the *Nipbl^+/-^* murine model of CdLS (Kawauchi et al., 2009).

In recent years, the cohesin complex has been increasingly recognized for its role in regulating global gene expression through chromatin loop-mediated cis-regulatory interactions (Alonso-Gil and Losada, 2023). That Nipbl levels are crucial to these interactions is supported by studies showing that alterations in the expression of hundreds to thousands of genes in all examined tissues are found in CdLS patients and *Nipbl*-deficient animal models (Liu et al., 2009, Kawauchi et al., 2009, Muto et al., 2011, Wu et al., 2015). Interestingly, most of the gene expression changes are small, typically less than 2-fold change. Though likely inconsequential individually, collectively, these changes contribute to the structural and functional defects seen in CdLS (Muto et al., 2011). Thus, CdLS exemplifies an emerging class of genetic disorders – recently termed “transcriptomopathies” – in which quantitative variations in multiple gene product levels lie at the root of developmental abnormalities. Such disorders provide a window into the kinds of multifactorial interactions potentially underlying polygenic traits, including that of CHDs.

While most attention has focused on cohesin and its loading factor *Nipbl*, emerging evidence suggests that other molecular systems also participate in chromatin looping and long-range enhancer-promoter communication. These include LIM-homeodomain (LIM-HD) transcription factors such as *Isl1* and *Lhx* family members, which interact with the adaptor protein *Ldb1* to mediate looping between regulatory elements (Bower and Kvon, 2025). *Ldb1* facilitates this process by binding to the LIM domains of LIM-HD proteins – a binding interaction made possible when the homeodomains of these transcription factors are anchored to chromatin (Caputo et al., 2015, Krivega et al., 2014). In the present study, we show that haploinsufficiency for the LIM-HD transcription factor *Isl1* also results in modest changes in gene expression, though the changes are less numerous than those obseved in *Nipbl*-haploinsufficient tissues. Such results support the view that LIM-HD proteins represent a distinct class of chromatin regulators capable of influencing gene expression through effects on chromatin architecture (Bower and Kvon, 2025).

Among the features of CdLS that are phenocopied in *Nipbl*-deficient animal models are CHDs. In *Nipbl*^+/-^ mice, these include atrial septal defects (ASDs) (Santos et al., 2016), which occur at the same frequency as in CdLS individuals (30%) (Chatfield et al., 2012); they are also among the most common kind of non-syndromic CHDs in humans (Botto et al., 2001). Thus, the *Nipbl^+/-^* mouse serves as an excellent model for studying the developmental origins of CHDs. Intriguingly, recent studies in *Nipbl^+/-^* mice have found that ASDs are preceded by heart abnormalities at earlier stages of development: these include incomplete fusion of the ventricular septum with the cardiac cushion in 77% of hearts at E13.5, smaller right ventricles in 100% of hearts at E10.5, and reduced expression of early cardiac transcription factors, *Nkx2.5* at E7.5 and *Mesp1* at E7.5 (Santos et al., 2016).

The mammalian heart arises from multiple tissue lineages: the heart is primarily derived from the mesoderm, one of the three germ layers in early embryogenesis (Gittenberger-de Groot et al., 2005). Within mesoderm, the first cardiac cells to differentiate form the initial cardiac crescent and are termed the first heart field (FHF) (Kelly et al., 2014). Later, cells from the second heart field (SHF) additionally contribute to the cardiac crescent, followed by elongation and looping of the heart tube, giving rise to the right ventricle (but not the left), outflow tract, atria, and atrial/ventricular septa (Kelly et al., 2014). Neural crest cells, which originate from ectoderm, also play a role in development of the outflow tract by contributing to division of the primitive heart tube into the aorta, pulmonary trunk, and heart valves (Keyte and Hutson, 2012). Collectively, these diverse tissue lineages orchestrate heart morphogenesis.

In a previous study, we asked in which cell lineages *Nipbl*-haploinsufficiency is necessary and sufficient to cause CHDs (Santos et al., 2016). In that study, we used a *Nipbl* allelic series containing a conditional/invertible gene trap and Cre recombinase expressing mouse lines, to create *Nipbl*-haploinsufficiency in neural crest cells (*Wnt-Cre*) and cells of the cardiac crescent (*Nkx2-5-Cre*). *Nipbl*-haploinsufficiency in neural crest cells alone did not result in CHDs (Santos et al., 2016). However, *Nipbl-*haploinsufficiency in cardiac crescent cells, which consist of both FHF and SHF cells, did result in CHDs (Santos et al., 2016). Interestingly, the heart abnormalities observed in *Nipbl^+/-^* mice (which include ASDs, incomplete fusion of the ventricular septum, and right ventricle hypoplasia) are found in tissues primarily derived from the SHF. Other studies have demonstrated that disruptions in the second heart field can result in outflow tract defects (Vincent and Buckingham, 2010), atrial and ventricular septal defects (Buckingham et al., 2005), and right ventricular hypoplasia (Rochais et al., 2009). Collectively, these findings suggested the SHF may be a potential source of CHDs in *Nipbl^+/-^* mice.

In this study, we used light sheet fluorescence microscopy (LSFM) to look for CHDs in mice made *Nipbl*-haploinsufficient specifically in the SHF using two Cre drivers: *Mef2c-Cre* (Verzi et al., 2005) and *Isl1^Cre/+^* (Yang et al., 2006). Surprisingly, only *Isl1^Cre/+^* driven *Nipbl*-haploinsufficiency (*Nipbl^Flox/+^; Isl1^Cre/+^*) resulted in CHDs, whereas *Mef2c-Cre* did not. While this difference may reflect differences in Cre activity – such as differences in timing, efficiency, or pattern of expression (e.g., in extracardiac tissues), we suspected and subsequently showed that *Isl1^Cre/+^* mice are themselves haploinsufficient for *Isl1* and develop CHDs themselves. These findings raised the possibility that the CHDs observed in *Nipbl^Flox/+^; Isl1^Cre/+^* mice result from the combined effects of reduced *Nipbl* and *Isl1* dosage within the *Isl1-Cre* expression domain. Below we show that this is the case, and investigate the relationship between the transcriptional effects of *Nipbl*- and *Isl*- haploinsufficiency in causing CHDs.

## RESULTS

### LSFM enhances sensitivity for detecting previously unrecognized CHDs in *Nipbl^+/-^* mice

Looking for CHDs in mouse hearts by histological analysis, is both time-consuming and relatively destructive to samples. In a previous study, we introduced micro-magnetic resonance imaging (µMRI) as an alternative method for CHD detection (Santos et al., 2016), but a limitation of µMRI is its low spatial resolution, which can blur fine anatomical details. To overcome the limitations of both methodologies, we turned to LSFM, which has enabled rapid, high-resolution three-dimensional imaging of other biological samples with minimal phototoxicity, preserving sample integrity. Bypassing the need for invasive tissue sectioning, LSFM avoids potential artifacts and provides an almost 10-fold improvement in spatial resolution over that of µMRI (LSFM: 6-7 µm voxel resolution versus µMRI: 50-70 µm). Furthermore, when combined with fluorescent reporter lines, LSFM also offers insights into tissue-specific dynamics that may contribute to the development of CHDs.

As a first step, we sought to validate that LSFM could reliably detect the same heart defects previously identified by histological analysis and µMRI in *Nipbl^+/-^* mice (Kawauchi et al., 2009, Santos et al., 2016). Using a *Nipbl* allelic series containing a conditional/invertible gene trap previously developed (Santos et al., 2016), we generated WT (*Nipbl^Flox/+^*) and *Nipbl^+/-^* (*Nipbl^FIN/+^*) mouse littermates on a *Td-tomato-EGFP* fluorescent reporter background in which cells expressing Cre recombinase is reported by enhanced green fluorescent protein (EGFP) expression. Global *Nipbl*-haploinsufficiency was achieved using the *Nanog-Cre* line (Economides et al., 2013), in which Cre recombinase is expressed in the earliest cells of the developing embryo – the inner cell mass. We also generated WT and *Nipbl^+/-^* littermates without the *Td-tomato-EGFP* fluorescent reporter background, since LSFM can also image tissues using autofluorescence.

LSFM detected a single ASD in WT mice and a *spectrum* of CHDs in *Nipbl^+/-^* mice. Previously, only ASDs were detected in *Nipbl^+/-^* mice by histological analysis or µMRI at the same stage (Santos et al., 2016). The spectrum of CHDs detected by LSFM included ASDs, VSDs, and outflow tract defects (OTDs), such as overriding aorta (OA) (Fig. 1A & Table S1), double outlet right ventricle (DORV) (Fig. 1B & Table S2), and transposition of the great arteries (TGA) (not shown). These OTDs always occurred in conjunction with either an ASD or both an ASD and VSD (Fig. 1C & Table S3). Despite this expanded spectrum, ASDs remained the most common CHD detected by LSFM, occurring in 80% of *Nipbl^+/-^* mice with CHDs. In total, LSFM detected CHDs in 36.59% of *Nipbl^+/-^*hearts, substantially higher than what was detected in WT hearts (1.82%) (Fig. 1C), and approximately 7% higher than what was previously detected by histological analysis and µMRI (∼30%) (Santos et al., 2016). Since we previously observed that *Nipbl^+/-^* mice have smaller hearts than WT mice (Santos et al., 2016), we also used LSFM to measure the ventricular volumes of hearts from E17.5 WT and *Nipbl^+/-^* mice. *Nipbl^+/-^* hearts were 28% smaller than WT hearts (Fig. 1D & Table S4). These results validate LSFM as a sensitive, high-resolution tool for detecting CHDs, and reveal that CHDs in *Nipbl^+/-^* mice extend beyond ASDs to include previously underrecognized abnormalities in ventricular septation and outflow tract development.

**Fig. 1.**
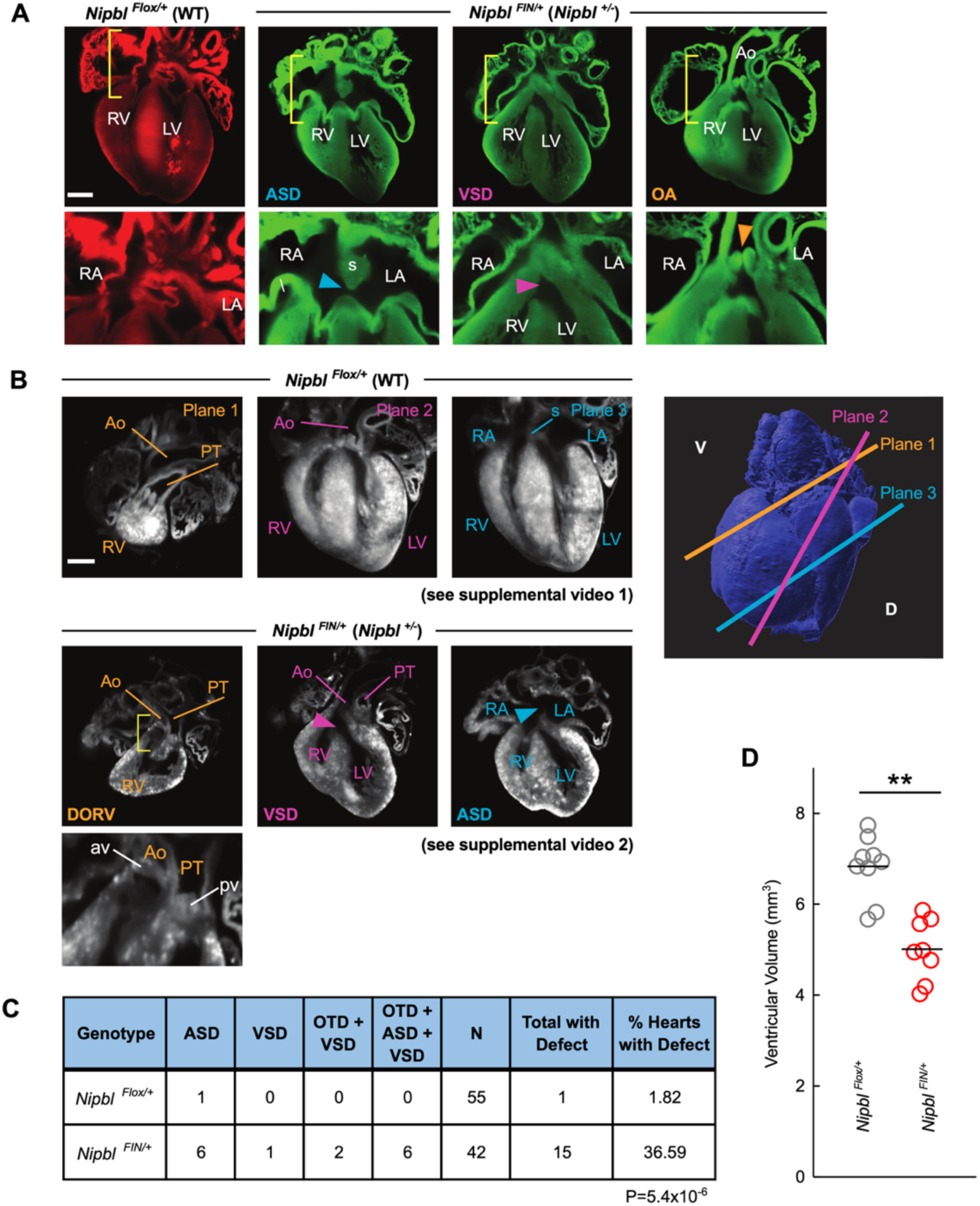
Light sheet fluorescence microscopy detects a spectrum of CHDs in *Nipbl^+/-^* mice. (A) LSFM images of one representative *Nipbl^Flox/+^* (WT) E17.5 heart and two different *Nipbl^FIN/+^* (*Nipbl*^+/-^) E17.5 hearts on a *Td-tomato-EGFP* fluorescent reporter background. *Nanog*-*Cre*- mediated recombination is reported by EGFP expression. *Nipbl^Flox/+^*mice showed no recombination. All tissues showed recombination in *Nipbl^FIN/+^* mice. *Nipbl^FIN/+^* hearts display a spectrum of heart defects (from left to right, panel A1 shows a normal *Nipbl^Flox/+^*heart; panels A2 & A3 show two different defects in the same *Nipbl^FIN/+^* heart; panel A4 shows overriding aorta in a different *Nipbl^FIN/+^* heart). Panels in bottom row show higher magnification of areas indicated by yellow brackets in panels in top row. In high magnification panels, blue arrowhead points to atrial septal defect (ASD), pink arrowhead points to ventricular septal defect (VSD), and orange arrowhead points to overriding aorta (OA). RV, right ventricle; LV, left ventricle; Ao, aorta; RA, right atrium; LA, left atrium; S, atrial septum; PT, pulmonary trunk. Scale bar = 500 um. (B) LSFM images of one representative *Nipbl^Flox/+^*E17.5 heart and one representative *Nipbl^FIN/+^* E17.5 heart without fluorescence reporting of *Cre*-mediated recombination. Images are of three planes of one *Nipbl^Flox/+^*and one *Nipbl^FIN/+^* heart. The positions of these planes in the heart are shown in the schematic to the right. From left to right, images show the plane progressing from ventral to dorsal in the heart. The *Nipbl^FIN/+^* heart displays a spectrum of heart defects. Bottom left panel shows higher magnification of area indicated by yellow bracket in panel above it, revealing double outlet right ventricle (DORV). RV, right ventricle; LV, left ventricle; Ao, aorta; RA, right atrium; LA, left atrium; S, atrial septum; PT, pulmonary trunk. Videos of LSFM images through whole *Nipbl^Flox/+^*and *Nipbl^FIN/+^* hearts can be viewed in Videos S1 and S2, respectively. (C) Table of frequency of heart defects in E17.5 *Nipbl^Flox/+^*and *Nipbl^FIN/+^* hearts as detected by LSFM. ASD, atrial septal defect; VSD, ventricular septal defect. Hearts showing outflow tract defects (OTD) include one or more of the following: overriding aorta (OA), double outlet right ventricle (DORV), transposition of the great arteries (TGA). P-value from Fisher’s Exact Test. (D) Ventricular volumes of E17.5 *Nipbl^Flox/+^* (N=8) and *Nipbl^FIN/+^*(N=9) hearts. Ventricular volumes were measured in ImageJ using LSFM images of hearts. Bars show means. ** represents P ≤ 0.01. P-value from Mann-Whitney U Test.

### *Nipbl-*haploinsufficiency in the *Mef2c-Cre* domain does not result in CHDs

To investigate whether *Nipbl*-haploinsuffiency restricted to the SHF results in CHDs, we generated E17.5 mice that were *Nipbl*-haploinsufficient specifically in the SHF using the *Mef2c- Cre* mouse line and that were on a *Td-tomato-EGFP* fluorescent reporter background. In these mice, tissue expressing Cre recombinase are *Nipbl*-haploinsufficient and express EGFP. As expected, EGFP expression was observed in SHF derivatives, including the right ventricle and parts of the atria (Fig. 2A). Surprisingly, only 4.5% of *Nipbl^Flox/+^; Mef2c-Cre* mice showed a CHD, and when they did, they only showed ASDs, a result that was not significantly different from WT (*Nipbl^Flox/+^*) mice (Fig. 2, A to B & Table S5). They also had hearts of the same size as WT mice (Fig. 2C & Table S6). These results indicate that *Nipbl*-haploinsufficiency restricted to the SHF is largely insufficient to disrupt heart development, suggesting that *Nipbl* function in non-SHF lineages plays a critical role in supporting normal cardiac morphogenesis.

**Fig. 2.**
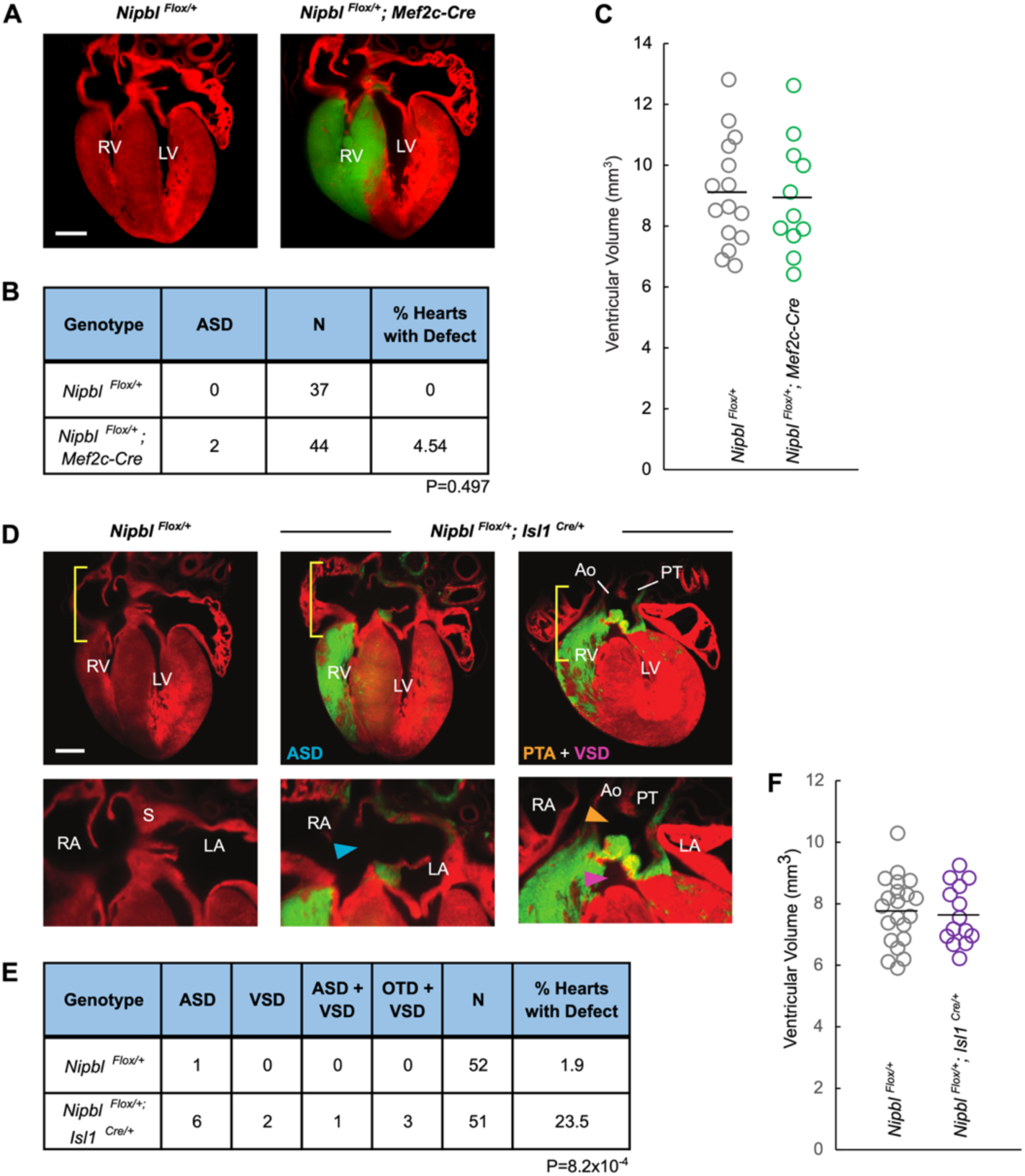
*Nipbl* made deficient in the SHF domain using one Cre driver (*Isl1-Cre*), but not another (*Mef2c-Cre*), results in CHDs. (A) LSFM image of representative E17.5 *Nipbl^Flox/+^* and *Nipbl^Flox/+^*; *Mef2c-Cre* heart on a *Td-tomato-EGFP* fluorescent reporter background showing normal heart development. *Mef2c-Cre* mediated recombination is reported by EGFP expression. RV, right ventricle; LV, left ventricle. Scale bar = 500 uM. (B) Table of frequency of atrial septal defects (ASDs) in E17.5 *Nipbl^Flox/+^* (N=37) and *Nipbl^Flox/+^*; *Mef2c-Cre* (N=44) hearts, as detected by LSFM (N=20, *Nipbl^Flox/+^*, N=21, *Nipbl^Flox/+^*; *Mef2c-Cre*) and histological analysis (N=17, *Nipbl^Flox/+^*, N=23, *Nipbl^Flox/+^*; *Mef2c-Cre*). P-value from Fisher’s Exact Test. (C) Ventricular volumes of E17.5 *Nipbl^Flox/+^* (N=15) and *Nipbl^Flox/+^*; *Mef2c-Cre* (N=11) hearts. Ventricular volumes were measured in ImageJ using LSFM images of hearts. Bars show means. (D) LSFM images of one representative *Nipbl^Flox/+^*E17.5 heart and two different *Nipbl^Flox/+^*; *Isl1^Cre/+^* E17.5 hearts on a *Td-tomato-EGFP* fluorescent reporter background. *Isl1*-*Cre* mediated recombination is reported by EGFP expression. *Nipbl^Flox/+^*; *Isl1^Cre/+^* hearts display a spectrum of heart defects. Panels in bottom row show higher magnification of areas indicated by yellow brackets in panels in top row. In high magnification panels, arrowheads point to atrial septal defect (ASD, blue), ventricular septal defect (VSD, pink), and overriding aorta (OA, orange). RV, right ventricle; LV, left ventricle; Ao, aorta; PT, pulmonary trunk; RA, right atrium; LA, left atrium; S, atrial septum. Scale bar = 500 uM. (E) Table of frequency of heart defects in E17.5 *Nipbl^Flox/+^* (N=52) and *Nipbl^Flox/+^*; *Isl1^Cre/+^* (N=51) hearts, as detected by LSFM. Of the three *Nipbl^Flox/+^*; *Isl1^Cre/+^* hearts showing both an outflow tract defect and ventricular septal defect (OTD + VSD), two showed an overriding aorta OTD and one showed a persistent truncus arteriosus OTD. P-value from Fisher’s Exact Test. (F) Ventricular volumes of E17.5 *Nipbl^Flox/+^* (N=21) and *Nipbl^Flox/+^*; *Isl1^Cre/+^* (N=14) hearts. Ventricular volumes were measured in ImageJ using LSFM images of hearts. Bars show means.

### *Nipbl-*haploinsufficiency in the *Isl1-Cre* domain generates a spectrum of CHDs

Given that *Mef2c-Cre*–mediated *Nipbl*-haploinsufficiency within the SHF did not result in a significant production of CHDs, we next asked whether using an alternative SHF-specific Cre line, *Isl1-Cre –* which is expressed a few hours earlier *–* might better model this phenotype (Dodou et al., 2004). To test this, we generated E17.5 mice SHF-restricted *Nipbl*- haploinsufficiency using the *Isl1^Cre/+^* mouse line crossed onto a *Td-tomato-EGFP* fluorescent reporter background, allowing Cre activity to be monitored by EGFP expression. As expected, EGFP expression was observed in SHF derivatives, including the right ventricle and parts of the atria (Fig. 2D). *Nipbl^Flox/+^; Isl1^Cre/+^* mice developed a significant number of CHDs (23.5%) compared to WT (*Nipbl^Flox/+^*) mice, which did not display any (Fig. 2E & Table S7). Although the frequency of CHDs displayed by *Nipbl^Flox/+^; Isl1^Cre/+^* mice was lower than that of globally *Nipbl*-haploinsufficient (*Nipbl^+/-^)* mice (∼37%) (Fig. 1C), the spectrum of defects was similar, which included: ASDs, VSDs, and OTDs, including OA and persistent truncus arteriosus (PTA) (Fig. 2D). As in *Nipbl^+/-^* mice (Fig. 1C), OTDs in *Nipbl^Flox/+^; Isl1^Cre/+^* mice occurred in conjunction with VSDs (Fig. 2E). Unlike, *Nipbl^+/-^* mice, however, which showed smaller hearts than WT (Fig. 1D), heart size was unaffected in *Nipbl^Flox/+^; Isl1^Cre/+^* mice compared to WT (*Nipbl^Flox/+^*) (Fig. 2F & Table S8). Thus, while the frequency of CHDs in *Isl1-Cre* mediated *Nipbl*-haploinsufficiency in the SHF leads to a spectrum of CHDs reminiscent of globally *Nipbl*- haploinsufficient mice.

### *Mef2c-Cre* and *Isl1-Cre* define similar expression domains within the heart, but *Isl1-Cre* also recombines extensively in non-cardiac tissues

Given the surprising difference in CHD outcomes between *Mef2c-Cre* and *Isl1-Cre* drivers, we next investigated whether differences in their recombination domains could account for the divergent phenotypes. To do so, we mapped Cre recombination patterns across embryonic tissues from E8.5 to E17.5 using Td-tomato-EGFP reporter mice, where Cre activity is marked by EGFP expression. Both *Mef2c-Cre* and *Isl1-Cre* drove recombination in SHF derivates at all examined stages: in the cardiac crescent at E8.5, the developing right ventricle and outflow tract at E10.5, and the ventricular and atrial septa, aorta, and pulmonary trunk at E15.5-E17.5 (Fig. 3, A & B). With exception of the cardiac crescent at E8.5, *Mef2c-Cre* recombined more extensively within the heart compared to *Isl1-Cre* at all stages. Based on this more extensive recombination, one might have expected that having an expanded domain of *Nipbl*-haploinsufficiency within *Nipbl^Flox/+^; Mef2c-Cre* hearts would result in more CHDs than *Nipbl^Flox/+^; Isl1^Cre/+^* hearts, but this was the opposite of what was observed (Fig. 2).

**Fig. 3.**
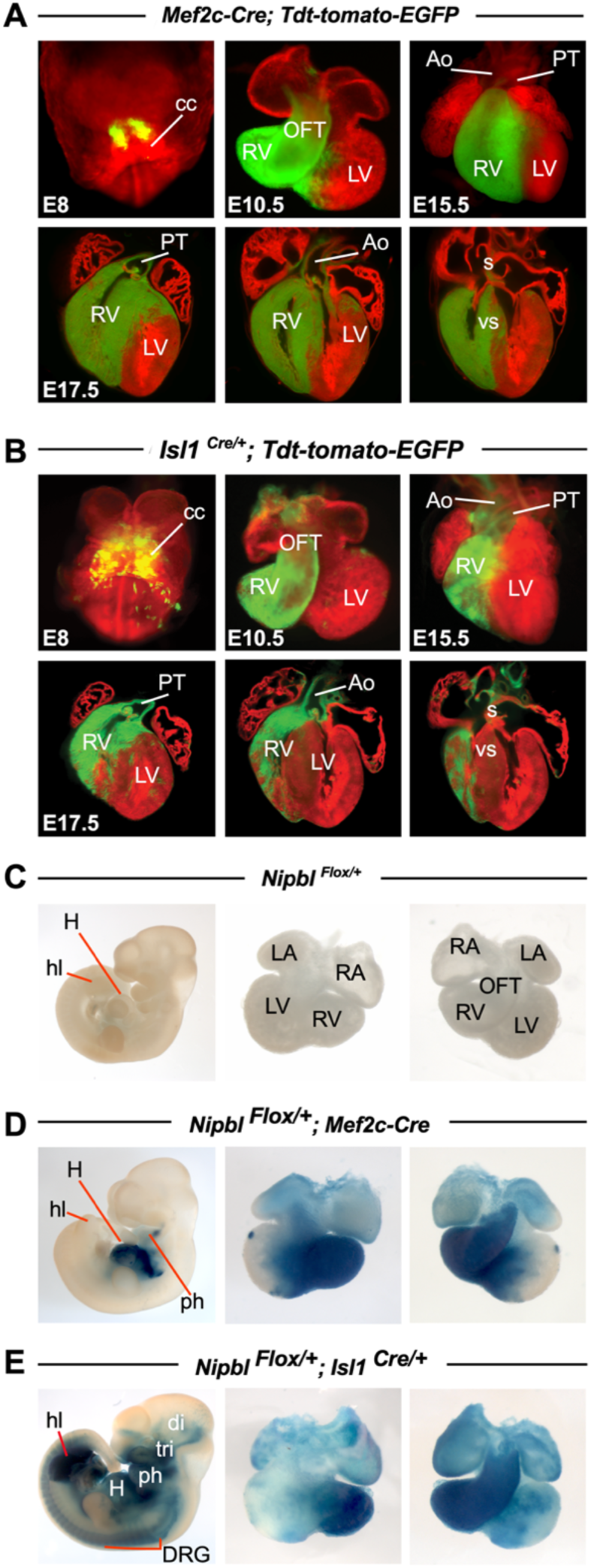
*Mef2c-Cre* and *Isl1-Cre* recombine similarly within the heart, but *Isl1-Cre* recombines extensively in non-cardiac tissues. LSFM images of representative (A) *Mef2c-Cre* and (B) *Isl1^Cre/+^* E8.5 whole embryos, and E10.5, E15.5, and E17.5 hearts on a *Td-tomato-EGFP* fluorescent reporter background, in which Cre recombination is reported by EGFP expression. Bottom row shows three different planes of one E17.5 heart progressing from ventral to dorsal from left to right. CC, cardiac crescent; RV, right ventricle; LV, left ventricle; OFT, outflow tract; Ao, aorta; PT, pulmonary trunk; S, atrial septum; VS, ventricular septum. X-gal stains of representative E10.5 (C) *Nipbl^Flox/+^*, (D) *Nipbl^Flox/+^*; *Mef2c-Cre*, and (E) *Nipbl^Flox/+^*; *Isl1^Cre/+^* embryos and hearts. H, heart; hl, hindlimb; LA, left atrium; RA, right atrium; LV, left ventricle; RV, right ventricle; OFT, outflow tract; DRG, dorsal root ganglia; ph, pharyngeal arches; tri, trigeminal ganglia; di, diencephalon.

In a previous study, we showed that *Nipbl*-haploinsufficiency in *non-cardiac* tissues can contribute to the development of CHDs (Santos et al., 2016). We therefore analyzed the recombination domains of Cre-mediated *Nipbl*-haploinsufficiency across the whole bodies of *Nipbl^Flox/+^; Mef2c-Cre* and *Nipbl^Flox/+^; Isl1^Cre/+^* mouse embryos at E10.5, the earliest stage before which CHDs were observed in mice of either genotype. In both genotypes, Cre-recombination was reported by lacZ expression. No recombination was observed in WT (*Nipbl^Flox/+^*) control mice (Fig. 3C). In *Nipbl^Flox/+^; Mef2c-Cre* mice, *Cre* recombination was observed outside the heart in the pharyngeal arches (Fig. 3D). In *Nipbl^Flox/+^; Isl1^Cre/+^* mice, Cre recombination extended to additional tissues including the hindlimbs, dorsal root ganglia, pharyngeal arches, trigeminal ganglia, and diencephalon. Thus, while both Cre lines target SHF derivatives, *Isl1-Cre* drives broader recombination in extracardiac tissues compared to *Mef2c-Cre*. Given that *Isl1-Cre* also recombines less extensively *within* the heart than *Mef2c* (Fig. 3, A & B), these results suggest that *Nipbl*-haploinsufficiency affecting more extracardiac tissues may contribute to the increased CHD burden observed in *Nipbl^Flox/+^; Isl1^Cre/+^* mice, even when a smaller fraction of the heart is *Nipbl*-haploinsufficient compared to *Nipbl^Flox/+^; Mef2c-Cre* mice.

### *Isl1*-haploinsufficiency alone results in CHDs

*Isl1-Cre* is a knock-in allele that targets the start codon of the endogenous *Isl1* locus (Yang et al., 2006), whereas *Mef2c-Cre* was produced using a bacterial artificial chromosome (BAC) as a transgene, leaving the endogenous gene intact (Verzi et al., 2005). Given this, *Isl1^Cre/+^* mice are expected to be haploinsufficient for *Isl1*. Indeed, qRT-PCR of E10.5 hearts showed that *Isl1* expression in *Isl1^Cre/+^* hearts was reduced by 71% compared to WT (*Isl1^+/+^*) littermates (Fig. 4D, Table S11, & Table S12). Interestingly, 22% of E17.5 *Isl1^Cre/+^* mice exhibited CHDs, significantly higher than the 2% frequency observed in WT controls mice (Fig. 4, A to B, & Table S9). ASDs were the only type of defect observed in *Isl1^Cre/+^* mice, and heart size remained unaffected (Fig. 4C & Table S10). Thus, these findings indicate that *Isl1*- haploinsufficiency alone results in CHDs, and identify *Isl1* as a dosage-sensitive regulator of atrial septation.

**Fig. 4.**
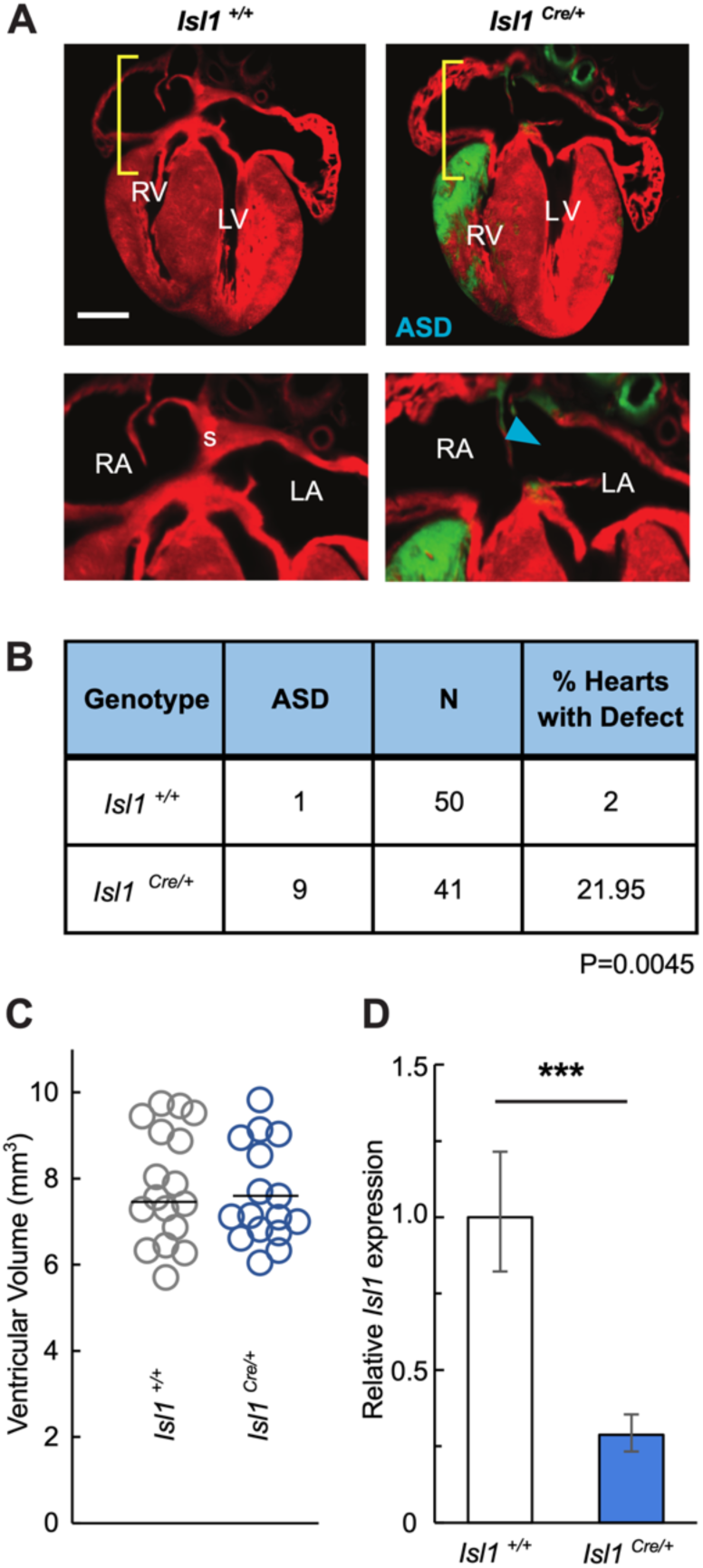
CHDs and reduced *Isl1* expression in *Isl1^Cre/+^* hearts. (A) LSFM images of representative E17.5 *Isl1*^+/+^ and *Isl1^Cre/+^* hearts on a *Td-tomato-EGFP* fluorescent reporter background. Panels in bottom row show higher magnification of areas indicated by yellow brackets in panels in top row. In high magnification panels, blue arrowhead points to atrial septal defect (ASD). RV, right ventricle; LV, left ventricle; RA, right atrium; LA, left atrium; S, atrial septum. (B) Table of frequency of atrial septal defects (ASDs) in E17.5 *Isl1*^+/+^ (N=50) and *Isl1^Cre/+^* (N=41) hearts, as detected by LSFM. P-value from Fisher’s Exact Test. (C) Ventricular volumes of E17.5 *Isl1*^+/+^ (N=17) and *Isl1^Cre/+^* (N=16) hearts. Volumes were measured in ImageJ using LSFM images of hearts and summed. Bars show means. (D) Relative expression of *Isl1* (normalized to *Rpl4* expression) in E10.5 *Isl1*^+/+^ (N=9) and *Isl1^Cre/+^* (N=9) hearts as measured by quantitative reverse transcription PCR. Error bars show standard error of the mean. *** represents P ≤ 0.001. P-value from T-Test.

### Mice doubly haploinsufficient for *Nipbl* and *Isl1* show fetal demise and reduced body size

Since *Nipbl*-haploinsuffiency in the *Isl1-Cre* expression domain results in CHDs and *Isl1*- haploinsufficiency alone also results in CHDs, we sought to understand the impact of *Isl1*- haploinsufficiency on heart development in the context of global (i.e., embryo-wide) *Nipbl*- haploinsufficiency. To investigate this, we used *Stra8-iCre*; *Nipbl*^Flox/+^ male mice, which make sperm that are either *Nipbl* ^+^ or *Nipbl* ^−^ (see Materials and Methods), and thus produce offspring that are either *Nipbl*-wildtype, or *Nipbl*-haploinsufficient in every tissue. We mated *Stra8-iCre*; *Nipbl*^Flox/+^ male mice with *Isl1^Cre/+^* female mice (in these experiments the *Isl1^Cre/+^* background is being used simply for its *Isl1* haploinsufficiency, as no further Cre-mediated recombination is possible in the offspring of this cross; for simplicity we hereafter refer to the *Isl1^Cre/+^* genotype as *Isl1^+/-^*). This cross generates four possible genotypes WT (*Nipbl^+/+^; Isl1^+/+^*), *Isl1*- haploinsufficient (*Isl1^+/-^*), *Nipbl*-haploinsufficient (*Nipbl^+/-^*), and doubly haploinsufficient (*Nipbl^+/-^*; *Isl1^+/-^*) (Fig. 5A).

**Fig. 5.**
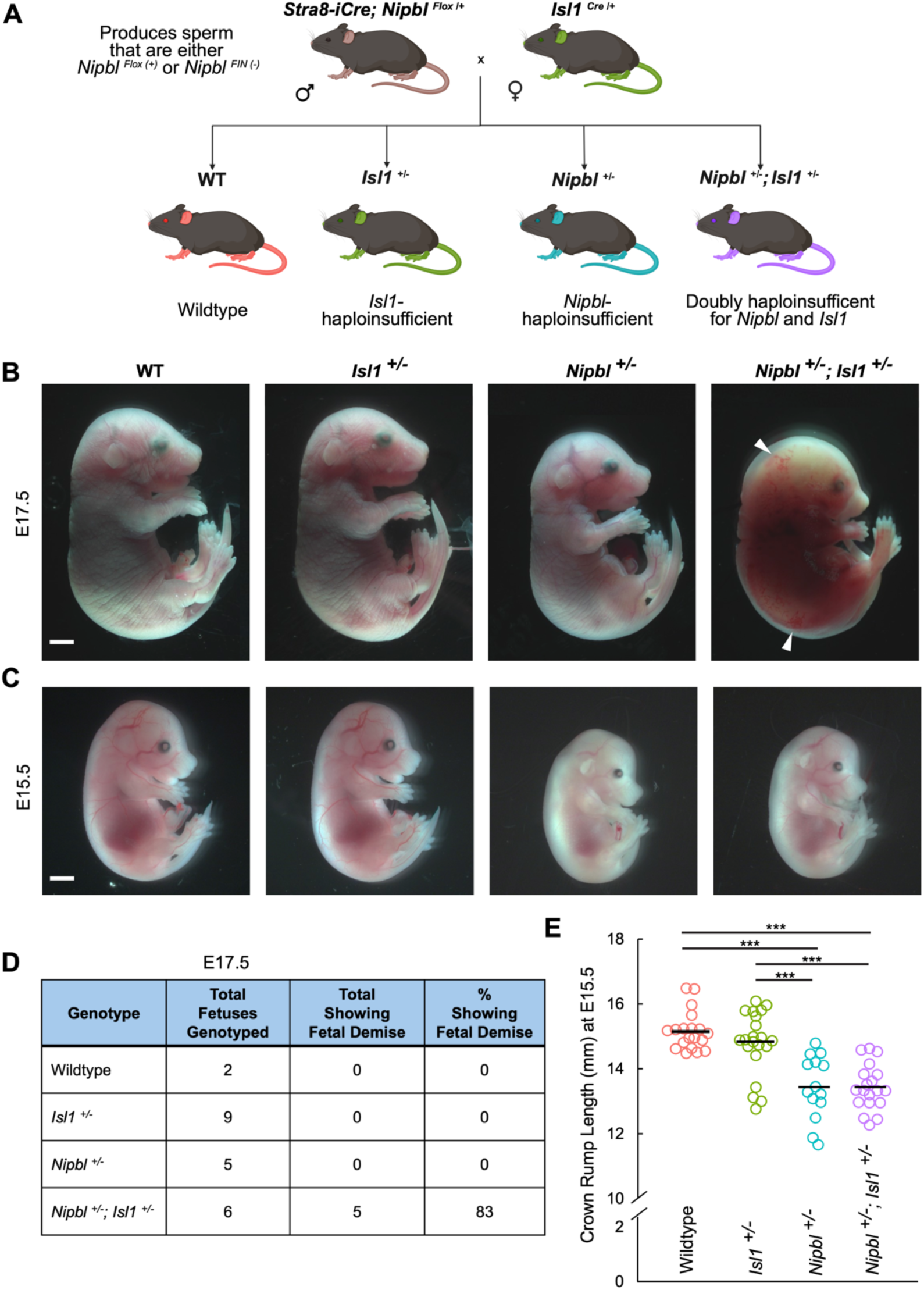
Mice doubly haploinsufficient for *Nipbl* and *Isl1* show fetal demise and reduced body size. (A) *Stra8-iCre*; *Nipbl*^Flox/+^ male mice make sperm that are either *Nipbl* ^+^ or *Nipbl* ^−^ (see Materials and Methods). Mating *Stra8-iCre*; *Nipbl*^Flox/+^ male mice with *Isl1^Cre/+^* female mice produces littermates that are either WT (*Nipbl^+/+^; Isl1^+/+^*), *Isl1*-haploinsufficient (*Isl1^+/-^*), *Nipbl*- haploinsufficient (*Nipbl^+/-^*), or doubly haploinsufficient for *Nipbl* and *Isl1* (*Nipbl^+/-^*; *Isl1^+/-^*). (B) Representative images of E17.5 WT, *Isl1^+/-^*, *Nipbl^+/-^* embryos. The *Nipbl^+/-^*; *Isl1^+/-^* embryo shown is in the early stage of fetal demise, showing hemorrhaging and edema (white arrow heads). Scale bar = 500 uM. (C) Representative images of E15.5 WT, *Isl1^+/-^*, *Nipbl^+/-^*, and *Nipbl^+/-^*; *Isl1^+/-^* embryos. (D) Table of frequency of E17.5 WT (N=2), *Isl1^+/-^* (N=9), *Nipbl^+/-^* (N=5), and *Nipbl^+/-^*; *Isl1^+/-^* (N=6) embryos showing signs of fetal demise. (E) Crown rump length (CRL) of E15.5 WT (N=19), *Isl1^+/-^* (N=21), *Nipbl^+/-^* (N=14), and *Nipbl^+/-^*; *Isl1^+/-^* (N=19) embryos. CRL refers to the length of an embryo from the top of its head (the crown) to the bottom of its torso (the rump). *** represents P ≤ 0.001. P-values from Tukey-Kramer Test.

Upon examining embryos at E17.5, we found, unexpectedly, that 83% of *Nipbl^+/-^*; *Isl1^+/-^* embryos showed signs of fetal demise, notably resorptions and superficial hemorrhage (Fig. 5B, Fig. 5D, Table S13, & Table S15). This prompted us to investigate the phenotype of *Nipbl^+/-^*; *Isl1^+/-^* embryos at an earlier stage, E15.5, at which point, no signs of fetal demise were observed (Fig. 5C & Table S14). Interestingly, we noticed at both E17.5 and E15.5, that both *Nipbl^+/-^* and *Nipbl^+/-^*; *Isl1^+/-^* embryos were smaller than their WT littermates (Fig. 5, B & C). Measurements of crown-rump lengths at E15.5, found that, as in previous studies (Kawauchi et al., 2009, Santos et al., 2016), *Nipbl^+/-^* embryos were 11% smaller than WT embryos (Fig. 5E & Table S16), and that *Nipbl^+/-^*; *Isl1^+/-^* were also 11% smaller than WT embryos. There was no difference between the crown-rump lengths of *Isl1^+/-^* and WT embryos. A decrease in body size of about this magnitude has previously been reported for embryos that are globally *Nipbl^+/-^*, so the data imply that *Isl1* haploinsufficiency does not further alter body size. In contrast, *Isl1* haploinsufficiency, when combined with global *Nipbl* haploinsufficiency, has a strong, negative effect on late embryonic survival.

### *Isl1*-haploinsufficiency enhances the frequency and spectrum of CHDs in *Nipbl*- haploinsufficient mice

Since most *Nipbl^+/-^; Isl1^+/-^* mouse embryos died at E17.5, we looked for CHDs in the hearts of WT, *Nipbl^+/-^*, *Isl1^+/-^* and *Nipbl^+/-^; Isl1^+/-^* mice at E15.5 (Fig. 6A), when no deaths and no signs of fetal demise were observed. Septation of all 4 chambers is also normally complete by E15.5, as is the division of the great vessels (Savolainen et al., 2009). As expected, both *Isl1^+/-^* mice and *Nipbl^+/-^* mice showed CHDs (9.09% and 20.00%, respectively) (Fig. 6B & Table S17). Surprisingly, 66.67% of *Nipbl^+/-^; Isl1^+/-^* mice showed CHDs – far higher than observed in mice haploinsufficient for either *Nipbl* or *Isl1* alone (Fig. 6B). As in *Nipbl^+/-^* mice, *Nipbl^+/-^; Isl1^+/-^* mice also showed ASDs, VSDs, and OTDs (Fig. 6, A & B), with OTDs always occurring in conjunction with either a VSD or both an ASD and VSD. Among the OTDs displayed by *Nipbl^+/-^; Isl1^+/-^* mice were DORVs and PTAs (Fig. 6B), the latter of which were exclusively observed in *Nipbl^+/-^; Isl1^+/-^* mice. Also observed in *Nipbl^+/-^; Isl1^+/-^* mice were CHDs exhibiting a combination of both an ASD and VSD, which occurred in one *Nipbl^+/-^; Isl1^+/-^* mouse (Fig. 6B). These results demonstrate that *Isl1*-haploinsufficiency enhances the frequency and spectrum of CHDs when mice are also *Nipbl*-haploinsufficient in the entire body.

**Fig. 6.**
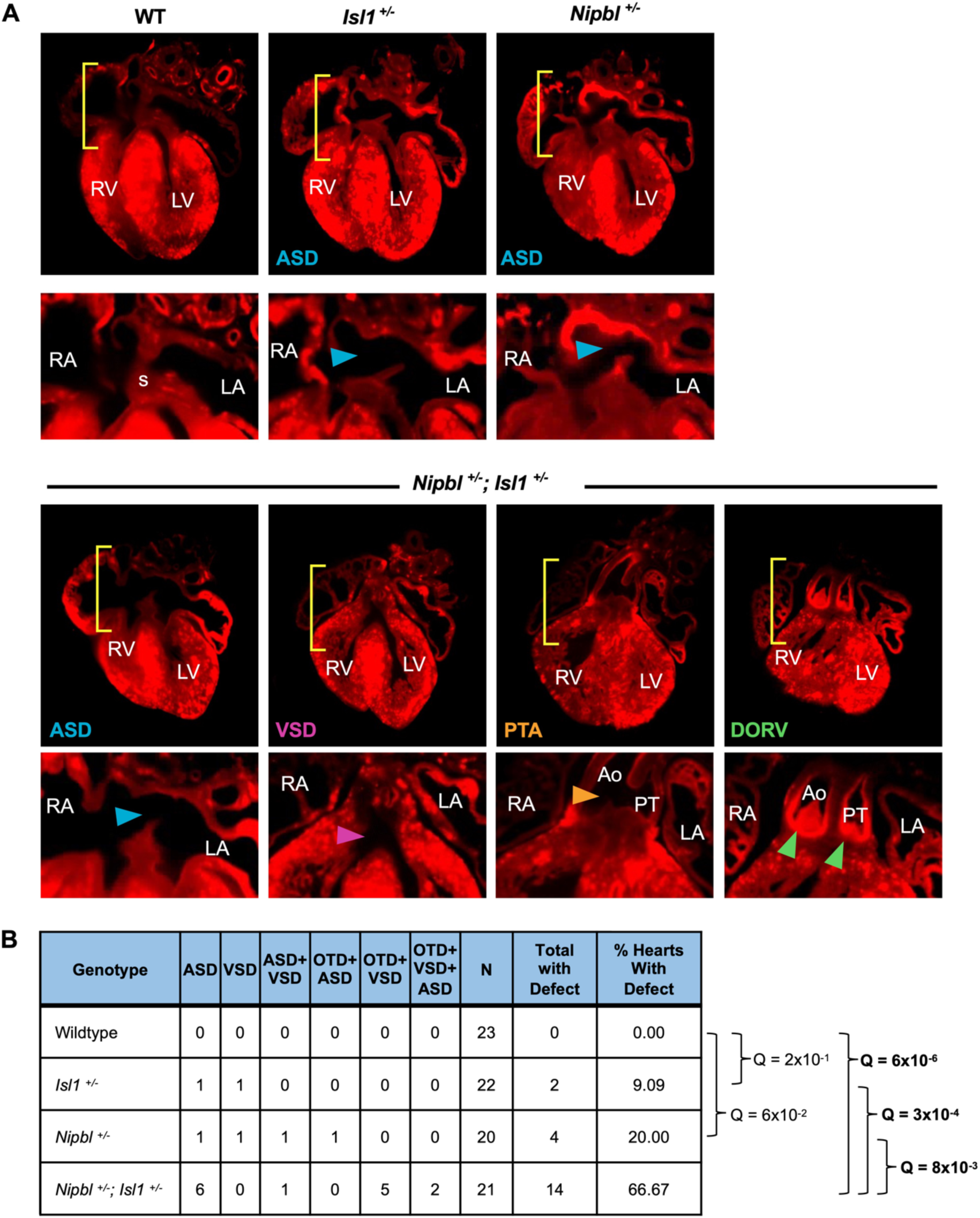
At E15.5, *Isl1*-haploinsufficiency increases the frequency and expands the spectrum of CHDs in *Nipbl*-haploinsufficient mice. (A) LSFM images of anterior view of coronal section of representative E15.5 WT, *Isl1^+/-^, Nipbl^+/-^*, and *Nipbl^+/-^; Isl1^+/-^* hearts. *Isl1^+/-^* and *Nipbl^+/-^* hearts shown show atrial septal defects (ASD, blue arrow). *Nipbl^+/-^; Isl1^+/-^* hearts show atrial septal defects (ASD, blue arrow), ventricular septal defects (VSD, purple arrow), persistent truncus arteriosus (PTA, yellow arrow), and/or double outlet right ventricle (DORV, green arrows). RV, right ventricle; LV, left ventricle; RA, right atrium; LA, left atrium; Ao, aorta; PT, pulmonary tract. (B) Table of frequencies of heart defects observed in WT, *Isl1^+/-^*, *Nipbl^+/-^,* and *Nipbl^+/-^; Isl1^+/-^* E15.5 hearts. Hearts with PTA or DORV were classified as outflow tract defects (OTD). Q-values are from Benjamini-Hochberg corrected Fisher’s Exact Tests comparing genotypes at the ends of each bracket. Q-values less than 0.05 are bolded.

### Gene expression changes preceding CHDs caused by *Nipbl* and *Isl1* haploinsufficiency

As mentioned in the Introduction, one potential mechanism underlying *Nipbl*- haploinsufficiency-mediated gene expression dysregulation is its role in chromatin looping, which facilitates long-range cis-regulatory chromatin interactions (Alonso-Gil and Losada, 2023). *Isl1*, which encodes a LIM-homeodomain (LIM-HD) transcription factor, has also been implicated in chromatin looping and transcriptional regulation (Caputo et al., 2015, Bower and Kvon, 2025). Because *Nipbl^+/-^; Isl1^+/-^* embryos exhibit more frequent and severe heart defects than either single heterozygote, we hypothesize that the increased phenotypic severity may reflect the combined effects of *Nipbl*- and *Isl1*- haploinsufficiency on chromatin looping and gene expression regulation. To investigate this, we performed bulk RNA sequencing on E10.5 hearts from 10 WT, 9 *Nipbl^+/-^*, 9 *Isl1^+/-^,* and 9 *Nipbl^+/-^*; *Isl1^+/-^* mice (Table S18). Principal component analysis (PCA) identified one outlier among *Nipbl^+/-^*samples (Fig. S1), which we excluded from all downstream analysis.

Differential gene expression analysis (DGEA) between *Nipbl^+/-^*, *Isl1^+/-^*, *Nipbl^+/-^*; *Isl1^+/-^* and WT hearts revealed that *Isl1^+/-^* hearts misexpressed 51 genes: 40 were overexpressed, while 11 were underexpressed (Fig. 7A). Since 44 of these genes exhibited small changes in expression (less than 2-fold change) (Fig. 7A), we performed gene set enrichment analysis (GSEA) to determine whether they were enriched or de-enriched for any Hallmark gene sets representing well-curated biological processes (Liberzon et al., 2015). Our analysis found that *Isl1^+/-^*hearts were enriched for 8 Hallmark gene sets (Fig. 7B), including the MTORC1 signaling gene set, whose genes regulate cardiomyocyte growth and proliferation (Sciarretta et al., 2018). Conversely, 7 gene sets were de-enriched, including the epithelial mesenchymal transition gene set, whose genes are involved in the formation of the cardiac endocardial cushions that give rise to the heart’s septa and valves (von Gise and Pu, 2012). Manual curation of overexpressed genes in *Isl1^+/-^* hearts identified three groups of genes (Fig. 7C): 1) genes encoding SH3 domain-containing proteins, which help maintain the structural integrity of the cardiomyocyte cytoskeleton (McLendon et al., 2022); 2) transcription factors, including *Ssh3*, *Zfp473*, and *Foxo6* – the latter of which has been implicated in cardiac pathological remodeling and dysfunction (Zhang et al., 2023); and 3) genes involved in anterior posterior specification, such as *Heyl* and two Hox genes (*Hoxa3* & *Hoxa1*) *(Leimeister et al., 2000, Hubert and Wellik, 2023)*. In contrast, *Pax6*, another gene involved in anterior-posterior specification, (Takamiya et al., 2020), was underexpressed in *Isl1^+/-^* hearts (Fig. 7C).

**Fig. 7.**
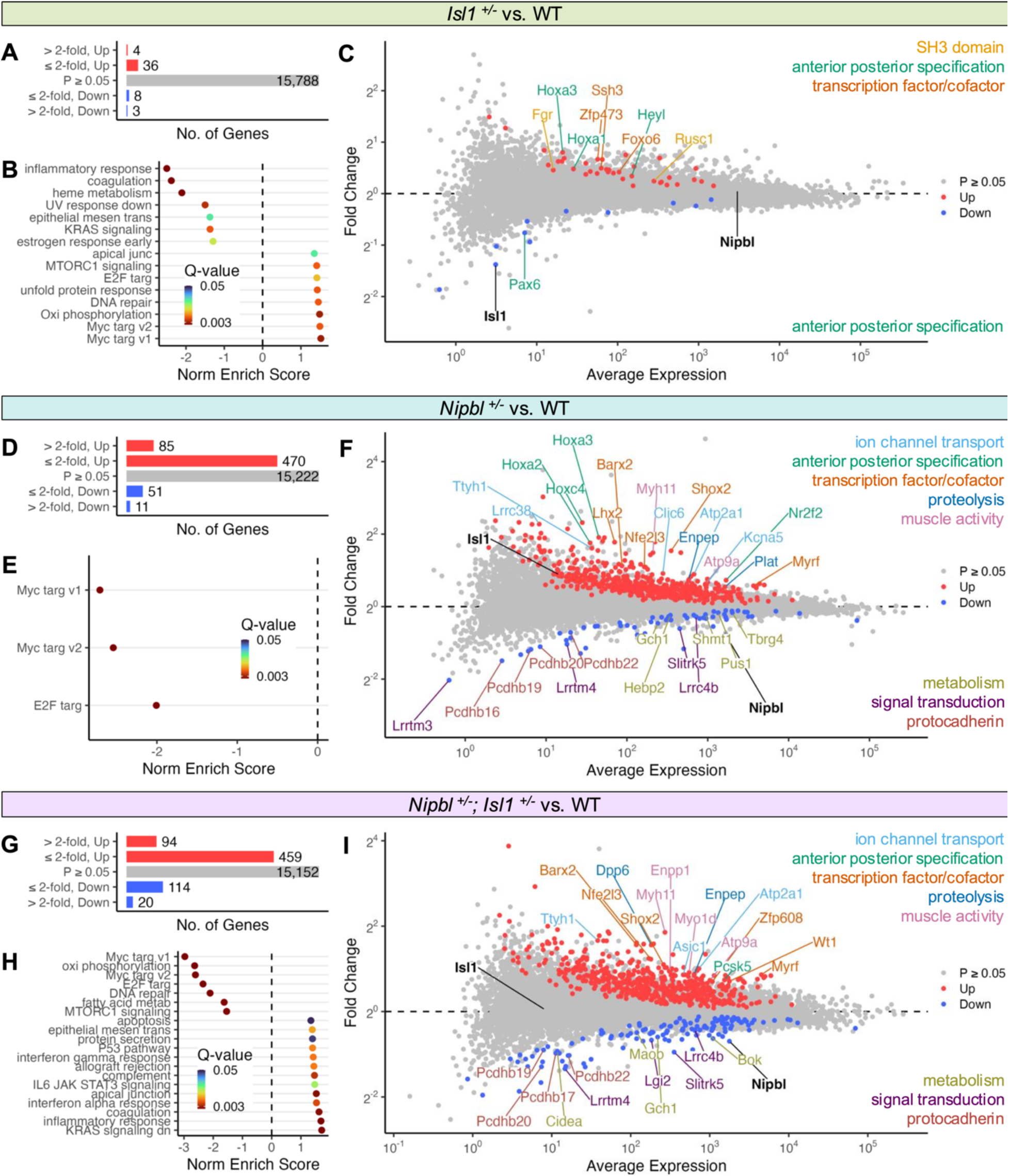
Gene expression changes in E10.5 hearts in *Isl1^+/-^, Nipbl^+/-^,* and *Nipbl^+/-^; Isl1^+/-^* mice. Number of genes differentially expressed in (A) *Isl1^+/-^,* (D) *Nipbl^+/-^,* or (G) *Nipbl^+/-^; Isl1^+/-^* E10.5 hearts relative to WT. Differentially expressed genes are binned according to whether they were differentially expressed less than or more than two-fold up or down. Differential gene expression analysis was performed using the ANOVA and Tukey-Kramer tests. Genes with Benjamini- Hochberg corrected P-values less than alpha 0.05 were considered differentially expressed. Hallmark gene sets enriched or de-enriched in their expression in (B) *Isl1^+/-^,* (E) *Nipbl^+/-^,* or (H) *Nipbl^+/-^; Isl1^+/-^* E10.5 hearts relative to WT. Gene set enrichment analysis was performed using the R package called FGSEA. Only statistically significant gene sets are shown (Q < 0.05). Fold change in expression of genes in (C) *Isl1^+/-^,* (F) *Nipbl^+/-^,* or (I) *Nipbl^+/-^; Isl1^+/-^* E10.5 hearts from that of WT hearts. The position of a gene along the x-axis correspond to its average expression in WT hearts. Genes are colored according to whether they were expressed at WT levels (gray, Q ≥ 0.05), overexpressed (red, Up), or underexpressed (blue, Down).

*Nipbl^+/-^* hearts misexpressed 617 genes, with 555 overexpressed and 62 underexpressed (Fig. 7D). Consistent with previous studies showing that *Nipbl*-haploinsufficiency results in hundreds of small (less than two-fold) gene expression changes (Santos et al., 2016, Kawauchi et al., 2009), 84% of the differentially expressed genes in *Nipbl^+/-^*hearts showed less than two-fold change. GSEA revealed de-enrichment for three gene sets: two Myc targets gene sets and one E2F targets gene set (Fig. 7E), whose genes are associated with cell growth/proliferation and cell cycle regulation, respectively (von Harsdorf et al., 1999, Bywater et al., 2020). *Nipbl^+/-^* hearts did not show significant enrichment for any Hallmark gene set. Manual curation of overexpressed genes in *Nipbl^+/-^* hearts identified 5 groups of genes (Fig. 7F): 1) ion channel transportation genes (*Ttyh1, Lrrc38, Clic6, Atpa21*, & *Kcna5*) (Suzuki, 2006, Gonzalez-Perez et al., 2022); 2) genes associated with anterior posterior specification, including *Nr2f2* (Alzu’bi et al., 2017) and three *Hox* genes (*Hoxa2, Hoxa3,* & *Hoxc4*) (Hubert and Wellik, 2023); 3) proteolysis-related genes (*Enpep* & *Plat*); 4) genes involved in muscle contraction and activity (*Myh11* & *Atp9a*) (Sweeney et al., 2020); and 5) transcription factors, including *Nfe2l3, Myrf*, *Barx2, Shox2, and Lhx2*. Interestingly, *Shox2* is involved in the formation of the heart’s sinoatrial node (Hu et al., 2018), and *Lhx2* is expressed in the pharyngeal mesoderm, which contributes to the heart’s outflow tract (Harel et al., 2012). In contrast, three groups of genes showed downregulation: 1) protocadherins (*Pcdhb16, Pcdhb19, Pcdhb20*, & *Pcdhb22*), which have also been found to be downregulated in other *Nipbl^+/-^* tissues (Kawauchi et al., 2009); 2) genes involved in metabolism (*Gch1, Shmt1, Tbrg4, Pus1*, & *Hebp2*); and 3) genes associated with signal transduction (*Cidea, Maob, Gch1,* & *Bok*).

*Nipbl^+/-^*; *Isl1^+/-^* (doubly haploinsufficient) hearts exhibited misexpression of 687 genes, with 553 overexpressed and 134 underexpressed (Fig. 7G). GSEA identified 13 enriched Hallmark gene sets, including the epithelial mesenchymal transition gene set, which we mentioned earlier, is involved in the formation of heart septa and valves (Fig. 7H). In contrast, seven gene sets were de-enriched, including two Myc targets gene sets and one E2F targets gene set, all three of which also showed de-enrichment in *Nipbl^+/-^; Isl1^+/-^* hearts (Fig. 7H). Manual curation of overexpressed genes in *Nipbl^+/-^*; *Isl1^+/-^*hearts found substantial overlap with the transcriptional changes observed in *Nipbl^+/-^* hearts. As in *Nipbl^+/-^* hearts, *Nipbl^+/-^; Isl1^+/-^* hearts showed overexpression of genes involved in ion channel transportation anterior-posterior specification, proteolysis, muscle contraction and activity, and transcription factors such as *Barx2, Shox2,* and *Nfe2l3*. Similarly, genes downregulated in *Nipbl^+/-^; Isl1^+/-^*hearts mirrored many of those in *Nipbl^+/-^* hearts, including protocadherins, genes involved in metabolism, and genes involved in signal transduction.

### Gene expression in *Nipbl^+/-^; Isl1^+/-^* hearts most closely resemble *Nipbl^+/-^* hearts

In Fig. 7, we noticed that *Nipbl^+/-^; Isl1^+/-^* hearts exhibited gene expression changes that closely resembled those in *Nipbl^+/-^* hearts, whereas *Isl1^+/-^* hearts showed minimal overlap with either *Nipbl^+/-^; Isl1^+/-^* or *Nipbl^+/-^*hearts. To quantitatively assess transcriptional similarity, we performed principal component analysis (PCA) on hearts of all genotypes. The first 2 components accounted for 35% and 25% of the total variance, respectively (Fig. 8A). Plotting the hearts along these components revealed a genotype-based separation along Component 1 into two groups (Fig. 8B): *Nipbl^+/-^; Isl1^+/-^* hearts clustered with *Nipbl^+/-^* hearts, while *Isl1^+/-^* hearts clustered with WT hearts. To further quantify the overlap in transcriptional profiles, we next compared the DEGs shared among *Isl1^+/-^*, *Nipbl^+/-^*, and *Nipbl^+/-^; Isl1^+/-^* hearts. We defined DEGS as shared when they were differentially expressed in the same direction (either overexpressed or underexpressed relative to WT). Of the 761 DEGs identified in *Nipbl^+/-^*or *Nipbl^+/-^; Isl1^+/-^* hearts, 543 (∼71%) were shared between the two genotypes (Fig. 8C). When categorized by expression direction, 59 of the 137 downregulated genes (∼43%) and 494 of the 634 upregulated genes (∼78%) were shared (Fig. 8D). In contrast, DEGs in *Isl1^+/-^*hearts overlapped with only ∼6% of the DEGs identified in either *Nipbl^+/-^*or *Nipbl^+/-^; Isl1^+/-^* hearts (Fig. 8C). Together, these results suggest that *Isl1* influences gene expression through mechanisms separate from those exploited by *Nipbl*.

**Fig. 8.**
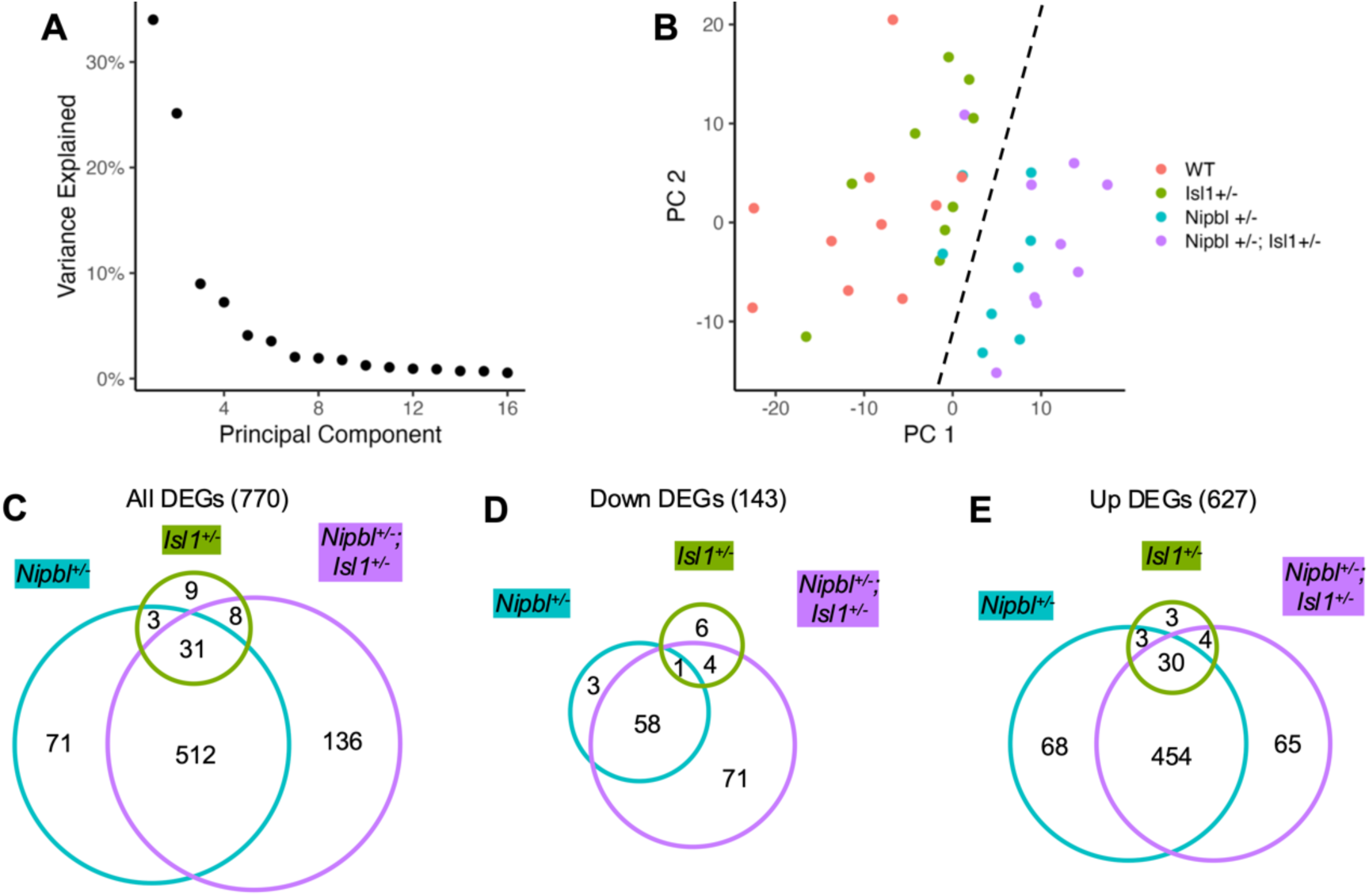
Comparison of differentially expressed genes among *Isl1^+/-^, Nipbl^+/-^,* and *Nipbl^+/-^; Isl1^+/-^* hearts. (A) Variance explained by first 16 principal components among WT, *Isl1^+/-^, Nipbl^+/-^,* and *Nipbl^+/-^; Isl1^+/-^* hearts. (B) Position of WT, *Isl1^+/-^, Nipbl^+/-^,* and *Nipbl^+/-^; Isl1^+/-^* hearts along Principal Components 1-2. *Isl1^+/-^* hearts separated along Component 1 with WT hearts (left of dashed line). *Nipbl^+/-^; Isl1^+/-^* hearts separated along Component 1 with *Nipbl^+/-^* hearts (right of dashed line). Number of genes (C) differentially expressed, (D) downregulated, or upregulated (E) in E10.5 *Isl1^+/-^, Nipbl^+/-^,* and *Nipbl^+/-^; Isl1^+/-^* hearts relative to WT hearts that are shared and/or unique to each genotype. DEGs were considered shared when they were differentially expressed in the same direction (overexpressed or underexpressed relative to WT).

### Effects of *Nipbl*- and *Isl1*-haploinsufficiency on gene expression are primarily additive

How can *Nipbl^+/-^; Isl1^+/-^* hearts share so many of the same DEGs as *Nipbl^+/-^* hearts, yet exhibit a substantially higher frequency and broader spectrum of heart defects (67% versus 20%, respectively) (Fig. 6)? One possible explanation is that the gene expression changes caused by *Nipbl*-haploinsufficiency interact epistatically with those caused by *Isl1*-haploinsufficiency in *Nipbl^+/-^; Isl1^+/-^* hearts, resulting in combined gene expression effects that deviate from the sum of their individual contributions. That epistatic effects may be present in *Nipbl^+/-^; Isl1^+/-^* hearts is also suggested by the frequency of defects observed at E15.5 – 67% – which is more than twice the expected frequency if the combined effects of *Nipbl*- and *Isl1*-haploinsufficiency on heart defect frequency were merely additive (20% in *Nipbl^+/-^* hearts and 9% in *Isl1^+/-^* hearts) (Fig. 6).

To identify epistatic gene expression effects in *Nipbl^+/-^; Isl1^+/-^* hearts, we used linear regression to calculate, for each gene, its expected additive effect and any observed deviation from this expectation. A T-test was used to determine which deviations were statistically significant. To account for multiple testing, we applied the Benjamini-Hochberg correction to the 770 genes identified earlier in Fig. 8C as being differentially expressed (relative to WT) among *Isl1^+/-^*, *Nipbl^+/-^*; *Nipbl^+/-^; Isl1^+/-^* hearts. Only three genes had Q-values < 0.05: *Lrrc38, Zfp846,* and *Clstn2*. To classify the nature of their epistatic interactions, we plotted the expected fold changes in expression against the observed fold changes in *Nipbl^+/-^; Isl1^+/-^*hearts (Fig. 9A). This revealed that *Lrrc38* and *Zfp846* exhibited negative epistasis, being less than downregulated than expected. *Clstn2* displayed sign epistasis, being downregulated when upregulation was expected. Among these, *Lrrc38* encodes a protein involved ion channel transport, which may influence muscle contraction (Ehrlich et al., 2020). Otherwise, 767 out of the 770 DEGs (99.6%) followed the expected additive model in *Nipbl^+/-^; Isl1^+/-^* hearts. Thus, the synergistic phenotypic effects observed in *Nipbl^+/-^; Isl1^+/-^* hearts appears to result primarily from additive gene expression effects rather than widespread epistatic interactions.

**Fig. 9.**
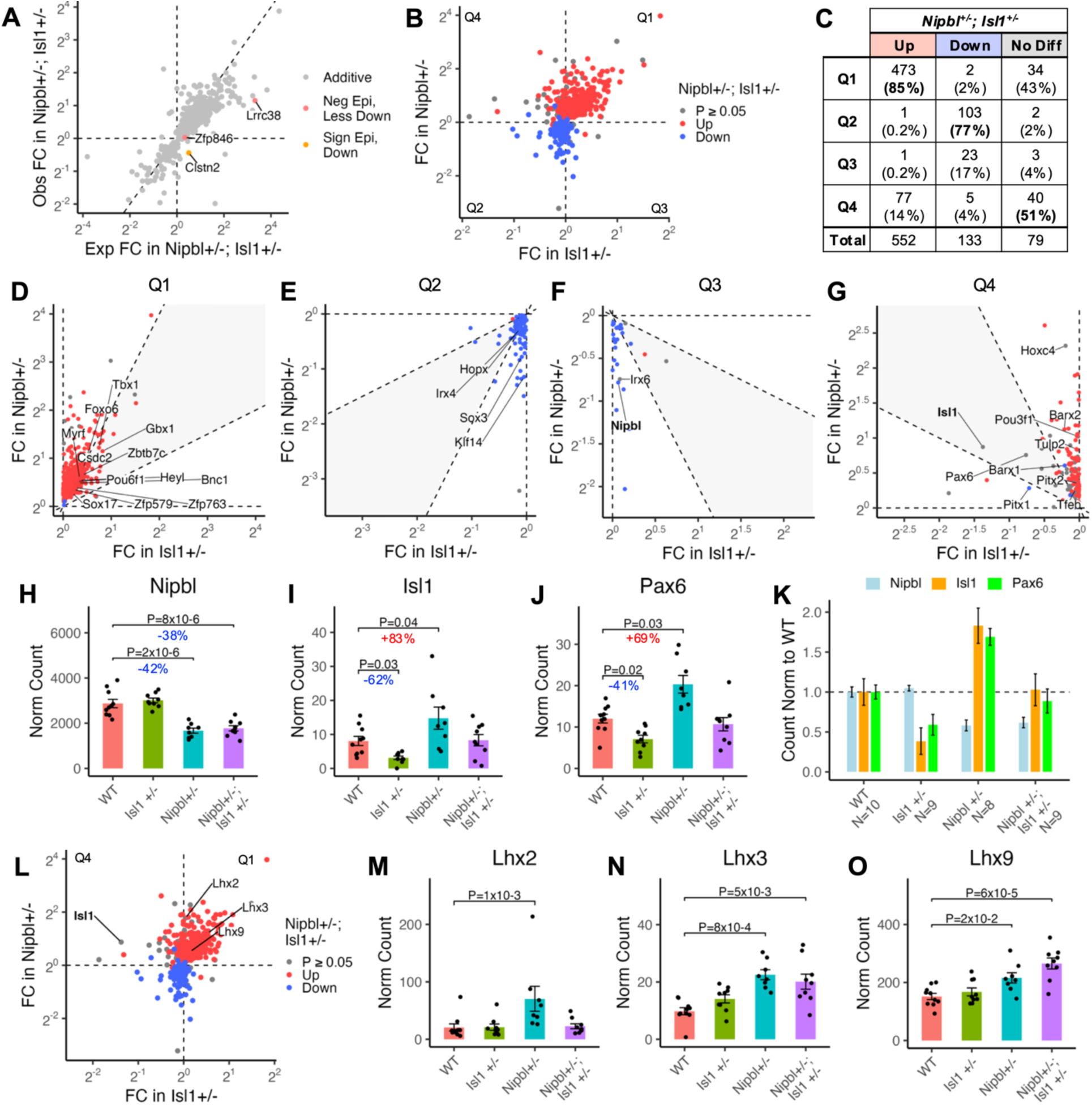
Epistatic and additive gene expression effects in *Nipbl^+/-^; Isl1^+/-^* hearts. (A) Observed vs. expected fold changes the expression differentially expressed genes (from Fig. 8C) in *Nipbl^+/-^; Isl1^+/-^*hearts, assuming purely additive effects of *Isl1-* and *Nipbl*-haploinsufficiency (see Materials and Methods). Genes are color-coded based on their classification as additive, negatively epistatic (less downregulated than expected), or showing sign epistasis (downregulated when upregulation was expected). (B) Fold changes in the expression of additive genes (from Panel A) in *Isl1^+/-^* hearts vs *Nipbl^+/-^* hearts. Genes are colored based on their differential expression status in *Nipbl^+/-^; Isl1^+/-^*hearts: significantly upregulated (red), downregulated (blue), or unchanged. (C) Number of genes that were upregulated, downregulated, or showing no difference in *Nipbl^+/-^; Isl1^+/-^* hearts within each quadrant of Panel B. (D-G) Fold changes in the expression of additive genes from each quadrant of Panel B: (D) Quadrant 1, (E) Quadrant 2, (F) Quadrant 3, and (G) Quadrant 4. Gray areas highlight genes whose fold changes reflect strong contributions from both *Nipbl*- and *Isl1*-haploinsufficiency. Labeled genes are transcription factors. (H-J) Expression of (H) *Nipbl,* (I) *Isl1,* and (J) *Pax6* expression in E10.5 WT, *Nipbl^+/-^, Isl1^+/-^,* and *Nipbl^+/-^; Isl1^+/-^* hearts. Error bars show standard error of the mean. P-values from Tukey-Kramer test. Read counts for *Isl1* reflect those overlapping the Exon 1-Exon 2 junction (see Materials and Methods). (K) Normalized expression of *Nipbl, Isl1,* and *Pax6* in E10.5 hearts of each genotype, relative to WT hearts. (L) Same as Panel B. All LIM-HD proteins are labeled. H) *Lhx2,* (I) *Lhx3,* and (J) *Lhx9* expression in E10.5 WT, *Nipbl^+/-^, Isl1^+/-^,* and *Nipbl^+/-^; Isl1^+/-^* hearts. Error bars show standard error of the mean. P-values from Tukey-Kramer test.

Next, we looked at the 767 genes classified as exhibiting additive effects in *Nipbl^+/-^; Isl1^+/-^* hearts. In Fig. 9B, we colored each gene based on whether it was significantly upregulated, downregulated, or unchanged in *Nipbl^+/-^; Isl1^+/-^* hearts and plotted their fold changes in *Isl1^+/-^* hearts against their fold changes in *Nipbl^+/-^* hearts. For each category (upregulated, downregulated, or no difference), we quantified the number of genes that fell into the four quadrants of the plot (Fig. 9C). Among the genes upregulated in *Nipbl^+/-^; Isl1^+/-^*hearts, 85% reflected the additive effects of being upregulated in both *Nipbl^+/-^*and Isl1+/- hearts (Q1). Similarly, 77% of the genes downregulated in *Nipbl^+/-^; Isl1^+/-^* hearts were downregulated in both *Nipbl^+/-^*and *Isl1^+/-^* hearts (Q2). Among genes showing no significant change in *Nipbl^+/-^; Isl1^+/-^* hearts, 43% were upregulated in both *Nipbl^+/-^*and *Isl1^+/-^* hearts (Q1). In contrast, the largest proportion (51%) of unchanged genes are the result being downregulated in *Isl1^+/-^*hearts and upregulated in *Nipbl^+/-^* hearts (Q4). We refer to these as “additive but opposing effects”, where *Nipbl*- and *Isl1*-haploinsufficiency independently alter expression in opposite directions, resulting in minimal net change when combined. Interestingly, only 4% of unchanged genes are the result of being upregulated in *Isl1^+/-^* hearts and downregulated in *Nipbl^+/-^*hearts (Q3). These results suggest that the increased frequency of heart defects in *Nipbl^+/-^; Isl1^+/-^* hearts may arise not only from additive gene expression changes that occur in the same direction, but also from opposing additive effects, particularly where upregulation in *Nipbl^+/-^* hearts and downregulation in *Isl1^+/-^* hearts cancel each other out.

### *Isl1*- and *Nipbl*-haploinsufficiency have opposing effects on the expression of *Pax6* and *Isl1*

To better understand the additive effects in *Nipbl^+/-^; Isl1^+/-^* hearts, we next examined the genes distributed across each quadrant, beginning with Q1, which contains genes that are upregulated in both *Isl1^+/-^* and *Nipbl^+/-^* hearts. To focus on genes whose expression is strongly influenced by both *Nipbl*- and *Isl1*-haploinsufficiency, rather than predominantly by one alone, we concentrated on genes located far from either the x- or y-axis, highlighted in gray (Fig. 9D). Within this region, we identified 12 transcription factors, which were additively upregulated in *Isl1^+/-^*, *Nipbl^+/-^*, *Nipbl^+/-^; Isl1^+/-^* hearts (Fig. 9D). Of these, 5 have been previously implicated in heart development or congenital heart defects: *Foxo6, Heyl, Tbx1, Bnc1*, and *Sox17* (Fig. S2, A to E) (Zhang et al., 2023, Fischer et al., 2007). For example, *Tbx1* plays a role in the development of the outflow tract (Xu et al., 2004), *Bnc1* is involved in the formation of the epicardium (Gambardella et al., 2019), and *Sox17* regulates the development of the endocardium (Saba et al., 2019). In Q2, *Irx4*, which marks the earliest cardiac progenitor populations destined to form the ventricular myocardium (Nelson et al., 2014) was additively downregulated in *Isl1^+/-^*, *Nipbl^+/-^*, *Nipbl^+/-^; Isl1^+/-^* hearts (Fig. 9E & Fig. S2F).

In Q3, no transcription factors were identified in the gray area. The only transcription factor in the entire quadrant was *Irx6* (Fig. 9F), which was more strongly downregulated in *Nipbl^+/-^* hearts than upregulated in *Isl1^+/-^* hearts and showed no significant difference in *Nipbl^+/-^; Isl1^+/-^* hearts (Fig. S2G). Like *Sox17*, there is evidence that *Irx6* plays a role in the development of the endocardium (Kim et al., 2012). *Nipbl* itself was also located in this quadrant, showing downregulation in *Nipbl^+/-^*and *Nipbl^+/-^; Isl1^+/-^* hearts, but no difference in *Isl1^+/-^* hearts (Fig. 9H). In Q4, transcription factors located outside the gray area included *Barx1, Barx2, Pitx1, Pitx2*, and *Hoxc4* (Fig. S2, H to L). Of these, *Pitx2* and *Hoxc4* showed no difference in expression in *Nipbl^+/-^; Isl1^+/-^* hearts (Fig. S2, K & L). Interestingly, both *Hoxc4* and *Pitx2* play roles in axial patterning, with *Hoxc4* regulating anterior-posterior patterning (Lescroart and Zaffran, 2018) and *Pitx2* contributing to left-right asymmetry (Franco et al., 2017). *Pax6* and *Isl1* also showed no difference in *Nipbl^+/-^; Isl1^+/-^* hearts and were both located in the gray area of Q4 (Fig. 9G). Interestingly, *Pax6* is a transcriptional target of *Isl1* (Gao et al., 2019) and *Pax6* has been implicated as an inhibitor of cardiac fibroblast differentiation (Feng et al., 2020).

That *Isl1* was expressed at WT levels in *Nipbl^+/-^; Isl1^+/-^* hearts was particularly surprising, since we fully expected it to be downregulated –given that *Nipbl^+/-^; Isl1^+/-^* hearts are haploinsufficient for *Isl1*, and *Isl1* is significantly downregulated in *Isl1^+/-^* hearts (by 62% relative to WT) (Fig. 9H). However, as shown in Fig. 9G, *Nipbl-* and *Isl1*-haploinsufficiency exert opposing effects on the expression of *Isl1*: *Isl1* expression is upregulated in *Nipbl^+/-^*hearts by 83% (relative to WT), whereas it is downregulated in *Isl1^+/-^* hearts (Fig. 9I). In *Nipbl^+/-^; Isl1^+/-^* hearts, downregulation of Isl1 caused by *Isl1*-haploinsufficiency is counteracted by upregulation driven by *Nipbl*-haploinsufficiency, effectively restoring *Isl1* expression to WT levels (Fig. 9K). A similar phenomenon is also observed for *Pax6*. It is significantly downregulated in *Isl1^+/-^* hearts (by 41%) and significantly upregulated in *Nipbl^+/-^* hearts (by 69%) (Fig. 9J) but expressed at WT levels in *Nipbl^+/-^; Isl1^+/-^*hearts. These results highlight how complex interactions between two global chromatin organizers (*Nipbl* & *Isl1*) can mask underlying genetic vulnerabilities (by returning expression levels to WT) that otherwise may potentially contribute to the phenotypic severity of congenital defects.

### *Isl1*- and *Nipbl*-haploinsufficiency upregulate other *Lhx* LIM-homeodomain proteins

As mentioned earlier, *Isl1* belongs to the LIM-HD family of transcription factors, which share conserved structural motifs and have been implicated in chromatin looping and gene expression regulation (Bower and Kvon, 2025). Given these shared features, we next asked whether other LIM-HD family members, specifically *Lhx* genes, show similar expression changes to *Isl1* in *Isl1^+/-^*, *Nipbl^+/-^*, and *Nipbl^+/-^; Isl1^+/-^* hearts. To investigate this, we took Fig. 9B and labeled all *Lhx* proteins (Fig. 9L). Three *Lhx* genes (*Lhx2*, *Lhx3*, and *Lhx9*) were identified, and all mapped to Q1, indicating upregulation in both *Isl1^+/-^* and *Nipbl^+/-^* hearts. However, only the upregulation in *Nipbl^+/-^* hearts were statistically significant (Fig. 9, M to O). In *Nipbl^+/-^; Isl1^+/-^* hearts, *Lhx2* expression was consistent with additive expectations, but was not significantly different from WT (Fig. 9M). In contrast, both *Lhx3* and *Lhx9* were additively further upregulated in *Nipbl^+/-^; Isl1^+/-^* hearts (Fig. 9, N & O). These results suggest that *Isl1*- and *Nipbl*-haploinsufficiency have additive, upregulatory effects on the *Lhx* members of LIM-HD genes, rather than the additive but opposing effects observed for *Isl1* itself.

## DISCUSSION

### How can additive gene expression changes produce synergistic CHD phenotypes?

In this study, we found that *Nipbl^+/-^; Isl1^+/-^*hearts exhibited a broader spectrum and a synergistically higher frequency of CHDs compared to either *Nipbl^+/-^* or *Isl1^+/-^* hearts alone (Fig. 6). Yet, transcriptional profiling revealed that *Nipbl^+/-^; Isl1^+/-^*hearts closely resembled *Nipbl^+/-^* hearts (Figs. 7 & 8), and identified only three genes (*Lrrc38, Zfp846,* & *Clstn2*) – none previously implicated in heart development – that significantly deviated from additive expectations (Fig. 9A). Since 99.6% of the gene expression changes were additive, the synergistic phenotype likely arises from the cumulative impact of many additive changes, rather than a few epistatic ones.

Most of the additive effects reflected gene expression changes in the same direction – either jointly up- or downregulated relative to WT (Fig. 9C). For example, the transcription factor *Irx4* was more strongly downregulated, while *Foxo6, Heyl, Tbx1, Bnc1*, and *Sox17* were more strongly upregulated, in *Nipbl^+/-^; Isl1^+/-^* hearts than in either single heterozygote (Fig. 9, D & E). However, because *Nipbl^+/-^; Isl1^+/-^* hearts clustered closely with *Nipbl^+/-^* hearts in PC space (Fig. 8), the same direction differences observed in gene expression between these two genotypes are modest.

Are *opposing* regulatory factors the source of the synergistic CHD burden observed in *Nipbl^+/-^; Isl1^+/-^* hearts? Approximately 19% of additive effects arose from opposing gene expression differences between *Nipbl*- and *Isl1*-haploinsufficiency, which partially or fully canceled each other out (Fig. 9C). One such example was the transcription factor *Irx6*, which was downregulated in *Nipbl^+/-^* hearts, upregulated in *Isl1^+/-^* hearts, but expressed at WT levels in *Nipbl^+/-^; Isl1^+/-^* hearts (Fig. 9F). Most commonly, we found that such genes were *upregulated* in *Nipbl^+/-^* hearts but *downregulated* in *Isl1^+/-^* hearts (Fig. 9C). Strikingly, the transcription factors *Hoxc4, Pitx2, Isl1* itself, and *Pax6* (a known target of *Isl1*), all returned to WT expression levels in the double heterozygote (Fig. 9, G to K; Fig. S2).

*Isl1* expression is significantly upregulated in *Nipbl^+/-^*hearts, but returns to WT levels in *Nipbl^+/-^; Isl1^+/-^*hearts, indicating that *Isl1* upregulation is lost. Since *Nipbl^+/-^; Isl1^+/-^* hearts exhibited a markedly higher burden of CHDs compared to *Nipbl^+/-^* hearts (Fig. 6), these findings suggest that loss of *Isl1* upregulation is critical to this higher burden of CHDs. We propose that in *Nipbl^+/-^* hearts, *Isl1* upregulation functions as a compensatory mechanism that partially mitigates CHD risk. When *Isl1* dosage is reduced, this compensatory effect is lost, resulting in a substantial increase in CHD frequency and severity, as well as increased fetal demise (Fig. 6).

What might *Isl1* upregulation be compensating for, in response to *Nipbl*- haploinsufficiency? One possibility stems from the shared molecular functions of *Nipbl* and *Isl1*. Both have been shown to mediate long-range chromatin interactions between enhancers and promoters (Alonso-Gil and Losada, 2023, Bower and Kvon, 2025, Muto et al., 2014). It therefore seems likely that *Isl1* upregulation compensates for pathogenic changes in chromatin interactions caused by *Nipbl*-haploinsufficiency. A proposed model for this is shown in Fig. 10.

**Fig. 10.**
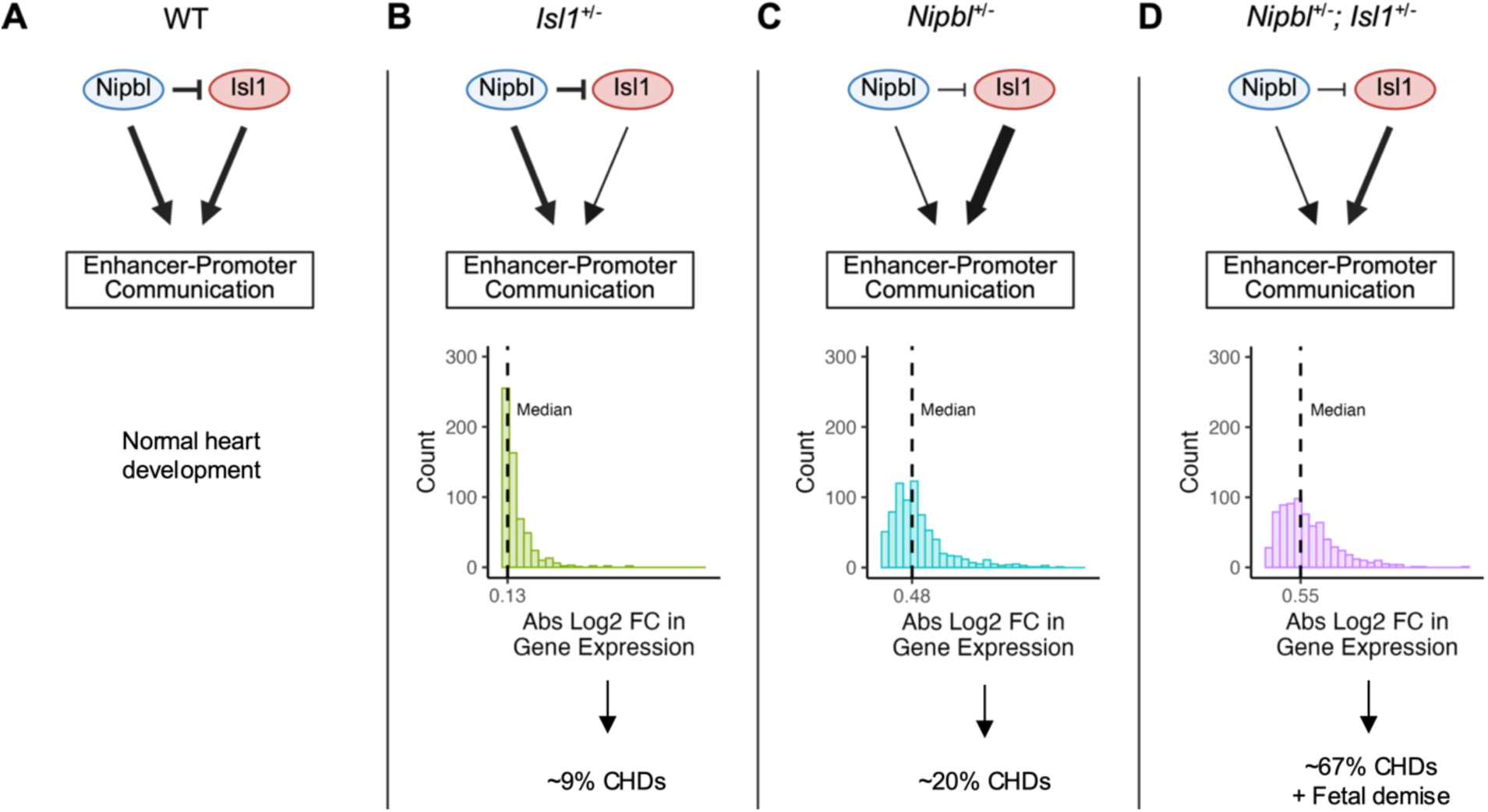
Proposed model of how *Nipbl* and *Isl1* dosage influence enhancer-promoter communication and CHD risk. (A) WT: *Nipbl* and *Isl1* are both expressed at normal levels and contribute independently to the formation of chromatin loops that facilitate enhancer-promoter interactions required for proper heart development. (B) *Isl1*-haploinsufficiency: Reduced *Isl1* dosage leads to the loss of a subset of *Isl1*-mediated enhancer-promoter interactions. Since *Nipbl* expression remains normal, most chromatin loops are preserved. This results in a modest median absolute log2 fold change in expression (0.13) and a low incidence of CHDs (∼9%). (C) *Nipbl*- haploinsufficiency: Lower *Nipbl* levels reduce *Nipbl*-mediated chromatin looping, disrupting some enhancer-promoter contacts. However, *Isl1* is transcriptionally upregulated in response, and this upregulation may partially compensate by preserving a subset of chromatin interactions. The resulting gene expression changes are more extensive (median absolute log2 fold change = 0.48), and CHD incidence increases (∼20%). (D) *Nipbl*- and *Isl1*-haploinsufficiency: *Nipbl*- mediated chromatin loops are lost due to Nipbl haploinsufficiency. Although *Isl1* is typically upregulated in this context, haploinsufficiency constrains *Isl1* expression to WT levels, preventing compensatory maintenance of chromatin interactions normally mediated by *Nipbl*. The combined loss of *Nipbl* function and *Isl1*-mediated compensation results in an additive increase in the median absolute log2 fold change in gene expression (0.55) and a dramatic increase in CHDs (∼67%) and fetal demise.

That *Nipbl* and *Isl1* functions converge on developmental programs involved in cardiac morphogenesis is supported by phenotypic evidence in the present study. ASDs were the most common defect observed in both *Isl1⁺^/^⁻* and *Nipbl^+/-^* hearts (Figs. 1, 4, & 6), but their incidence rose dramatically when *Isl1*- and *Nipbl*-haploinsufficiency were combined (Fig. 6): ASD occurrence was 5% in *Isl1⁺^/^⁻* hearts, and 15% in *Nipbl^+/-^*hearts, but 43% in *Nipbl^+/-^; Isl1^+/-^* hearts. Moreover, the majority (64%) of *Nipbl^+/-^; Isl1^+/-^* hearts with a CHD exhibited an ASD, even when other defects were present (Fig. 6). These observations suggest that atrial septal morphogenesis is particularly sensitive to the functional convergence of *Nipbl* and *Isl1*.

While *Isl1* upregulation buffers against defects in *Nipbl^+/-^*hearts, this compensation is clearly incomplete. Even with elevated *Isl1* expression, ∼37% of *Nipbl^+/-^* hearts still exhibit CHDs (Fig. 1). This suggests that *Nipbl* regulates a broader repertoire of enhancer-promoter communications than *Isl1* can compensate for. Consistent with this, *Isl1^+/-^* hearts exhibited only 51 differentially expressed genes (Fig. 7A) and had the lowest CHD frequency (9%) among the three mutant genotypes. Additionally, *Isl1^+/-^* hearts more closely resembled WT than *Nipbl^+/-^*hearts in PC space (Fig. 8), reinforcing the idea that *Isl1* regulates only a subset of *Nipbl*- dependent chromatin interactions.

Interestingly, other LIM-HD family members – specifically *Lhx2, Lhx3*, and *Lhx9* – were also upregulated in *Nipbl^+/-^* hearts (Fig. 9, L to O), suggesting that they too may function as compensatory factors in *Nipbl*-haploinsufficiency. These genes were also modestly upregulated (though not significantly) in *Isl^+/-^*hearts (Fig. 9, M to O), raising the possibility that they may compensate for the loss of *Isl1* itself as well. In *Nipbl^+/-^; Isl1^+/-^* hearts, *Lhx3* and *Lhx9*, were upregulated at even higher levels than in either single mutant (Fig. 9, M to O), potentially reflecting a heightened compensatory response to double haploinsufficiency. Yet, their upregulation was insufficient to prevent the synergistic rise in CHDs (Fig. 6). Together, these findings suggest that *Lhx2, Lhx3*, and *Lhx9* may operate alongside *Isl1* in a broader network of LIM-HD transcription factors that help stabilize chromatin architecture during heart development.

### *Nipbl*-haploinsufficiency in extracardiac tissues contribute to the development of CHDs

We initially asked whether *Nipbl*-haploinsufficiency confined to the SHF is necessary and sufficient to cause CHDs using two Cre lines: *Mef2c-Cre* and *Isl1-Cre*, in which only *Isl1-Cre*- driven *Nipbl*-haploinsufficiency resulted in CHDs. Interestingly, despite recombining less extensively *within* the heart (Fig. 3, A & B), *Isl1-Cre* still produced a greater burden of CHDs than *Mef2c-Cre* (Fig. 2). However, *Isl1-Cre* exhibited much broader recombination outside the heart, including in the pharyngeal arches, dorsal root ganglia (DRG), hindlimbs, and diencephalon (Fig. 3E). While *Mef2c-Cre* also targeted the pharyngeal arches (Fig. 3D), its recombination was spatially more restricted and weaker. These findings reinforce the notion that *Nipbl*-haploinsufficiency in *extracardiac* tissues contributes substantially to CHD pathogenesis (Santos et al., 2016).

The potential role of extracardiac lineages in CHD pathogenesis is supported by known developmental contributions of the pharyngeal arches. These structures give rise to the pharyngeal arch arteries, which remodel into major vessels such as the aortic arch, pulmonary arteries, and brachiocephalic artery (Graham et al., 2023). Neural crest cells migrating through the arches also contribute to outflow tract septation and the smooth muscle lining of the great vessels (Kirby and Waldo, 1995). Disruptions in pharyngeal arch development are a well-established cause of outflow tract defects (Goldmuntz et al., 1998), and as shown in Fig. 6, are a prevalent defect in *Nipbl^+/-^; Isl1^+/-^* mice.

Additional support for the involvement of the pharyngeal mesoderm comes from our identification of *Tbx1*, *Pitx2*, and *Lhx2* among the genes showing those showing additive effects (Fig. 9, D & M). These genes form a regulatory circuit within the pharyngeal mesoderm (Harel et al., 2012): *Tbx1* and *Pitx2* cooperatively activate Lhx2 expression, and *Tbx1* normally negatively regulates *Pitx2*. In our data, *Lhx2* expression was highest in *Nipbl^+/-^* hearts (Fig. 9M), where both *Tbx1* and *Pitx2* were significantly upregulated (Fig. S2, C & K). In contrast, *Tbx1* remained elevated in *Nipbl^+/-^; Isl1^+/-^* hearts, but *Pitx2* expression occurred at WT levels (Fig. S2, C & K). We speculate that the loss of *Pitx2* upregulation limits *Lhx2* activation, despite increased *Tbx1* expression. Given the contributions of the pharyngeal mesoderm to outflow tract morphogenesis (Harel et al., 2012), disruptions of this regulatory circuit may contribute to the increased burden of outflow tract defects observed in *Nipbl^+/-^; Isl1^+/-^* embryos (Fig. 6).

### *Nipbl^+/-^* and *Isl1^+/-^* mice as models for chromatin architecture-based congenital diseases

In CdLS, CHDs occur in ∼30% of affected individuals, with VSDs being the most common, followed by ASDs, pulmonary stenosis, coarctation of the aorta, and complex malformations such as Tetralogy of Fallot (Chatfield et al., 2012). Previous studies using histological analysis or µMRI identified only ASDs in *Nipbl^+/-^* mice (Santos et al., 2016), raising questions about how well this mouse model recapitulates the cardiac phenotypes seen in CdLS. However, application of LSFM revealed that *Nipbl^+/-^* mice actually exhibit a broader array of CHDs, including ASDs, VSDs and OTDs such as overriding aorta and double outlet right ventricle (Fig. 1). These previously undetected phenotypes demonstrate that the *Nipbl^+/-^* mouse more faithfully models the spectrum of CHDs observed in CdLS than previously appreciated, strengthening the utility of this mouse model in studying the developmental origins of human congenital disease.

Given the emerging role of LIM-HD transcription factors like *Isl1* in mediating enhancer–promoter communication via chromatin looping, the *Isl1^⁺/⁻^* mouse also represents a valuable, mechanistically distinct model for studying chromatin structure-sensitive morphogenesis. At E17.5, approximately 22% of *Isl1^⁺/⁻^* mice exhibited ASDs (Fig. 4), indicating that *Isl1* is dosage-sensitive for atrial septation. Since *Isl1* is broadly expressed in multiple embryonic tissues (Fig. 3), it is plausible that *Isl1*-haploinsufficiency may also affect non-cardiac structures. Together, the *Nipbl^+/-^* and *Isl1*^+/-^ mouse models complementary systems for dissecting how disruptions to chromatin architecture and enhancer–promoter dynamics contribute to congenital defects.

### Summary

Taken together, our findings reveal how complex transcriptional interactions can influence the manifestation of CHDs in sensitized genetic backgrounds. We show that even largely additive changes in gene expression can produce synergistic phenotypic outcomes – especially when key buffering mechanisms, such as the proposed *Isl1*-mediated compensation of enhancer-promoter communications (Fig. 10), are lost. By identifying *opposing* gene expression differences as the source of synergistic phenotypes in *Nipbl^+/-^; Isl1^+/-^* hearts, this study identifies a new framework for understanding how gene dosage interactions can result in birth defects, such as CHDs.

## MATERIALS AND METHODS

### Animal Ethics Statement

All animals were handled in accordance with approved procedures as defined by the National Institutes of Health, and all animal work was approved by the Institutional Animal Care and Use Committee of the University of California, Irvine. For collection of mouse tissues, pregnant dams were humanely euthanized by CO_2_ anesthesia followed by cervical dislocation.

### Mouse Lines and Genotyping

All mouse lines were genotyped by polymerase chain reaction (PCR) using genomic DNA obtained from tail biopsies.

#### _NipblFlox/Flox Line_

*The Nipbl*^Flox/Flox^ line (*Nipbl^Gt(EUCE313f02)1.1Hmgu^/Clf*) has an inverted gene trap cassette encoding *b-geo* that is flanked by *Cre* recombinase target sites in intron 1 of both *Nipbl* alleles (Santos et al., 2016). In this inverted orientation, referred herein as *Flox*, there is no trapping of the *Nipbl* gene and *Nipbl* is expressed normally. As a result, the *Nipbl*^Flox/Flox^ line is phenotypically WT. However, when this cassette is exposed to Cre recombinase, the gene trap cassette gets inverted into a non-inverted orientation which we call FIN. In this non-inverted orientation, trapping of the *Nipbl* gene occurs and *b-geo* is expressed as a reporter of successful gene trapping, making the *Nipbl FIN* allele a null allele. The *Nipbl^Flox/Flox^*line was maintained on a C57BL/6J background for more than 15 generations and was genotyped as described in (Santos et al., 2016).

#### Recombinase Mouse Lines

*<u>Nanog-Cre Line:</u>* The *Nanog-Cre* line (Tg(Nanog-cre)#Vlcg, MGI: 5545911) was a gift from Dr. A. Economides of Regeneron Pharmaceuticals (Economides et al., 2013). The *Nanog- Cre* line carries a transgene encoding a *Cre* recombinase downstream of the promoter of the *Nanog* gene and was maintained on C57BL/6J background. The *Nanog-Cre* line was genotyped using the following primers: Forward: 5’- GCA CTG ATT TCG ACC AGG TT-3’ and Reverse: 5’- GCT AAC CAG CGT TTT CGT TC-3’. *Nanog-Cre* amplicon: 200 bp.

*<u>Mef2c-Cre Line:</u>* The *Mef2c-Cre* line (Tg(Mef2c-cre)2Blk, MGI: 3639735) carries a transgene encoding a *Cre* recombinase downstream of the promoter of the *Mef2c* gene, generated using a BAC transgenic approach (Verzi et al., 2005), and was maintained on a C57BL/6J background. The *Mef2c-Cre* line was genotyped using the following primers: Forward: 5’- GGC TGG TTG GGA TAG AGA AAC- 3’ and Reverse: 5’- TTT TCC ATG AGT GAA CGA ACC-3’.

#### *Mef2c-Cre* amplicon: 612 base pairs (bp)

*Isl1^Cre/+^ Line:* The *Isl1^Cre/+^* line (*Isl1^tm1(cre)Sev^*, MGI: 3623159) is a knock-in transgenic that targeted the start codon of the *Isl1* locus (Yang et al., 2006). The *Isl1^Cre/+^* line was maintained on a C57BL/6J background. The *Isl1^Cre/+^*line was genotyped using the following primers: Forward 1: 5’- GCC ACT ATT TGC CAC CTA GC-3’, Reverse 1: 5’- CAA ATC CAA AGA GCC CTG TC-3’, Reverse 2: 5’- AGG CAA ATT TTG GTG TAC GG-3’. *Isl1-WT* amplicon: 434 bp. *Isl1-Cre* amplicon: 300 bp.

*<u>Stra8-iCre Line:</u>* The Stra8-iCre line (Tg(Stra8-icre)1Reb, MGI:3779079) carries a transgene encoding an iCre recombinase downstream of the promoter of the *Stra8* gene (Sadate-Ngatchou et al., 2008) and was maintained on a C57BL/6J background. The *Stra8-iCre* line was genotyped using the following primers: Forward: 5’-AGG TCA TCT TGC TCC TTC CA-3’ and Reverse: 5’-TCC TGT TGT TCA GCT TGC AC-3’. *Stra8-iCre* amplicon: 700 bp.

*Stra8-iCre; Nipbl^Flox/+^ Line:* The *Stra8-iCre; Nipbl^Flox/+^* line was generated by crossing the *Stra8-iCre* line with the *Nipbl^Flox/Flox^* line, which were both maintained on a C57BL/6J background. The *Stra8-iCre; Nipbl^Flox/+^*line produces sperm that are either *Nipbl^+^* or *Nipbl^Flox^*(which are both WT alleles) or *Nipbl^FIN^* (which is a null allele). The *Stra8-iCre; Nipbl^Flox/+^* line was genotyped for *Stra8-Cre* (see *Stra8-iCre* Line) and *Nipbl^Flox/+^* as described above (see *Nipbl^Flox/Flox^* Line).

#### Reporter Mouse Lines

*<u>Td-tomato-EGFP Line:</u>* One reporter line used in this study was *Td-tomato-EGFP* (Gt(ROSA)26Sor^tm4(ACTB-tdTomato,-EGFP)Luo^, MGI:3716464) (Muzumdar et al., 2007). Homozygous *Td-tomato-EGFP* mice on a C57BL/6J background were purchased from The Jackson Laboratory and maintained on a C57BL/6J background. The *Td-tomato-EGFP* line was genotyped using the following primers: Forward 1: 5’-CTC TGC TGC CTC CTG GCT TCT-3’, Reverse 1: 5’-CGA GGC GGA TCA CAA GCA ATA-3’, and Reverse 2: 5’-TCA ATG GGC GGG GGT CGT T-3’. *Td-tomato-WT* amplicon: 330 bp. *Td-tomato-EGFP* amplicon: 250 bp.

*Nipbl^Flox/Flox^; Td-tomato-EGFP Line:* The *Nipbl^Flox/Flox^; Td-tomato-EGFP* line was generated by crossing the *Nipbl^F^*^lox/Flox^ line with the *Td-tomato-EGFP* line. The *Nipbl^Flox/Flox^; Td- tomato-EGFP* line was maintained on a C57BL/6J background and genotyped for *Nipbl^Flox/Flox^* (see *Nipbl^Flox/Flox^* Line) and *Td-tomato-EGFP* (see *Td-tomato-EGFP* Line) as described above.

### Generation of Mouse Embryos

#### For Visualization of Cre-Mediated Recombination Domains and Tissue-Specific Depletion of Nipbl Expression

To generate embryos for the visualization of Cre-mediated recombination domains and tissue-specific depletion of *Nipbl* expression, *Nipbl^Flox/Flox^; Td-tomato-EGFP* doubly homozygous mice were mated with either *Mef2c-Cre* or *Isl1^Cre/+^* mice. When *Nipbl ^Flox/Flox^; Td- tomato-EGFP* doubly homozygous mice are mated with *Mef2c-Cre* mice, the resulting progeny are either *Nipbl^Flox/+^; Td-tomato-EGFP* (which are phenotypically WT) or *Mef2c-Cre; Nipbl^Flox/+^; Td-tomato-EGFP* (which are depleted for *Nipbl* expression in the *Mef2c-Cre* expression domain). When *Nipbl ^Flox/Flox^; Td-tomato-EGFP* doubly homozygous mice are mated with *Isl1^Cre/+^* mice, the resulting progeny are either *Nipbl^Flox/+^; Td-tomato-EGFP* (which are phenotypically WT) or *Isl1^Cre/+^; Nipbl^Flox/+^; Td-tomato-EGFP* (which are depleted for *Nipbl* expression in the *Isl1-Cre* expression domain).

#### For Tissue-Specific Depletion of Nipbl Expression Only

To generate embryos for the tissue-specific depletion of *Nipbl* expression, *Nipbl^Flox/Flox^* mice were mated with either *Mef2c-Cre* or *Isl1^Cre/+^* mice. When *Nipbl ^Flox/Flox^* mice are mated with *Mef2c-Cre* mice, the resulting progeny are either *Nipbl^Flox/+^* (which are phenotypically WT) or *Mef2c-Cre; Nipbl^Flox/+^*(which are depleted for *Nipbl* expression in the *Mef2c-Cre* expression domain). When *Nipbl ^Flox/Flox^*mice are mated with *Isl1^Cre/+^* mice, the resulting progeny are either *Isl1^+/+^*; *Nipbl^Flox/+^* (which are phenotypically WT) or *Isl1^Cre/+^; Nipbl^Flox/+^* (which are depleted for *Nipbl* expression in the *Isl1-Cre* expression domain).

#### For Visualization of Cre-mediated Recombination Domains Only

To generate embryos for the visualization of Cre-mediated recombination domains, *Td- tomato-EGFP* homozygous mice were mated with either *Mef2c-Cre* or *Isl1^Cre/+^* mice. When *Td- tomato-EGFP* doubly homozygous mice are mated with *Mef2c-Cre* mice, the resulting progeny are either *Td-tomato-EGFP* or *Mef2c-Cre; Td-tomato-EGFP*. When *Td-tomato-EGFP* homozygous mice are mated with *Isl1^Cre/+^* mice, the resulting progeny are either *Isl1^+/+^; Td- tomato-EGFP* or *Isl1^Cre/+^; Td-tomato-EGFP*.

#### For Generation of Nipbl^+/-^; Isl1^+/-^ Mouse Embryos

To generate embryos that are either WT, haploinsufficient for *Isl1* (*Isl1^+/-^*), haploinsufficient for *Nipbl* (*Nipbl^+/-^*); or haploinsufficient for both *Nipbl* and *Isl1 (Nipbl^+/-^; Isl1^+/-^)*, we performed in vitro fertilization of oocytes from *Isl1^Cre/+^*females with sperm from *Stra8-iCre*; *Nipbl*^Flox/+^ males mice through the Transgenic Mouse Facility at the University of California Irvine. The resulting fertilized oocytes were implanted into pseudopregnant CD1 females for gestation. *Isl1^Cre/+^*mice encode a *Cre*-recombinase in one allele of *Isl1* and are haploinsufficient for *Isl1*, as shown in Fig. 4D. *Stra8*-*iCre* mice encode a *Cre*-recombinase downstream of the promoter for the *Stra8* gene and *Stra8*-*iCre* recombines in the male germ line. *Nipbl*^Flox/+^ mice have an inverted gene trap cassette encoding *Bgeo* that is flanked by target sites for *Cre*-recombinase in intron 1 of 1 allele of the *Nipbl* gene. In this inverted orientation, which we call *Flox*, there is no trapping of the *Nipbl* gene and *Nipbl* is expressed normally. However, when this cassette is exposed to Cre-recombinase (in this case from the *Stra8-iCre* allele), the gene trap cassette gets inverted into a non-inverted orientation which we call *FIN*. In this non-inverted orientation, trapping of the *Nipbl* gene occurs into the *FIN* conformation and *Bgeo* is expressed as a reporter of successful gene trapping. Thus, *Stra8-iCre; Nipbl*^Flox/+^ male mice make sperm that are either *Nipbl^+^ Nipbl^Flox^* (which are both wildtype alleles and referred to here as *Nipbl^+^*) or *Nipbl*^FIN^ (which is a null allele and referred to here as *Nipbl^-^*) and when sperm from these mice fertilize eggs from *Isl1^Cre/+^* female mice, it results in littermates that are either WT, *Isl1^+/-^, Nipbl^+/-^*, or *Nipbl^+/-^; Isl1^+/-^*).

### Collection of Mouse Hearts

#### Collection of E17.5, E15.5, and E10.5 Mouse Hearts

At either 17.5-,15.5-, or 10.5-days post copulation, pregnant dams were euthanized by CO_2_ inhalation followed by cervical dislocation. The uterine horns were dissected and placed in 1X DEPC PBS on ice, and different tissues were dissected with the aid of a dissection microscope. Tissues were collected for genotyping and stored at -20℃. Hearts for LFSM imaging were dissected out of the body and fixed in 4% Paraformaldehyde (PFA) for 2-4 hours at 4°C before they were transferred into clear, unobstructed brain/body imaging cocktails and computational analysis (CUBIC) solution (Gomez-Gaviro et al., 2017). Hearts for RNA sequencing were dissected out of the body, frozen in liquid nitrogen, and stored at -80℃. Bodies were fixed in 4% PFA at 4℃ overnight, before they were transferred into 1X PBS, and stored at 4℃ for somite counting.

### Zeiss Z.1 Light Sheeting Imaging

CUBIC solution was prepared with the following reagents: 25% w/v urea (Fisher Scientific: AC327380010), 25% w/v N,N,N,N-tetrakis(2-hydroxypropyl)ethylenediamine (Fisher Scientific: T0781500G), 15% w/v TritonX-100 (Fisher Scientific: AC327372500), and 35% w/v nanopure water (Gomez-Gaviro et al., 2017). Hearts were cleared in 1 ml of CUBIC solution in a 2 ml tube at 37°C with a rocking motion at 20 rpm for 5-7 days. CUBIC solution was changed every 3-4 days.

1 ml embedding syringes with a diameter of 4.5mm (Fisher Scientific: 14823434) were prepared by cutting off the tip of syringe with a single-edge No. 9 razor blade (VWR: 55411- 050). Embedding syringes were checked for jagged edges by drawing up agarose, letting the agarose cool, plunging the solidified agarose up and down, and checking under the microscope for cuts to the agarose. Embedding syringes with jagged edges were discarded. Hearts were removed from CUBIC solution and placed in a 1% low melting point agarose (Invitrogen: 16520050) solution (in water). 1% low melting point agarose was aspirated into the embedding syringe and an individual heart was embedded near the aperture. Then the syringe was placed on a flat surface on ice for at least 10-15 minutes. Embedded hearts were then equilibrated by being suspended and completely submerged in CUBIC solution at 4°C for 3-5 days.

Once equilibrated, embedded hearts were suspended in CUBIC solution and imaged using a Zeiss Z.1 Light Sheet microscope and ZEN software (using a 1.45x chamber and 5x illumination and detection objectives) at the Optical Biology Core Facility, at the University of California, Irvine. The distance settings for the illumination objectives were set to -0.4700 and 0.4700 and the configuration for the clearing chamber was selected. Each sample was rotated to image from either a dorsal or ventral view, and the laser focus and exposure time was adjusted to obtain the best quality image. All samples were imaged as a Z-stack and excited at a 100% laser intensity with laser 561 nm for tdTomato excitation and 488nm for EGFP excitation. The Z-stack for each sample was set to capture the whole sample within the Z-stack. Z-stack settings: 746 slices, 6.71µm slice thickness. Z-stacks (CZI files) were processed and converted to Imaris files using the Imaris software converter.

### Histology

Fixed heart samples were embedded in a 24x24x5mm mold (Fisher Scientific: 22363552) with 1.5% LMP agarose. For orientation purposes, a thin black strip of 1.5% agarose made with India Ink (Pro Art PRO-4100) was added. All hearts were also oriented differently for identification purposes and noted on a map. Once all hearts were embedded, the agarose block was carefully removed from the mold, placed in a Omnisette Tissue Casette (Fisher Scientific: 15197700A), and submerged in a container with 1X PBS. Samples were taken to the Department of Pathology at the University of California Irvine to be paraffin-embedded, sectioned serially at 7 μm with 2 sections per slide, and stained with hematoxylin and Eosin Y (H&E). Paraffin-sectioned heart images were obtained with a Discovery V8 stereomicroscope equipped with Axiovision software (Zeiss).

### Heart Analysis

All heart samples from paraffin sectioning and LSFM imaging were positioned or oriented to a canonical angle for heart analysis where the atria are superior to the ventricles, the apex of the heart is inferior, the left atrium and ventricle is on the right, and the right atrium and ventricle is on the left. All samples were assessed from ventral to dorsal. Heart files from LFSM imaging were evaluated for the presence or absence of ASDs or VSDs using the volume viewer plug-in of ImageJ (v. 1.52t) and using a similar criterion outlined in (Santos et al., 2016). Paraffin-sectioned heart images were also evaluated with the same criteria outlined in (Santos et al., 2016). Imaris files of light sheet imaged hearts were evaluated in ImageJ (v. 1.52t) for the presence or absence of ASD or VSD using the volume viewer plug-in and the criterion outlined in (Santos et al., 2016).

Hearts were scored positive for outflow tract defects as follows: Hearts were scored positive for persistent truncus arteriosus (PTA) when the aorta and pulmonary trunk were observed as fused. Hearts were scored positive for double outlet right ventricle (DORV), in which the aorta and pulmonary trunks are observed as both exiting the right ventricle. Hearts were scored positive for overriding aorta (OA) when the aorta was positioned directly over a ventricular septal defect (VSD), instead of over the left ventricle. Fisher’s exact tests were used to calculate *P-*values for the frequency of heart defects (ASD, VSD, and/or outflow tract defects) at E17.5 and E15.5 for all crosses. Ventricular volumes were calculated using the surfaces function on Imaris by manually contouring various sections throughout the ventricles and creating a 3D surface. T-Tests were used to calculate *P*-values for ventricular volume measurements at E17.5 and E15.5.

### X-gal Staining

Whole mount x-gal staining was performed as described (Heffner and Murray, 2024) and imaged with a Zeiss Discovery V8 stereomicroscope equipped with Axiovision software.

### Reverse Transcription-Quantitative Polymerase Chain Reaction

RNA was extracted from E10.5 WT and *Isl1^Cre/+^* hearts using the Monarch Total RNA Miniprep Kit (New England Biolabs). cDNA was made using iSCRIPT reverse transcriptase (BioRad). qRT-PCR for *Isl1* was performed using iTaq SYBR green (BioRad) as per manufacturer’s instructions, *Rpl4* was used as the housekeeping gene. *Rpl4* Primer 1: 5’-ATC TGG ACG GAG AGT GCT TT-3’. *Rpl4* Primer 2: 5’-GGT CGG TGT TCA TCA TCT TG-3’. *Isl1* Primer 1: 5’- ACC AAT TGT CCA ACC ACC AT-3’. *Isl1* Primer 2: 5’- TCC CAT CCC TAA CAA AGC AC-3’.

### RNA Purification, Library Construction, and RNA sequencing

To isolate RNA from 10.5 dpc mouse hearts for library construction, we used the New England BioLabs Monarch Nucleic Acid Purification kit (Cat No: T2010S). Libraries for RNA sequencing were constructed using the Ultra II RNA Library Prep Kit for Illumina (NEB #E7765). RNA libraries were submitted to the UCI Genomics High-Throughput Facility (GHTF) for sequencing on the Illumina Nova Seq6000. RNA libraries were multiplexed and sequenced on a single lane to a depth of approximately 20 million reads each. Sequences were demultiplexed by GHTF.

### Read Mapping and Depth Normalization

Read mapping was performed by GHTF using the GRCm38 mm10 mouse reference genome. Genes with high quality alignments were assigned a transcript count of 1 for each read. GHTF subsequently delivered a matrix of transcript counts for each gene across all libraries. To normalize transcript counts for sequencing depth across all libraries, we calculated the sum of transcript counts in each library and divided these sums by the lowest sum, resulting in a scaling factor for each library. Transcript counts were then multiplied by their library’s respective scaling factors to arrive at depth normalized counts.

### Differential Gene Expression Analysis

To identify differentially expressed genes (DEGs) between pairs of genotypes, we performed an ANOVA test for all genes using normalized transcript counts. Resulting P-values were adjusted for false-discovery using the Benjamini-Hochberg method. Genes with Q-values less than 0.05 indicated that expression of the given gene differed between at least two or more four genotypes. To identify which genotypes differed in its expression of that gene, we performed a Tukey-Kramer test using an alpha of 0.05.

### Principal Component Analysis

Transcript counts from hearts of the same genotype were aggregated together to generate pseudobulked libraries representing hearts of all 4 genotypes: WT, *Isl1^+/-^, Nipbl^+/-^, and Nipbl^+/-^; Isl1^+/-^.* Sequencing depth across all the pseudobulked libraries were normalized by calculating a sum of transcript counts in each library and dividing these sums by the lowest sum, producing a scaling factor for each library. Transcript counts were then multiplied by their library’s respective scaling factors to arrive at depth normalized counts.

The mean expressions of detected genes were calculated across all pseudobulked libraries, from which coefficients of variation were also calculated. The coefficients of variation for all genes were plotted against their mean expression. A LOESS fit was performed across all points in R using the *loess* function of the *stats* package. The command syntax used was *loess(dependent_variable ∼ independent_variable)*, where *dependent_variable* represents a gene’s coefficient of variation and *indepdent_variable* represents a gene’s mean expression. All genes were binned into 32 bins along their mean expression and for each bin, the standard deviation of the coefficient of variation from the LOESS fit was calculated. A Z-score was calculated for gene by dividing its coefficient of variation by the standard deviation of its bin. Genes with Z-score greater than 2 were used as input into principal component analysis (PCA). PCA was performed in R using the *prcomp* function from the *stats* package. The command syntax used was *prcomp(data)*, where *data* represents the expression of genes selected as input into PCA across all hearts of all four genotypes.

### Identification of epistatic and additive gene expression effects

To determine whether combined *Isl1*- and *Nipbl*-haploinsufficiency results in additive effects on gene expression, we used the base linear modeling command from R (lm) to calculate, for each gene, its expected additive effect and the observed deviation from this expectation. A T- test within the command was used to determine whether each deviation was statistically significant. To account for multiple testing, we applied the Benjamini-Hochberg correction to the 770 genes identified earlier in Fig. 8C as being differentially expressed (relative to WT) among *Isl1^+/-^*, *Nipbl^+/-^*; *Nipbl^+/-^; Isl1^+/-^* hearts. Genes with Q-values ≥ 0.05 were considered to follow additive effects, as their observed expression did not significantly differ from expectation. Genes with Q-values < 0.05 were classified as exhibiting epistatic effects.

### Counting transcripts originating from the wildtype allele of *Isl1*

*Isl1-Cre* and *Nipbl^+/−^; Isl1-Cre* hearts carry an *Isl1* allele in which Cre recombinase was inserted 6 bp upstream of the translation initiation codon in Exon 1. Consequently, using the total reads mapped across all *Isl1* exons to measure *Isl1* expression would be inaccurate, as reads mapping to Exon 1 could originate from either the wildtype or mutant allele. Since transcripts from the mutant allele do not span the Exon 1–Exon 2 junction, we quantified reads spanning this junction as specific measure of wildtype *Isl1* expression.

## Acknowledgements

The authors are grateful to Louise Villagomez and Brian Bui for assistance with mouse husbandry, genotyping, and data analysis; to Shimako Kawauchi and the UCI Transgenic Mouse Facility for performing in vitro fertilization; and to Melanie Oakes and the UCI Genomics and Research Technology Hub (GRTH) for RNA-seq library sequencing. Early pilot studies for this project were supported by the UCI Center for Complex Biological Systems and the NSF-Simons Center for Multiscale Cell Fate Research. SC acknowledges support from a training grant award (NINDS/NIH T32NS082174). ADL and ALC dedicate this paper to the memory of Isabel Eva Calof Lander.

## Author contribution

Conceptualization: RS, SC, ADL, ALC

Methodology: RS, SC, ADL, ALC

Software: SC

Validation: MEL, RS, SC, ADL, ALC

Formal Analysis: MEL, RS, SC, ADL, ALC

Investigation: MEL, RS, SC

Resources: ADL, ALC

Data Curation: SC

Writing – Original Draft: RS, SC, MEL, ADL, ALC

Writing – Review & Editing: RS, SC, ADL, ALC

Visualization: MEL, RS, SC, ADL, ALC

Supervision: ADL, ALC

Project Administration: ADL, ALC

Funding Acquisition: ADL, ALC

## Data availability

All relevant data can be found within the article, supplementary information, Sequence Read Archive (Accession: PRJNA1167790), Gene Expression Omnibus (Accession: GSE278563), and Zenodo (DOI: 10.5281/zenodo.15427946).

