## Supplementary Materials for "Interaction between long-range chromatin regulators *Nipbl* & *Isl1* synergistically drives heart defects in mice"

**Author Information**

[1]Dept. of Developmental and Cell Biology, University of California, Irvine, Irvine, California 92697, USA
[2]Center for Complex Biological Systems, University of California, Irvine, Irvine, California 92697, USA
[3]Department of Biomedical Science, Kaiser Permanente Bernard J. Tyson School of Medicine, Pasadena, California, 91101, USA
[4]Dept. of Anatomy and Neurobiology, University of California, Irvine, Irvine, California 92697, USA

[$]These authors contributed equally to this work

**This PDF file includes:**

Figs. S1 & S2
Tables S1 to S18

**Other Supplementary Materials for this manuscript include the following:**

Videos S1 & S2

**Table of Contents**


| | |
|---|---|
| Fig. S1 | Principal component analysis of E10.5 WT, *Isl1*$^{+/-}$, *Nipbl*$^{+/-}$, and *Nipbl*$^{+/-}$; *Isl1*$^{+/-}$ hearts |
| Fig. S2 | Expression of transcription factors showing additive effects of combined *Nipbl*- and *Isl1*-haploinsufficiency |
| Table S1 | Sample identifiers of hearts shown in Fig. 1A |
| Table S2 | Sample identifiers of hearts shown in Fig. 1B |
| Table S3 | Defects observed in hearts analyzed in Fig. 1C. |
| Table S4 | Ventricular volumes of hearts analyzed in Fig. 1D |
| Table S5 | Defects observed in hearts analyzed in Fig. 2B, and method used |
| Table S6 | Ventricular volumes of hearts analyzed in Fig. 2C |
| Table S7 | Defects observed in hearts analyzed in Fig. 2E, and method used |
| Table S8 | Ventricular volumes of hearts analyzed in Fig. 2F |
| Table S9 | Defects observed in hearts analyzed in Fig. 4B |
| Table S10 | Ventricular volumes of hearts analyzed in Fig. 4C |
| Table S11 | qRT-PCR data for *Isl1* in hearts analyzed in Fig. 4D |
| Table S12 | Comparative Ct analysis of *Isl1* in hearts shown in Fig. 4D |
| Table S13 | Sample identifiers of embryos shown in Fig. 5B |
| Table S14 | Sample identifiers of embryos shown in Fig. 5C |
| Table S15 | Signs of fetal demise observed in embryos analyzed in Fig. 5D |
| Table S16 | Crown rump lengths of embryos analyzed in Fig. 5E |
| Table S17 | Defects observed in hearts analyzed in Fig. 6B |
| Table S18 | Sample identifiers of hearts subjected to RNA sequencing in Fig. 7 |
| Video S1 | Video of LSFM images captured through a whole *Nipbl*$^{Flox/+}$ heart |
| Video S2 | Video of LSFM images captured through a whole *Nipbl*$^{FIN/+}$ heart |

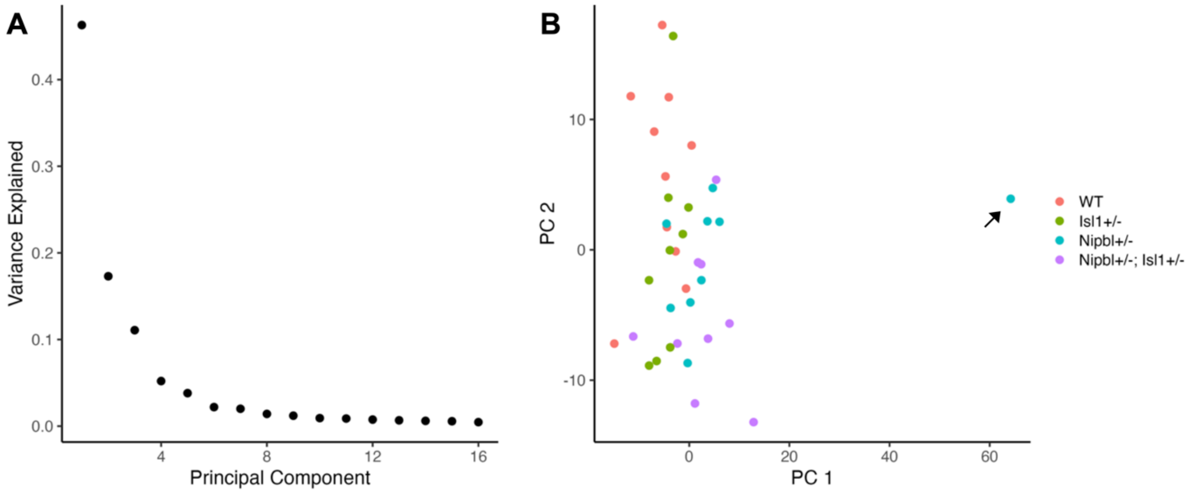


**Figure S1. Principal component analysis of E10.5 WT, *Isl1*$^{+/-}$, *Nipbl*$^{+/-}$, and *Nipbl*$^{+/-}$; *Isl1*$^{+/-}$ hearts.** (A) Variance explained by first 16 principal components among E10.5 WT, *Isl1*$^{+/-}$, *Nipbl*$^{+/-}$, and *Nipbl*$^{+/-}$; *Isl1*$^{+/-}$ hearts. (E) Position of E10.5 WT, *Isl1*$^{+/-}$, *Nipbl*$^{+/-}$, and *Nipbl*$^{+/-}$; *Isl1*$^{+/-}$ hearts along Principal Components 1-2. Arrow points to outlier *Nipbl*$^{+/-}$ heart.

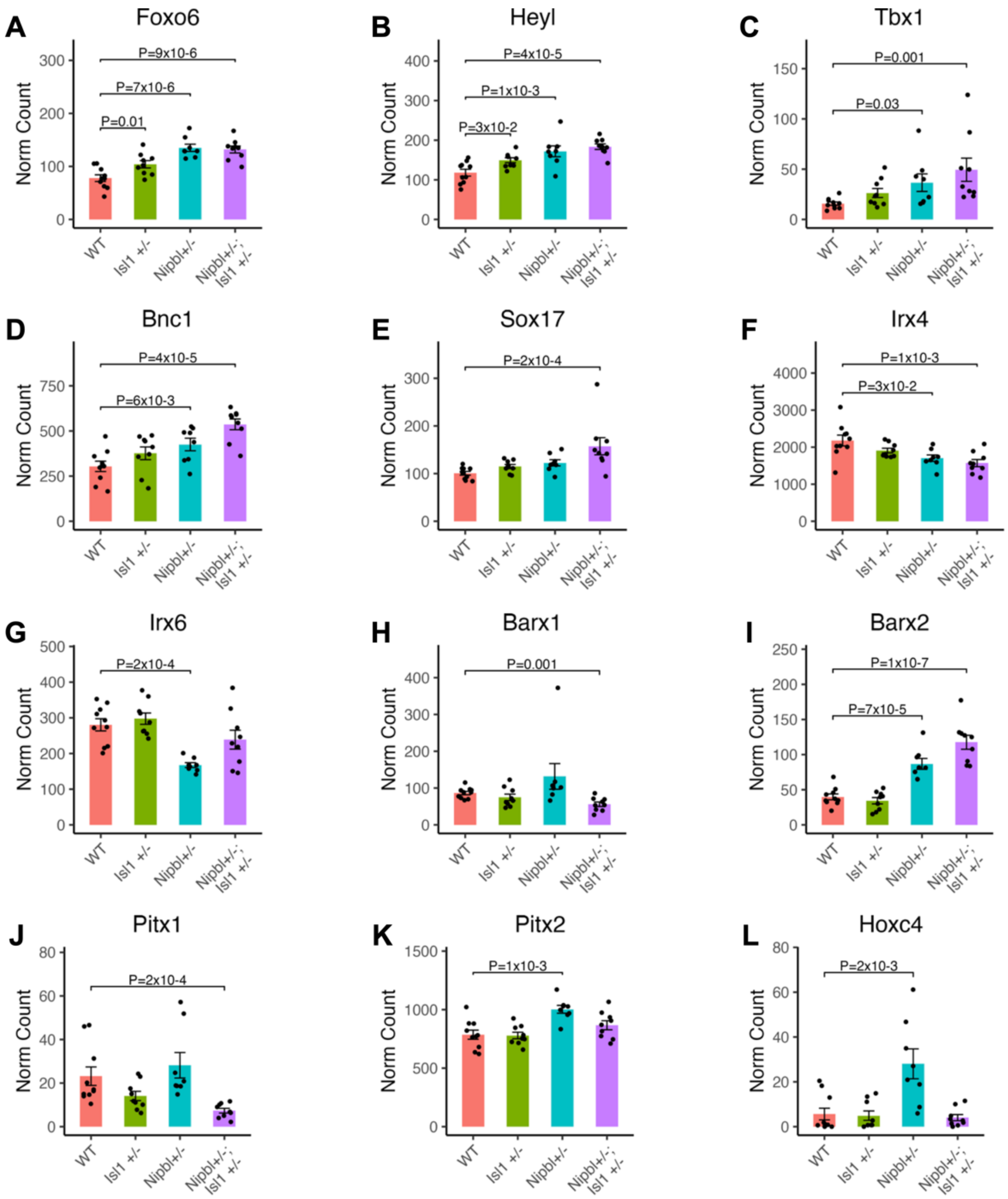


**Figure S2. Expression of transcription factors showing additive effects of combined *Nipbl*- and *Isl1*-haploinsufficiency.** Expression of A) *Foxo6,* B) *Heyl,* C) *Tbx1,* D) *Bnc1,* E) *Sox17,* F) *Irx4,* G) *Irx6,* H) *Barx1,* I) *Barx2,* J) *Pitx1,* K) *Pitx2*, and L) *Hoxc4* in E10.5 WT, $Nipbl^{+/-}$, $Isl1^{+/-}$, and $Nipbl^{+/-}$; $Isl1^{+/-}$ hearts. Error bars show standard error of the mean. P-values from Tukey-Kramer test.

**Table S1.** Sample identifiers of hearts shown in Fig. 1A.

| Age | Column | Sample | Genotype |
|---|---|---|---|
| E17.5 | 1 | D774-81 | Nipbl $^{Flox/+}$; TomatoGFP/+ |
| E17.5 | 2 | D774-84 | Nipbl $^{FIN/+}$; TomatoGFP/+ |
| E17.5 | 3 | D774-84 | Nipbl $^{FIN/+}$; TomatoGFP/+ |
| E17.5 | 4 | D775-87 | Nipbl $^{FIN/+}$; TomatoGFP/+ |

**Table S2.** Sample identifiers of hearts shown in Fig. 1B. Schematic shows sample D741-3.

| Age | Row | Sample | Genotype |
|---|---|---|---|
| E17.5 | 1 | D743-18 | Nipbl $^{Flox/+}$ |
| E17.5 | 2 | D741-3 | Nipbl $^{Fin/+}$ |

**Table S3.** Defects observed in hearts analyzed in Fig. 1C. ASD OP = Atrial septal defect ostium primum type, ASD OS = Atrial septal defect ostium secundum type, DORV = Double outlet right ventricle. OA = Overriding aorta, TGA = Transposition of the Great Arteries, VSD = Ventricular septal defect.

| Age | No | Sample | Genotype | Outcome |
|---|---|---|---|---|
| E17.5 | 1 | D774-79 | Nipbl $^{Flox/+}$; TomatoGFP/+ | No defect |
| E17.5 | 2 | D774-81 | Nipbl $^{Flox/+}$; TomatoGFP/+ | No defect |
| E17.5 | 3 | D774-82 | Nipbl $^{Flox/+}$; TomatoGFP/+ | No defect |
| E17.5 | 4 | D774-83 | Nipbl $^{Flox/+}$; TomatoGFP/+ | No defect |
| E17.5 | 5 | D775-88 | Nipbl $^{Flox/+}$; TomatoGFP/+ | No defect |
| E17.5 | 6 | D775-89 | Nipbl $^{Flox/+}$; TomatoGFP/+ | No defect |
| E17.5 | 7 | D775-90 | Nipbl $^{Flox/+}$; TomatoGFP/+ | No defect |
| E17.5 | 8 | D776-91 | Nipbl $^{Flox/+}$; TomatoGFP/+ | No defect |
| E17.5 | 9 | D776-92 | Nipbl $^{Flox/+}$; TomatoGFP/+ | No defect |
| E17.5 | 10 | D776-93 | Nipbl $^{Flox/+}$; TomatoGFP/+ | No defect |
| E17.5 | 11 | D776-94 | Nipbl $^{Flox/+}$; TomatoGFP/+ | No defect |
| E17.5 | 12 | D776-95 | Nipbl $^{Flox/+}$; TomatoGFP/+ | No defect |
| E17.5 | 13 | D776-96 | Nipbl $^{Flox/+}$; TomatoGFP/+ | No defect |
| E17.5 | 14 | D741-1 | Nipbl $^{Flox/+}$ | No defect |
| E17.5 | 15 | D741-2 | Nipbl $^{Flox/+}$ | No defect |
| E17.5 | 16 | D741-4 | Nipbl $^{Flox/+}$ | ASD OS |
| E17.5 | 17 | D741-5 | Nipbl $^{Flox/+}$ | No defect |
| E17.5 | 18 | D742-10 | Nipbl $^{Flox/+}$ | No defect |
| E17.5 | 19 | D742-11 | Nipbl $^{Flox/+}$ | No defect |
| E17.5 | 20 | D743-15 | Nipbl $^{Flox/+}$ | No defect |
| E17.5 | 21 | D743-18 | Nipbl $^{Flox/+}$ | No defect |
| E17.5 | 22 | D745-21 | Nipbl $^{Flox/+}$ | No defect |
| E17.5 | 23 | D745-23 | Nipbl $^{Flox/+}$ | No defect |
| E17.5 | 24 | D745-24 | Nipbl $^{Flox/+}$ | No defect |
| E17.5 | 25 | D745-25 | Nipbl $^{Flox/+}$ | No defect |

| | | | | |
|---|---|---|---|---|
| E17.5 | 26 | D745-26 | Nipbl $^{Flox/+}$ | No defect |
| E17.5 | 27 | D753-34 | Nipbl $^{Flox/+}$ | No defect |
| E17.5 | 28 | D753-35 | Nipbl $^{Flox/+}$ | No defect |
| E17.5 | 29 | D753-36 | Nipbl $^{Flox/+}$ | No defect |
| E17.5 | 30 | D753-37 | Nipbl $^{Flox/+}$ | No defect |
| E17.5 | 31 | D753-38 | Nipbl $^{Flox/+}$ | No defect |
| E17.5 | 32 | D753-39 | Nipbl $^{Flox/+}$ | No defect |
| E17.5 | 33 | D753-40 | Nipbl $^{Flox/+}$ | No defect |
| E17.5 | 34 | D754-41 | Nipbl $^{Flox/+}$ | No defect |
| E17.5 | 35 | D754-44 | Nipbl $^{Flox/+}$ | No defect |
| E17.5 | 36 | D754-46 | Nipbl $^{Flox/+}$ | No defect |
| E17.5 | 37 | D755-47 | Nipbl $^{Flox/+}$ | No defect |
| E17.5 | 38 | D755-48 | Nipbl $^{Flox/+}$ | No defect |
| E17.5 | 39 | D755-50 | Nipbl $^{Flox/+}$ | No defect |
| E17.5 | 40 | D756-53 | Nipbl $^{Flox/+}$ | No defect |
| E17.5 | 41 | D756-56 | Nipbl $^{Flox/+}$ | No defect |
| E17.5 | 42 | D756-59 | Nipbl $^{Flox/+}$ | No defect |
| E17.5 | 43 | D761--62 | Nipbl $^{Flox/+}$ | No defect |
| E17.5 | 44 | D761--63 | Nipbl $^{Flox/+}$ | No defect |
| E17.5 | 45 | D761--64 | Nipbl $^{Flox/+}$ | No defect |
| E17.5 | 46 | D764-66 | Nipbl $^{Flox/+}$ | No defect |
| E17.5 | 47 | D764-70 | Nipbl $^{Flox/+}$ | No defect |
| E17.5 | 48 | D764-71 | Nipbl $^{Flox/+}$ | No defect |
| E17.5 | 49 | D764-72 | Nipbl $^{Flox/+}$ | No defect |
| E17.5 | 50 | D764-74 | Nipbl $^{Flox/+}$ | No defect |
| E17.5 | 51 | D768-67 | Nipbl $^{Flox/+}$ | No defect |
| E17.5 | 52 | D769-71 | Nipbl $^{Flox/+}$ | No defect |
| E17.5 | 53 | D769-73 | Nipbl $^{Flox/+}$ | No defect |
| E17.5 | 54 | D769-75 | Nipbl $^{Flox/+}$ | No defect |

| | | | | |
|---|---|---|---|---|
| E17.5 | 55 | D769-78 | Nipbl $^{Flox/+}$ | No defect |
| E17.5 | 1 | D774-80 | Nipbl $^{FIN/+}$; TomatoGFP/+ | No defect |
| E17.5 | 2 | D774-84 | Nipbl $^{FIN/+}$; TomatoGFP/+ | DORV VSD, ASD OP |
| E17.5 | 3 | D774-85 | Nipbl $^{FIN/+}$; TomatoGFP/+ | VSD |
| E17.5 | 4 | D775-86 | Nipbl $^{FIN/+}$; TomatoGFP/+ | No defect |
| E17.5 | 5 | D775-87 | Nipbl $^{FIN/+}$; TomatoGFP/+ | OA;VSD; ASD OP |
| E17.5 | 6 | D741-3 | Nipbl $^{Fin/+}$ | ASD OS, VSD, DORV |
| E17.5 | 7 | D741-6 | Nipbl $^{Fin/+}$ | VSD, TGA, ASD OP |
| E17.5 | 8 | D742-7 | Nipbl $^{Fin/+}$ | No defect |
| E17.5 | 9 | D742-8 | Nipbl $^{Fin/+}$ | No defect |
| E17.5 | 10 | D742-9 | Nipbl $^{Fin/+}$ | No defect |
| E17.5 | 11 | D742-12 | Nipbl $^{Fin/+}$ | No defect |
| E17.5 | 12 | D742-13 | Nipbl $^{Fin/+}$ | No defect |
| E17.5 | 13 | D743-14 | Nipbl $^{Fin/+}$ | No defect |
| E17.5 | 14 | D743-16 | Nipbl $^{Fin/+}$ | No defect |
| E17.5 | 15 | D743-17 | Nipbl $^{Fin/+}$ | No defect |
| E17.5 | 16 | D743-19 | Nipbl $^{Fin/+}$ | No defect |
| E17.5 | 17 | D743-20 | Nipbl $^{Fin/+}$ | No defect |
| E17.5 | 18 | D745-22 | Nipbl $^{Fin/+}$ | VSD, TGA, ASD OP |
| E17.5 | 19 | D753-31 | Nipbl $^{Fin/+}$ | No defect |
| E17.5 | 20 | D754-42 | Nipbl $^{Fin/+}$ | No defect |
| E17.5 | 21 | D754-43 | Nipbl $^{Fin/+}$ | No defect |
| E17.5 | 22 | D754-45 | Nipbl $^{Fin/+}$ | No defect |
| E17.5 | 23 | D755-49 | Nipbl $^{Fin/+}$ | No defect |
| E17.5 | 24 | D755-52 | Nipbl $^{Fin/+}$ | No defect |
| E17.5 | 25 | D756-54 | Nipbl $^{Fin/+}$ | No defect |
| E17.5 | 26 | D756-55 | Nipbl $^{Fin/+}$ | No defect |
| E17.5 | 27 | D756-57 | Nipbl $^{Fin/+}$ | VSD, DORV |
| E17.5 | 28 | D756-58 | Nipbl $^{Fin/+}$ | No defect |

| | | | | |
|---|---|---|---|---|
| E17.5 | 29 | D761--60 | Nipbl $^{Fin/+}$ | No defect |
| E17.5 | 30 | D761--65 | Nipbl $^{Fin/+}$ | ASD OS |
| E17.5 | 31 | D764-67 | Nipbl $^{Fin/+}$ | VSD; OA |
| E17.5 | 32 | D764-68 | Nipbl $^{Fin/+}$ | ASD OP |
| E17.5 | 33 | D764-69 | Nipbl $^{Fin/+}$ | ASD OP |
| E17.5 | 34 | D764-73 | Nipbl $^{Fin/+}$ | No defect |
| E17.5 | 35 | D768-66 | Nipbl $^{Fin/+}$ | ASD OP |
| E17.5 | 36 | D768-68 | Nipbl $^{Fin/+}$ | No defect |
| E17.5 | 37 | D768-69 | Nipbl $^{Fin/+}$ | No defect |
| E17.5 | 38 | D769-70 | Nipbl $^{Fin/+}$ | ASD OP |
| E17.5 | 39 | D769-72 | Nipbl $^{Fin/+}$ | No defect |
| E17.5 | 40 | D769-74 | Nipbl $^{Fin/+}$ | No defect |
| E17.5 | 41 | D769-76 | Nipbl $^{Fin/+}$ | ASD OS |
| E17.5 | 42 | D769-77 | Nipbl $^{Fin/+}$ | VSD, OA, ASD OP |

**Table S4.** Ventricular volumes of hearts analyzed in Fig. 1D.

| Age | No | Sample | Genotype | Ventricular Volume ($mm^3$) |
|---|---|---|---|---|
| E17.5 | 1 | D764-67 | Nipbl $^{Fin/+}$ | 5.57 |
| E17.5 | 2 | D764-68 | Nipbl $^{Fin/+}$ | 4.95 |
| E17.5 | 3 | D764-69 | Nipbl $^{Fin/+}$ | 4.19 |
| E17.5 | 4 | D764-73 | Nipbl $^{Fin/+}$ | 4.77 |
| E17.5 | 5 | D769-70 | Nipbl $^{Fin/+}$ | 4.99 |
| E17.5 | 6 | D769-72 | Nipbl $^{Fin/+}$ | 5.87 |
| E17.5 | 7 | D769-76 | Nipbl $^{Fin/+}$ | 5.68 |
| E17.5 | 8 | D769-77 | Nipbl $^{Fin/+}$ | 4.03 |
| E17.5 | 1 | D764-66 | Nipbl $^{Flox/+}$ | 7.74 |
| E17.5 | 2 | D764-70 | Nipbl $^{Flox/+}$ | 7.49 |
| E17.5 | 3 | D764-71 | Nipbl $^{Flox/+}$ | 7.08 |
| E17.5 | 4 | D764-72 | Nipbl $^{Flox/+}$ | 6.8 |
| E17.5 | 5 | D764-74 | Nipbl $^{Flox/+}$ | 7.04 |
| E17.5 | 6 | D769-71 | Nipbl $^{Flox/+}$ | 5.67 |
| E17.5 | 7 | D769-73 | Nipbl $^{Flox/+}$ | 5.83 |
| E17.5 | 8 | D769-75 | Nipbl $^{Flox/+}$ | 6.94 |
| E17.5 | 9 | D769-78 | Nipbl $^{Flox/+}$ | 6.84 |

**Table S5.** Defects observed in hearts analyzed in Fig. 2B, and method used. ASD = Atrial septal defect. LS = Light sheet, S = Section.

| Age | No | Sample | Genotype | Outcome | Method |
|---|---|---|---|---|---|
| E17.5 | 1 | D694-1 | Mef2c-cre/+; Nipbl $^{Flox/+}$ | No defect | LS |
| E17.5 | 2 | D694-3 | Mef2c-cre/+; Nipbl $^{Flox/+}$ | No defect | S |
| E17.5 | 3 | D694-4 | Mef2c-cre/+; Nipbl $^{Flox/+}$ | No defect | LS |
| E17.5 | 4 | D694-6 | Mef2c-cre/+; Nipbl $^{Flox/+}$ | No defect | LS |
| E17.5 | 5 | D694-7 | Mef2c-cre/+; Nipbl $^{Flox/+}$ | No defect | LS |
| E17.5 | 6 | D694-8 | Mef2c-cre/+; Nipbl $^{Flox/+}$ | ASD | S |
| E17.5 | 7 | D694-9 | Mef2c-cre/+; Nipbl $^{Flox/+}$ | No defect | LS |
| E17.5 | 8 | D696-11 | Mef2c-cre/+; Nipbl $^{Flox/+}$ | No defect | S |
| E17.5 | 9 | D696-14 | Mef2c-cre/+; Nipbl $^{Flox/+}$ | No defect | S |
| E17.5 | 10 | D696-15 | Mef2c-cre/+; Nipbl $^{Flox/+}$ | ASD | S |
| E17.5 | 11 | D696-16 | Mef2c-cre/+; Nipbl $^{Flox/+}$ | No defect | LS |
| E17.5 | 12 | D696-17 | Mef2c-cre/+; Nipbl $^{Flox/+}$ | No defect | LS |
| E17.5 | 13 | D696-18 | Mef2c-cre/+; Nipbl $^{Flox/+}$ | No defect | LS |
| E17.5 | 14 | D698-23 | Mef2c-cre/+; Nipbl $^{Flox/+}$ | No defect | S |
| E17.5 | 15 | D698-24 | Mef2c-cre/+; Nipbl $^{Flox/+}$ | No defect | S |
| E17.5 | 16 | D698-25 | Mef2c-cre/+; Nipbl $^{Flox/+}$ | No defect | LS |
| E17.5 | 17 | D699-31 | Mef2c-cre/+; Nipbl $^{Flox/+}$ | No defect | LS |
| E17.5 | 18 | D700-35 | Mef2c-cre/+; Nipbl $^{Flox/+}$ | No defect | S |
| E17.5 | 19 | D700-36 | Mef2c-cre/+; Nipbl $^{Flox/+}$ | No defect | LS |
| E17.5 | 20 | D700-37 | Mef2c-cre/+; Nipbl $^{Flox/+}$ | No defect | LS |
| E17.5 | 21 | D700-39 | Mef2c-cre/+; Nipbl $^{Flox/+}$ | No defect | LS |
| E17.5 | 22 | D701-42 | Mef2c-cre/+; Nipbl $^{Flox/+}$ | No defect | S |
| E17.5 | 23 | D701-44 | Mef2c-cre/+; Nipbl $^{Flox/+}$ | No defect | S |
| E17.5 | 24 | D723-46 | Mef2c-cre/+; Nipbl $^{Flox/+}$ | No defect | S |
| E17.5 | 25 | D723-47 | Mef2c-cre/+; Nipbl $^{Flox/+}$ | No defect | S |
| E17.5 | 26 | D723-50 | Mef2c-cre/+; Nipbl $^{Flox/+}$ | No defect | S |

| | | | | | |
|---|---|---|---|---|---|
| E17.5 | 27 | D723-53 | Mef2c-cre/+; Nipbl $^{Flox/+}$ | No defect | S |
| E17.5 | 28 | D724-58 | Mef2c-cre/+; Nipbl $^{Flox/+}$ | No defect | S |
| E17.5 | 29 | D724-60 | Mef2c-cre/+; Nipbl $^{Flox/+}$ | No defect | S |
| E17.5 | 30 | D724-61 | Mef2c-cre/+; Nipbl $^{Flox/+}$ | No defect | S |
| E17.5 | 31 | D724-62 | Mef2c-cre/+; Nipbl $^{Flox/+}$ | No defect | S |
| E17.5 | 32 | D725-63 | Mef2c-cre/+; Nipbl $^{Flox/+}$ | No defect | S |
| E17.5 | 33 | D725-66 | Mef2c-cre/+; Nipbl $^{Flox/+}$ | No defect | S |
| E17.5 | 34 | D725-69 | Mef2c-cre/+; Nipbl $^{Flox/+}$ | No defect | S |
| E17.5 | 35 | D725-70 | Mef2c-cre/+; Nipbl $^{Flox/+}$ | No defect | S |
| E17.5 | 36 | D725-71 | Mef2c-cre/+; Nipbl $^{Flox/+}$ | No defect | S |
| E17.5 | 37 | D758-74 | Mef2c-cre/+; Nipbl $^{Flox/+}$; Tomato GFP/+ | No defect | LS |
| E17.5 | 38 | D758-75 | Mef2c-cre/+; Nipbl $^{Flox/+}$; Tomato GFP/+ | No defect | LS |
| E17.5 | 39 | D758-76 | Mef2c-cre/+; Nipbl $^{Flox/+}$; Tomato GFP/+ | No defect | LS |
| E17.5 | 40 | D758-77 | Mef2c-cre/+; Nipbl $^{Flox/+}$; Tomato GFP/+ | No defect | LS |
| E17.5 | 41 | D758-79 | Mef2c-cre/+; Nipbl $^{Flox/+}$; Tomato GFP/+ | No defect | LS |
| E17.5 | 42 | D760-82 | Mef2c-cre/+; Nipbl $^{Flox/+}$; Tomato GFP/+ | No defect | LS |
| E17.5 | 43 | D760-84 | Mef2c-cre/+; Nipbl $^{Flox/+}$; Tomato GFP/+ | No defect | LS |
| E17.5 | 44 | D760-88 | Mef2c-cre/+; Nipbl $^{Flox/+}$; Tomato GFP/+ | No defect | LS |
| E17.5 | 1 | D694-2 | Nipbl $^{Flox/+}$ | No defect | S |
| E17.5 | 2 | D694-5 | Nipbl $^{Flox/+}$ | No defect | S |
| E17.5 | 3 | D696-10 | Nipbl $^{Flox/+}$ | No defect | S |
| E17.5 | 4 | D696-12 | Nipbl $^{Flox/+}$ | No defect | S |
| E17.5 | 5 | D696-13 | Nipbl $^{Flox/+}$ | No defect | LS |
| E17.5 | 6 | D698-19 | Nipbl $^{Flox/+}$ | No defect | S |
| E17.5 | 7 | D698-20 | Nipbl $^{Flox/+}$ | No defect | LS |
| E17.5 | 8 | D698-21 | Nipbl $^{Flox/+}$ | No defect | LS |
| E17.5 | 9 | D698-22 | Nipbl $^{Flox/+}$ | No defect | S |
| E17.5 | 10 | D699-26 | Nipbl $^{Flox/+}$ | No defect | S |
| E17.5 | 11 | D699-27 | Nipbl $^{Flox/+}$ | No defect | LS |

| | | | | | |
|---|---|---|---|---|---|
| E17.5 | 12 | D699-28 | Nipbl $^{Flox/+}$ | No defect | LS |
| E17.5 | 13 | D699-29 | Nipbl $^{Flox/+}$ | No defect | LS |
| E17.5 | 14 | D699-30 | Nipbl $^{Flox/+}$ | No defect | LS |
| E17.5 | 15 | D699-32 | Nipbl $^{Flox/+}$ | No defect | LS |
| E17.5 | 16 | D700-33 | Nipbl $^{Flox/+}$ | No defect | S |
| E17.5 | 17 | D700-34 | Nipbl $^{Flox/+}$ | No defect | LS |
| E17.5 | 18 | D701-40 | Nipbl $^{Flox/+}$ | No defect | S |
| E17.5 | 19 | D723-45 | Nipbl $^{Flox/+}$ | No defect | S |
| E17.5 | 20 | D723-48 | Nipbl $^{Flox/+}$ | No defect | S |
| E17.5 | 21 | D724-54 | Nipbl $^{Flox/+}$ | No defect | S |
| E17.5 | 22 | D724-59 | Nipbl $^{Flox/+}$ | No defect | S |
| E17.5 | 23 | D725-64 | Nipbl $^{Flox/+}$ | No defect | S |
| E17.5 | 24 | D725-65 | Nipbl $^{Flox/+}$ | No defect | S |
| E17.5 | 25 | D725-67 | Nipbl $^{Flox/+}$ | No defect | S |
| E17.5 | 26 | D725-68 | Nipbl $^{Flox/+}$ | No defect | S |
| E17.5 | 27 | D752-1 | Nipbl $^{Flox/+}$; Tomato GFP/+ | No defect | LS |
| E17.5 | 28 | D752-2 | Nipbl $^{Flox/+}$; Tomato GFP/+ | No defect | LS |
| E17.5 | 29 | D752-3 | Nipbl $^{Flox/+}$; Tomato GFP/+ | No defect | LS |
| E17.5 | 30 | D758-73 | Nipbl $^{Flox/+}$; Tomato GFP/+ | No defect | LS |
| E17.5 | 31 | D758-78 | Nipbl $^{Flox/+}$; Tomato GFP/+ | No defect | LS |
| E17.5 | 32 | D760-80 | Nipbl $^{Flox/+}$; Tomato GFP/+ | No defect | LS |
| E17.5 | 33 | D760-81 | Nipbl $^{Flox/+}$; Tomato GFP/+ | No defect | LS |
| E17.5 | 34 | D760-83 | Nipbl $^{Flox/+}$; Tomato GFP/+ | No defect | LS |
| E17.5 | 35 | D760-85 | Nipbl $^{Flox/+}$; Tomato GFP/+ | No defect | LS |
| E17.5 | 36 | D760-86 | Nipbl $^{Flox/+}$; Tomato GFP/+ | No defect | LS |
| E17.5 | 37 | D760-87 | Nipbl $^{Flox/+}$; Tomato GFP/+ | No defect | LS |

**Table S6.** Ventricular volumes of hearts analyzed in Fig. 2C.

| Age | No | Sample | Genotype | Ventricular Volume ($mm^3$) |
|---|---|---|---|---|
| E17.5 | 1 | D758-74 | Mef2c-cre/+; Nipbl $^{Flox/+}$; Tomato GFP/+ | 8.336 |
| E17.5 | 2 | D758-75 | Mef2c-cre/+; Nipbl $^{Flox/+}$; Tomato GFP/+ | 9.99 |
| E17.5 | 3 | D758-76 | Mef2c-cre/+; Nipbl $^{Flox/+}$; Tomato GFP/+ | 7.936 |
| E17.5 | 4 | D758-77 | Mef2c-cre/+; Nipbl $^{Flox/+}$; Tomato GFP/+ | 7.68 |
| E17.5 | 5 | D758-79 | Mef2c-cre/+; Nipbl $^{Flox/+}$; Tomato GFP/+ | 9.12 |
| E17.5 | 6 | D760-82 | Mef2c-cre/+; Nipbl $^{Flox/+}$; Tomato GFP/+ | 11.03 |
| E17.5 | 7 | D760-84 | Mef2c-cre/+; Nipbl $^{Flox/+}$; Tomato GFP/+ | 12.62 |
| E17.5 | 8 | D760-88 | Mef2c-cre/+; Nipbl $^{Flox/+}$; Tomato GFP/+ | 10.32 |
| E17.5 | 9 | D694-1 | Mef2c-cre/+; Nipbl $^{Flox/+}$; Tomato GFP/+ | 7.91 |
| E17.5 | 10 | D698-25 | Mef2c-cre/+; Nipbl $^{Flox/+}$; Tomato GFP/+ | 6.42 |
| E17.5 | 11 | D699-31 | Mef2c-cre/+; Nipbl $^{Flox/+}$; Tomato GFP/+ | 6.95 |
| E17.5 | 1 | D758-73 | Nipbl $^{Flox/+}$; Tomato GFP/+ | 8.64 |
| E17.5 | 2 | D758-78 | Nipbl $^{Flox/+}$; Tomato GFP/+ | 8.42 |
| E17.5 | 3 | D760-80 | Nipbl $^{Flox/+}$; Tomato GFP/+ | 9.33 |
| E17.5 | 4 | D760-81 | Nipbl $^{Flox/+}$; Tomato GFP/+ | 10.92 |
| E17.5 | 5 | D760-83 | Nipbl $^{Flox/+}$; Tomato GFP/+ | 10 |
| E17.5 | 6 | D760-85 | Nipbl $^{Flox/+}$; Tomato GFP/+ | 12.81 |
| E17.5 | 7 | D760-86 | Nipbl $^{Flox/+}$; Tomato GFP/+ | 11.46 |
| E17.5 | 8 | D760-87 | Nipbl $^{Flox/+}$; Tomato GFP/+ | 10.63 |
| E17.5 | 9 | D698-20 | Nipbl $^{Flox/+}$; Tomato GFP/+ | 7.62 |
| E17.5 | 10 | D698-21 | Nipbl $^{Flox/+}$; Tomato GFP/+ | 6.9 |
| E17.5 | 11 | D698-27 | Nipbl $^{Flox/+}$; Tomato GFP/+ | 7.78 |
| E17.5 | 12 | D698-28 | Nipbl $^{Flox/+}$; Tomato GFP/+ | 7.19 |
| E17.5 | 13 | D699-29 | Nipbl $^{Flox/+}$; Tomato GFP/+ | 9.36 |
| E17.5 | 14 | D699-30 | Nipbl $^{Flox/+}$; Tomato GFP/+ | 6.71 |
| E17.5 | 15 | D699-32 | Nipbl $^{Flox/+}$; Tomato GFP/+ | 8.52 |

**Table S7.** Defects observed in hearts analyzed in Fig. 2E, and method used. ASD OP = Atrial septal defect ostium primum type, ASD OS = Atrial septal defect ostium secundum type, OA = Overriding aorta, PTA = Persistent truncus arteriosis, VSD = Ventricular septal defect.

| Age | No | Sample | Genotype | Outcome | Method |
|---|---|---|---|---|---|
| E17.5 | 1 | D750-3 | Nipbl $^{Flox/+}$; Tomato GFP/+ | NO DEFECT | Light Sheet |
| E17.5 | 2 | D750-5 | Nipbl $^{Flox/+}$; Tomato GFP/+ | NO DEFECT | Light Sheet |
| E17.5 | 3 | D750-7 | Nipbl $^{Flox/+}$; Tomato GFP/+ | NO DEFECT | Light Sheet |
| E17.5 | 4 | D732-27 | Nipbl $^{Flox/+}$; Tomato GFP/+ | NO DEFECT | Section |
| E17.5 | 5 | D773-33 | Nipbl $^{Flox/+}$; Tomato GFP/+ | NO DEFECT | Light Sheet |
| E17.5 | 6 | D773-35 | Nipbl $^{Flox/+}$; Tomato GFP/+ | NO DEFECT | Light Sheet |
| E17.5 | 7 | D777-40 | Nipbl $^{Flox/+}$; Tomato GFP/+ | NO DEFECT | Light Sheet |
| E17.5 | 8 | D777-42 | Nipbl $^{Flox/+}$; Tomato GFP/+ | NO DEFECT | Light Sheet |
| E17.5 | 9 | D777-44 | Nipbl $^{Flox/+}$; Tomato GFP/+ | NO DEFECT | Light Sheet |
| E17.5 | 10 | D777-46 | Nipbl $^{Flox/+}$; Tomato GFP/+ | NO DEFECT | Light Sheet |
| E17.5 | 11 | D777-47 | Nipbl $^{Flox/+}$; Tomato GFP/+ | NO DEFECT | Light Sheet |
| E17.5 | 12 | D778-48 | Nipbl $^{Flox/+}$; Tomato GFP/+ | ASD OS | Light Sheet |
| E17.5 | 13 | D778-50 | Nipbl $^{Flox/+}$; Tomato GFP/+ | NO DEFECT | Light Sheet |
| E17.5 | 14 | D778-53 | Nipbl $^{Flox/+}$; Tomato GFP/+ | NO DEFECT | Light Sheet |
| E17.5 | 15 | D779-54 | Nipbl $^{Flox/+}$; Tomato GFP/+ | NO DEFECT | Light Sheet |
| E17.5 | 16 | D779-55 | Nipbl $^{Flox/+}$; Tomato GFP/+ | NO DEFECT | Light Sheet |
| E17.5 | 17 | D779-57 | Nipbl $^{Flox/+}$; Tomato GFP/+ | NO DEFECT | Light Sheet |
| E17.5 | 18 | D779-58 | Nipbl $^{Flox/+}$; Tomato GFP/+ | NO DEFECT | Light Sheet |
| E17.5 | 19 | D779-59 | Nipbl $^{Flox/+}$; Tomato GFP/+ | NO DEFECT | Light Sheet |
| E17.5 | 20 | D779-60 | Nipbl $^{Flox/+}$; Tomato GFP/+ | NO DEFECT | Light Sheet |
| E17.5 | 21 | D726-2 | Nipbl $^{Flox/+}$; Tomato GFP/+ | NO DEFECT | Section |
| E17.5 | 22 | D726-6 | Nipbl $^{Flox/+}$; Tomato GFP/+ | NO DEFECT | Light Sheet |
| E17.5 | 23 | D751-9 | Nipbl $^{Flox/+}$; Tomato GFP/+ | NO DEFECT | Light Sheet |
| E17.5 | 24 | D729-8 | Nipbl $^{Flox/+}$; Tomato GFP/+ | NO DEFECT | Light Sheet |
| E17.5 | 25 | D729-11 | Nipbl $^{Flox/+}$; Tomato GFP/+ | NO DEFECT | Light Sheet |

| | | | | | |
|---|---|---|---|---|---|
| E17.5 | 26 | D729-12 | Nipbl $^{Flox/+}$; Tomato GFP/+ | NO DEFECT | Light Sheet |
| E17.5 | 27 | D728-2 | Nipbl $^{Flox/+}$; Tomato GFP/+ | NO DEFECT | Light Sheet |
| E17.5 | 28 | D728-3 | Nipbl $^{Flox/+}$; Tomato GFP/+ | NO DEFECT | Light Sheet |
| E17.5 | 29 | D728-6 | Nipbl $^{Flox/+}$; Tomato GFP/+ | NO DEFECT | Light Sheet |
| E17.5 | 30 | D728-7 | Nipbl $^{Flox/+}$; Tomato GFP/+ | NO DEFECT | Light Sheet |
| E17.5 | 31 | D728-8 | Nipbl $^{Flox/+}$; Tomato GFP/+ | NO DEFECT | Light Sheet |
| E17.5 | 32 | D730-10 | Nipbl $^{Flox/+}$; Tomato GFP/+ | NO DEFECT | Light Sheet |
| E17.5 | 33 | D757-15 | Nipbl $^{Flox/+}$; Tomato GFP/+ | NO DEFECT | Light Sheet |
| E17.5 | 34 | D757-16 | Nipbl $^{Flox/+}$; Tomato GFP/+ | NO DEFECT | Light Sheet |
| E17.5 | 35 | D757-17 | Nipbl $^{Flox/+}$; Tomato GFP/+ | NO DEFECT | Light Sheet |
| E17.5 | 36 | D757-18 | Nipbl $^{Flox/+}$; Tomato GFP/+ | NO DEFECT | Light Sheet |
| E17.5 | 37 | D757-20 | Nipbl $^{Flox/+}$; Tomato GFP/+ | NO DEFECT | Light Sheet |
| E17.5 | 38 | D757-22 | Nipbl $^{Flox/+}$; Tomato GFP/+ | NO DEFECT | Light Sheet |
| E17.5 | 39 | D759-23 | Nipbl $^{Flox/+}$; Tomato GFP/+ | NO DEFECT | Light Sheet |
| E17.5 | 40 | D759-24 | Nipbl $^{Flox/+}$; Tomato GFP/+ | NO DEFECT | Light Sheet |
| E17.5 | 41 | D759-27 | Nipbl $^{Flox/+}$; Tomato GFP/+ | NO DEFECT | Light Sheet |
| E17.5 | 42 | D759-28 | Nipbl $^{Flox/+}$; Tomato GFP/+ | NO DEFECT | Light Sheet |
| E17.5 | 43 | D762-29 | Nipbl $^{Flox/+}$; Tomato GFP/+ | NO DEFECT | Light Sheet |
| E17.5 | 44 | D762-32 | Nipbl $^{Flox/+}$; Tomato GFP/+ | NO DEFECT | Light Sheet |
| E17.5 | 45 | D731-17 | Nipbl $^{Flox/+}$ | NO DEFECT | Section |
| E17.5 | 46 | D731-19 | Nipbl $^{Flox/+}$ | NO DEFECT | Light Sheet |
| E17.5 | 47 | D731-21 | Nipbl $^{Flox/+}$ | NO DEFECT | Section |
| E17.5 | 48 | D731-22 | Nipbl $^{Flox/+}$ | NO DEFECT | Light Sheet |
| E17.5 | 49 | D732-25 | Nipbl $^{Flox/+}$ | NO DEFECT | Light Sheet |
| E17.5 | 50 | D732-26 | Nipbl $^{Flox/+}$ | NO DEFECT | Light Sheet |
| E17.5 | 51 | D732-29 | Nipbl $^{Flox/+}$ | NO DEFECT | Light Sheet |
| E17.5 | 52 | D732-31 | Nipbl $^{Flox/+}$ | NO DEFECT | Light Sheet |
| E17.5 | 1 | D750-1 | Isl1 $^{Cre/+}$; Nipbl $^{Flox/+}$; Tomato GFP/+ | NO DEFECT | Light Sheet |
| E17.5 | 2 | D750-2 | Isl1 $^{Cre/+}$; Nipbl $^{Flox/+}$; Tomato GFP/+ | NO DEFECT | Light Sheet |

| | | | | | |
|---|---|---|---|---|---|
| E17.5 | 3 | D750-4 | Isl1 $^{Cre/+}$; Nipbl $^{Flox/+}$; Tomato GFP/+ | ASD OS | Light Sheet |
| E17.5 | 4 | D750-6 | Isl1 $^{Cre/+}$; Nipbl $^{Flox/+}$; Tomato GFP/+ | NO DEFECT | Light Sheet |
| E17.5 | 5 | D750-8 | Isl1 $^{Cre/+}$; Nipbl $^{Flox/+}$; Tomato GFP/+ | ASD OS | Light Sheet |
| E17.5 | 6 | D773-34 | Isl1 $^{Cre/+}$; Nipbl $^{Flox/+}$; Tomato GFP/+ | NO DEFECT | Light Sheet |
| E17.5 | 7 | D773-36 | Isl1 $^{Cre/+}$; Nipbl $^{Flox/+}$; Tomato GFP/+ | NO DEFECT | Light Sheet |
| E17.5 | 8 | D773-37 | Isl1 $^{Cre/+}$; Nipbl $^{Flox/+}$; Tomato GFP/+ | NO DEFECT | Light Sheet |
| E17.5 | 9 | D773-38 | Isl1 $^{Cre/+}$; Nipbl $^{Flox/+}$; Tomato GFP/+ | NO DEFECT | Light Sheet |
| E17.5 | 10 | D777-39 | Isl1 $^{Cre/+}$; Nipbl $^{Flox/+}$; Tomato GFP/+ | NO DEFECT | Light Sheet |
| E17.5 | 11 | D777-41 | Isl1 $^{Cre/+}$; Nipbl $^{Flox/+}$; Tomato GFP/+ | ASD OS | Light Sheet |
| E17.5 | 12 | D777-43 | Isl1 $^{Cre/+}$; Nipbl $^{Flox/+}$; Tomato GFP/+ | ASD OP | Light Sheet |
| E17.5 | 13 | D777-45 | Isl1 $^{Cre/+}$; Nipbl $^{Flox/+}$; Tomato GFP/+ | ASD OS | Light Sheet |
| E17.5 | 14 | D778-49 | Isl1 $^{Cre/+}$; Nipbl $^{Flox/+}$; Tomato GFP/+ | ASD OS | Light Sheet |
| E17.5 | 15 | D778-51 | Isl1 $^{Cre/+}$; Nipbl $^{Flox/+}$; Tomato GFP/+ | NO DEFECT | Light Sheet |
| E17.5 | 16 | D778-52 | Isl1 $^{Cre/+}$; Nipbl $^{Flox/+}$; Tomato GFP/+ | NO DEFECT | Light Sheet |
| E17.5 | 17 | D779-56 | Isl1 $^{Cre/+}$; Nipbl $^{Flox/+}$; Tomato GFP/+ | VSD;OA | Light Sheet |
| E17.5 | 18 | D751-10 | Isl1 $^{Cre/+}$; Nipbl $^{Flox/+}$; Tomato GFP/+ | NO DEFECT | Light Sheet |
| E17.5 | 19 | D751-11 | Isl1 $^{Cre/+}$; Nipbl $^{Flox/+}$; Tomato GFP/+ | NO DEFECT | Light Sheet |
| E17.5 | 20 | D751-12 | Isl1 $^{Cre/+}$; Nipbl $^{Flox/+}$; Tomato GFP/+ | VSD; OA | Light Sheet |
| E17.5 | 21 | D751-13 | Isl1 $^{Cre/+}$; Nipbl $^{Flox/+}$; Tomato GFP/+ | NO DEFECT | Light Sheet |
| E17.5 | 22 | D751-14 | Isl1 $^{Cre/+}$; Nipbl $^{Flox/+}$; Tomato GFP/+ | VSD; PTA | Light Sheet |
| E17.5 | 23 | D729-9 | Isl1 $^{Cre/+}$; Nipbl $^{Flox/+}$; Tomato GFP/+ | VSD | Section |
| E17.5 | 24 | D729-10 | Isl1 $^{Cre/+}$; Nipbl $^{Flox/+}$; Tomato GFP/+ | NO DEFECT | Light Sheet |
| E17.5 | 25 | D729-13 | Isl1 $^{Cre/+}$; Nipbl $^{Flox/+}$; Tomato GFP/+ | NO DEFECT | Light Sheet |
| E17.5 | 26 | D729-14 | Isl1 $^{Cre/+}$; Nipbl $^{Flox/+}$; Tomato GFP/+ | NO DEFECT | Section |
| E17.5 | 27 | D729-15 | Isl1 $^{Cre/+}$; Nipbl $^{Flox/+}$; Tomato GFP/+ | NO DEFECT | Section |
| E17.5 | 28 | D728-1 | Isl1 $^{Cre/+}$; Nipbl $^{Flox/+}$; Tomato GFP/+ | NO DEFECT | Light Sheet |
| E17.5 | 29 | D728-4 | Isl1 $^{Cre/+}$; Nipbl $^{Flox/+}$; Tomato GFP/+ | NO DEFECT | Section |
| E17.5 | 30 | D728-5 | Isl1 $^{Cre/+}$; Nipbl $^{Flox/+}$; Tomato GFP/+ | NO DEFECT | Section |
| E17.5 | 31 | D730-9 | Isl1 $^{Cre/+}$; Nipbl $^{Flox/+}$; Tomato GFP/+ | NO DEFECT | Light Sheet |

| | | | | | |
|---|---|---|---|---|---|
| E17.5 | 32 | D730-11 | Isl1 $^{Cre/+}$; Nipbl $^{Flox/+}$; Tomato GFP/+ | NO DEFECT | Light Sheet |
| E17.5 | 33 | D730-12 | Isl1 $^{Cre/+}$; Nipbl $^{Flox/+}$; Tomato GFP/+ | NO DEFECT | Section |
| E17.5 | 34 | D730-14 | Isl1 $^{Cre/+}$; Nipbl $^{Flox/+}$; Tomato GFP/+ | NO DEFECT | Light Sheet |
| E17.5 | 35 | D730-15 | Isl1 $^{Cre/+}$; Nipbl $^{Flox/+}$; Tomato GFP/+ | NO DEFECT | Section |
| E17.5 | 36 | D730-16 | Isl1 $^{Cre/+}$; Nipbl $^{Flox/+}$; Tomato GFP/+ | NO DEFECT | Light Sheet |
| E17.5 | 37 | D757-19 | Isl1 $^{Cre/+}$; Nipbl $^{Flox/+}$; Tomato GFP/+ | NO DEFECT | Light Sheet |
| E17.5 | 38 | D757-21 | Isl1 $^{Cre/+}$; Nipbl $^{Flox/+}$; Tomato GFP/+ | NO DEFECT | Light Sheet |
| E17.5 | 39 | D759-25 | Isl1 $^{Cre/+}$; Nipbl $^{Flox/+}$; Tomato GFP/+ | NO DEFECT | Light Sheet |
| E17.5 | 40 | D759-26 | Isl1 $^{Cre/+}$; Nipbl $^{Flox/+}$; Tomato GFP/+ | NO DEFECT | Light Sheet |
| E17.5 | 41 | D762-30 | Isl1 $^{Cre/+}$; Nipbl $^{Flox/+}$; Tomato GFP/+ | NO DEFECT | Light Sheet |
| E17.5 | 42 | D762-31 | Isl1 $^{Cre/+}$; Nipbl $^{Flox/+}$; Tomato GFP/+ | NO DEFECT | Light Sheet |
| E17.5 | 43 | D726-1 | Isl1 $^{Cre/+}$; Nipbl $^{Flox/+}$; Tomato GFP/+ | VSD | Section |
| E17.5 | 44 | D726-3 | Isl1 $^{Cre/+}$; Nipbl $^{Flox/+}$; Tomato GFP/+ | ASD; VSD | Section |
| E17.5 | 45 | D726-5 | Isl1 $^{Cre/+}$; Nipbl $^{Flox/+}$; Tomato GFP/+ | NO DEFECT | Light Sheet |
| E17.5 | 46 | D726-7 | Isl1 $^{Cre/+}$; Nipbl $^{Flox/+}$; Tomato GFP/+ | NO DEFECT | Light Sheet |
| E17.5 | 47 | D731-20 | Isl1 $^{Cre/+}$; Nipbl $^{Flox/+}$ | NO DEFECT | Section |
| E17.5 | 48 | D731-23 | Isl1 $^{Cre/+}$; Nipbl $^{Flox/+}$ | NO DEFECT | Light Sheet |
| E17.5 | 49 | D732-24 | Isl1 $^{Cre/+}$; Nipbl $^{Flox/+}$ | NO DEFECT | Section |
| E17.5 | 50 | D732-28 | Isl1 $^{Cre/+}$; Nipbl $^{Flox/+}$ | NO DEFECT | Light Sheet |
| E17.5 | 51 | D732-30 | Isl1 $^{Cre/+}$; Nipbl $^{Flox/+}$ | NO DEFECT | Light Sheet |

**Table S8.** Ventricular volumes of hearts analyzed in Fig. 2F.

| Age | No | Sample | Genotype | Ventricular Volume ($mm^3$) |
|---|---|---|---|---|
| E17.5 | 1 | D750-1 | Isl1 $^{Cre/+}$; Nipbl $^{Flox/+}$; Tomato GFP/+ | 6.96 |
| E17.5 | 2 | D750-2 | Isl1 $^{Cre/+}$; Nipbl $^{Flox/+}$; Tomato GFP/+ | 8.84 |
| E17.5 | 3 | D750-4 | Isl1 $^{Cre/+}$; Nipbl $^{Flox/+}$; Tomato GFP/+ | 8.57 |
| E17.5 | 4 | D750-6 | Isl1 $^{Cre/+}$; Nipbl $^{Flox/+}$; Tomato GFP/+ | 8.85 |
| E17.5 | 5 | D757-19 | Isl1 $^{Cre/+}$; Nipbl $^{Flox/+}$; Tomato GFP/+ | 6.71 |
| E17.5 | 6 | D757-21 | Isl1 $^{Cre/+}$; Nipbl $^{Flox/+}$; Tomato GFP/+ | 7.55 |
| E17.5 | 7 | D759-25 | Isl1 $^{Cre/+}$; Nipbl $^{Flox/+}$; Tomato GFP/+ | 6.95 |
| E17.5 | 8 | D759-26 | Isl1 $^{Cre/+}$; Nipbl $^{Flox/+}$; Tomato GFP/+ | 6.22 |
| E17.5 | 9 | D762-30 | Isl1 $^{Cre/+}$; Nipbl $^{Flox/+}$; Tomato GFP/+ | 9.25 |
| E17.5 | 10 | D762-31 | Isl1 $^{Cre/+}$; Nipbl $^{Flox/+}$; Tomato GFP/+ | 8.3 |
| E17.5 | 11 | D773-34 | Isl1 $^{Cre/+}$; Nipbl $^{Flox/+}$; Tomato GFP/+ | 7.13 |
| E17.5 | 12 | D773-36 | Isl1 $^{Cre/+}$; Nipbl $^{Flox/+}$; Tomato GFP/+ | 8 |
| E17.5 | 13 | D773-37 | Isl1 $^{Cre/+}$; Nipbl $^{Flox/+}$; Tomato GFP/+ | 6.68 |
| E17.5 | 14 | D773-38 | Isl1 $^{Cre/+}$; Nipbl $^{Flox/+}$; Tomato GFP/+ | 7.18 |
| E17.5 | 1 | D728-2 | Nipbl $^{Flox/+}$; Tomato GFP/+ | 8.18 |
| E17.5 | 2 | D728-3 | Nipbl $^{Flox/+}$; Tomato GFP/+ | 8.08 |
| E17.5 | 3 | D728-6 | Nipbl $^{Flox/+}$; Tomato GFP/+ | 9.02 |
| E17.5 | 4 | D728-7 | Nipbl $^{Flox/+}$; Tomato GFP/+ | 7.32 |
| E17.5 | 5 | D728-8 | Nipbl $^{Flox/+}$; Tomato GFP/+ | 8.4 |
| E17.5 | 6 | D750-3 | Nipbl $^{Flox/+}$; Tomato GFP/+ | 8.71 |
| E17.5 | 7 | D750-5 | Nipbl $^{Flox/+}$; Tomato GFP/+ | 8.76 |
| E17.5 | 8 | D750-7 | Nipbl $^{Flox/+}$; Tomato GFP/+ | 7.93 |
| E17.5 | 9 | D757-15 | Nipbl $^{Flox/+}$; Tomato GFP/+ | 6.2 |
| E17.5 | 10 | D757-16 | Nipbl $^{Flox/+}$; Tomato GFP/+ | 7.54 |
| E17.5 | 11 | D757-17 | Nipbl $^{Flox/+}$; Tomato GFP/+ | 7.59 |
| E17.5 | 12 | D757-18 | Nipbl $^{Flox/+}$; Tomato GFP/+ | 6.86 |
| E17.5 | 13 | D757-22 | Nipbl $^{Flox/+}$; Tomato GFP/+ | 7.38 |

| | | | | |
|---|---|---|---|---|
| E17.5 | 14 | D759-23 | Nipbl $^{Flox/+}$; Tomato GFP/+ | 6.81 |
| E17.5 | 15 | D759-24 | Nipbl $^{Flox/+}$; Tomato GFP/+ | 5.91 |
| E17.5 | 16 | D759-27 | Nipbl $^{Flox/+}$; Tomato GFP/+ | 6.56 |
| E17.5 | 17 | D759-28 | Nipbl $^{Flox/+}$; Tomato GFP/+ | 6.12 |
| E17.5 | 18 | D762-29 | Nipbl $^{Flox/+}$; Tomato GFP/+ | 10.3 |
| E17.5 | 19 | D762-32 | Nipbl $^{Flox/+}$; Tomato GFP/+ | 8.83 |
| E17.5 | 20 | D773-33 | Nipbl $^{Flox/+}$; Tomato GFP/+ | 8.19 |
| E17.5 | 21 | D773-35 | Nipbl $^{Flox/+}$; Tomato GFP/+ | 8.3 |

**Table S9.** Defects observed in hearts analyzed in Fig. 4B. ASD OS = Atrial septal defect ostium secundum type. Light sheet images were analyzed for all.

| Age | No | Sample | Genotype | Outcome |
|---|---|---|---|---|
| E17.5 | 1 | D736-3 | $Isl1^{+/+}$ | NO DEFECT |
| E17.5 | 2 | D736-4 | $Isl1^{+/+}$ | NO DEFECT |
| E17.5 | 3 | D736-5 | $Isl1^{+/+}$ | NO DEFECT |
| E17.5 | 4 | D736-7 | $Isl1^{+/+}$ | NO DEFECT |
| E17.5 | 5 | D736-8 | $Isl1^{+/+}$ | NO DEFECT |
| E17.5 | 6 | D736-9 | $Isl1^{+/+}$ | NO DEFECT |
| E17.5 | 7 | D738-10 | $Isl1^{+/+}$ | NO DEFECT |
| E17.5 | 8 | D738-11 | $Isl1^{+/+}$ | NO DEFECT |
| E17.5 | 9 | D738-14 | $Isl1^{+/+}$ | NO DEFECT |
| E17.5 | 10 | D738-17 | $Isl1^{+/+}$ | NO DEFECT |
| E17.5 | 11 | D738-18 | $Isl1^{+/+}$ | NO DEFECT |
| E17.5 | 12 | D738-20 | $Isl1^{+/+}$ | NO DEFECT |
| E17.5 | 13 | D739-21 | $Isl1^{+/+}$ | NO DEFECT |
| E17.5 | 14 | D739-22 | $Isl1^{+/+}$ | NO DEFECT |
| E17.5 | 15 | D739-23 | $Isl1^{+/+}$ | NO DEFECT |
| E17.5 | 16 | D739-24 | $Isl1^{+/+}$ | NO DEFECT |
| E17.5 | 17 | D739-29 | $Isl1^{+/+}$ | NO DEFECT |
| E17.5 | 18 | D739-30 | $Isl1^{+/+}$ | NO DEFECT |
| E17.5 | 19 | D740-31 | $Isl1^{+/+}$ | NO DEFECT |
| E17.5 | 20 | D740-32 | $Isl1^{+/+}$ | NO DEFECT |
| E17.5 | 21 | D740-35 | $Isl1^{+/+}$ | NO DEFECT |
| E17.5 | 22 | D740-38 | $Isl1^{+/+}$ | NO DEFECT |
| E17.5 | 23 | D740-40 | $Isl1^{+/+}$ | NO DEFECT |
| E17.5 | 24 | D744-43 | $Isl1^{+/+}$ | NO DEFECT |
| E17.5 | 25 | D744-44 | $Isl1^{+/+}$ | NO DEFECT |
| E17.5 | 26 | D744-47 | $Isl1^{+/+}$ | NO DEFECT |

| | | | | |
|---|---|---|---|---|
| E17.5 | 27 | D744-50 | Isl1 $^{+/+}$ | NO DEFECT |
| E17.5 | 28 | D763-51 | Isl1 $^{+/+}$; TomatoGFP/+ | NO DEFECT |
| E17.5 | 29 | D763-52 | Isl1 $^{+/+}$; TomatoGFP/+ | NO DEFECT |
| E17.5 | 30 | D763-53 | Isl1 $^{+/+}$; TomatoGFP/+ | ASD OS |
| E17.5 | 31 | D763-54 | Isl1 $^{+/+}$; TomatoGFP/+ | NO DEFECT |
| E17.5 | 32 | D765-62 | Isl1 $^{+/+}$; TomatoGFP/+ | NO DEFECT |
| E17.5 | 33 | D765-63 | Isl1 $^{+/+}$; TomatoGFP/+ | NO DEFECT |
| E17.5 | 34 | D765-64 | Isl1 $^{+/+}$; TomatoGFP/+ | NO DEFECT |
| E17.5 | 35 | D765-65 | Isl1 $^{+/+}$; TomatoGFP/+ | NO DEFECT |
| E17.5 | 36 | D765-66 | Isl1 $^{+/+}$; TomatoGFP/+ | NO DEFECT |
| E17.5 | 37 | D766-70 | Isl1 $^{+/+}$; TomatoGFP/+ | NO DEFECT |
| E17.5 | 38 | D766-71 | Isl1 $^{+/+}$; TomatoGFP/+ | NO DEFECT |
| E17.5 | 39 | D766-75 | Isl1 $^{+/+}$; TomatoGFP/+ | NO DEFECT |
| E17.5 | 40 | D766-76 | Isl1 $^{+/+}$; TomatoGFP/+ | NO DEFECT |
| E17.5 | 41 | D767-77 | Isl1 $^{+/+}$; TomatoGFP/+ | NO DEFECT |
| E17.5 | 42 | D767-79 | Isl1 $^{+/+}$; TomatoGFP/+ | NO DEFECT |
| E17.5 | 43 | D767-82 | Isl1 $^{+/+}$; TomatoGFP/+ | NO DEFECT |
| E17.5 | 44 | D771-87 | Isl1 $^{+/+}$; TomatoGFP/+ | NO DEFECT |
| E17.5 | 45 | D771-89 | Isl1 $^{+/+}$; TomatoGFP/+ | NO DEFECT |
| E17.5 | 46 | D772-93 | Isl1 $^{+/+}$; TomatoGFP/+ | NO DEFECT |
| E17.5 | 47 | D772-95 | Isl1 $^{+/+}$; TomatoGFP/+ | NO DEFECT |
| E17.5 | 48 | D772-96 | Isl1 $^{+/+}$; TomatoGFP/+ | NO DEFECT |
| E17.5 | 49 | D772-98 | Isl1 $^{+/+}$; TomatoGFP/+ | NO DEFECT |
| E17.5 | 50 | D772-99 | Isl1 $^{+/+}$; TomatoGFP/+ | NO DEFECT |
| E17.5 | 1 | D736-1 | Isl1 $^{Cre/+}$ | ASD OS |
| E17.5 | 2 | D736-6 | Isl1 $^{Cre/+}$ | ASD OS |
| E17.5 | 3 | D738-12 | Isl1 $^{Cre/+}$ | NO DEFECT |
| E17.5 | 4 | D738-13 | Isl1 $^{Cre/+}$ | NO DEFECT |
| E17.5 | 5 | D738-15 | Isl1 $^{Cre/+}$ | NO DEFECT |

| | | | | |
|---|---|---|---|---|
| E17.5 | 6 | D738-16 | Isl1 $^{Cre/+}$ | NO DEFECT |
| E17.5 | 7 | D738-19 | Isl1 $^{Cre/+}$ | NO DEFECT |
| E17.5 | 8 | D739-25 | Isl1 $^{Cre/+}$ | NO DEFECT |
| E17.5 | 9 | D739-26 | Isl1 $^{Cre/+}$ | ASD OS |
| E17.5 | 10 | D739-27 | Isl1 $^{Cre/+}$ | NO DEFECT |
| E17.5 | 11 | D739-28 | Isl1 $^{Cre/+}$ | NO DEFECT |
| E17.5 | 12 | D740-33 | Isl1 $^{Cre/+}$ | NO DEFECT |
| E17.5 | 13 | D740-34 | Isl1 $^{Cre/+}$ | NO DEFECT |
| E17.5 | 14 | D740-36 | Isl1 $^{Cre/+}$ | ASD OS |
| E17.5 | 15 | D740-39 | Isl1 $^{Cre/+}$ | NO DEFECT |
| E17.5 | 16 | D740-41 | Isl1 $^{Cre/+}$ | NO DEFECT |
| E17.5 | 17 | D744-42 | Isl1 $^{Cre/+}$ | NO DEFECT |
| E17.5 | 18 | D744-45 | Isl1 $^{Cre/+}$ | NO DEFECT |
| E17.5 | 19 | D744-46 | Isl1 $^{Cre/+}$ | NO DEFECT |
| E17.5 | 20 | D744-48 | Isl1 $^{Cre/+}$ | NO DEFECT |
| E17.5 | 21 | D744-49 | Isl1 $^{Cre/+}$ | NO DEFECT |
| E17.5 | 22 | D763-55 | Isl1 $^{Cre/+}$; TomatoGFP/+ | ASD OS |
| E17.5 | 23 | D765-57 | Isl1 $^{Cre/+}$; TomatoGFP/+ | NO DEFECT |
| E17.5 | 24 | D765-58 | Isl1 $^{Cre/+}$; TomatoGFP/+ | NO DEFECT |
| E17.5 | 25 | D765-59 | Isl1 $^{Cre/+}$; TomatoGFP/+ | ASD OS |
| E17.5 | 26 | D765-60 | Isl1 $^{Cre/+}$; TomatoGFP/+ | ASD OS |
| E17.5 | 27 | D765-61 | Isl1 $^{Cre/+}$; TomatoGFP/+ | ASD OS |
| E17.5 | 28 | D766-69 | Isl1 $^{Cre/+}$; TomatoGFP/+ | NO DEFECT |
| E17.5 | 29 | D766-73 | Isl1 $^{Cre/+}$; TomatoGFP/+ | NO DEFECT |
| E17.5 | 30 | D766-74 | Isl1 $^{Cre/+}$; TomatoGFP/+ | NO DEFECT |
| E17.5 | 31 | D767-80 | Isl1 $^{Cre/+}$; TomatoGFP/+ | NO DEFECT |
| E17.5 | 32 | D767-81 | Isl1 $^{Cre/+}$; TomatoGFP/+ | NO DEFECT |
| E17.5 | 33 | D767-84 | Isl1 $^{Cre/+}$; TomatoGFP/+ | NO DEFECT |
| E17.5 | 34 | D767-85 | Isl1 $^{Cre/+}$; TomatoGFP/+ | ASD OS |

| | | | | |
|---|---|---|---|---|
| E17.5 | 35 | D767-86 | Isl1 $^{Cre/+}$; TomatoGFP/+ | NO DEFECT |
| E17.5 | 36 | D771-88 | Isl1 $^{Cre/+}$; TomatoGFP/+ | NO DEFECT |
| E17.5 | 37 | D771-90 | Isl1 $^{Cre/+}$; TomatoGFP/+ | NO DEFECT |
| E17.5 | 38 | D771-91 | Isl1 $^{Cre/+}$; TomatoGFP/+ | NO DEFECT |
| E17.5 | 39 | D771-92 | Isl1 $^{Cre/+}$; TomatoGFP/+ | NO DEFECT |
| E17.5 | 40 | D772-94 | Isl1 $^{Cre/+}$; TomatoGFP/+ | NO DEFECT |
| E17.5 | 41 | D772-97 | Isl1 $^{Cre/+}$; TomatoGFP/+ | NO DEFECT |

**Table S10.** Ventricular volumes of hearts analyzed in Fig. 4C.

| Age | No | Sample | Genotype | Ventricular Volume ($mm^3$) |
|---|---|---|---|---|
| E17.5 | 1 | D765-62 | Isl1 $^{+/+}$; TomatoGFP/+ | 9.46 |
| E17.5 | 2 | D765-63 | Isl1 $^{+/+}$; TomatoGFP/+ | 9.53 |
| E17.5 | 3 | D765-64 | Isl1 $^{+/+}$; TomatoGFP/+ | 9.69 |
| E17.5 | 4 | D765-65 | Isl1 $^{+/+}$; TomatoGFP/+ | 9.75 |
| E17.5 | 5 | D766-67 | Isl1 $^{+/+}$; TomatoGFP/+ | 6.46 |
| E17.5 | 6 | D766-70 | Isl1 $^{+/+}$; TomatoGFP/+ | 7.42 |
| E17.5 | 7 | D766-71 | Isl1 $^{+/+}$; TomatoGFP/+ | 6.28 |
| E17.5 | 8 | D766-76 | Isl1 $^{+/+}$; TomatoGFP/+ | 7.29 |
| E17.5 | 9 | D767-77 | Isl1 $^{+/+}$; TomatoGFP/+ | 7.31 |
| E17.5 | 10 | D767-78 | Isl1 $^{+/+}$; TomatoGFP/+ | 6.33 |
| E17.5 | 11 | D767-79 | Isl1 $^{+/+}$; TomatoGFP/+ | 6.87 |
| E17.5 | 12 | D767-82 | Isl1 $^{+/+}$; TomatoGFP/+ | 7.89 |
| E17.5 | 13 | D772-93 | Isl1 $^{+/+}$; TomatoGFP/+ | 7.56 |
| E17.5 | 14 | D772-95 | Isl1 $^{+/+}$; TomatoGFP/+ | 8.05 |
| E17.5 | 15 | D772-96 | Isl1 $^{+/+}$; TomatoGFP/+ | 5.71 |
| E17.5 | 16 | D772-98 | Isl1 $^{+/+}$; TomatoGFP/+ | 9.09 |
| E17.5 | 17 | D772-99 | Isl1 $^{+/+}$; TomatoGFP/+ | 8.88 |
| E17.5 | 1 | D765-58 | Isl1 $^{Cre/+}$; TomatoGFP/+ | 7.71 |
| E17.5 | 2 | D765-59 | Isl1 $^{Cre/+}$; TomatoGFP/+ | 9.14 |
| E17.5 | 3 | D765-60 | Isl1 $^{Cre/+}$; TomatoGFP/+ | 8.55 |
| E17.5 | 4 | D765-61 | Isl1 $^{Cre/+}$; TomatoGFP/+ | 8.95 |
| E17.5 | 5 | D766-68 | Isl1 $^{Cre/+}$; TomatoGFP/+ | 6.33 |
| E17.5 | 6 | D766-69 | Isl1 $^{Cre/+}$; TomatoGFP/+ | 6.05 |
| E17.5 | 7 | D766-72 | Isl1 $^{Cre/+}$; TomatoGFP/+ | 7.01 |
| E17.5 | 8 | D766-73 | Isl1 $^{Cre/+}$; TomatoGFP/+ | 7.11 |
| E17.5 | 9 | D766-74 | Isl1 $^{Cre/+}$; TomatoGFP/+ | 9.04 |
| E17.5 | 10 | D767-80 | Isl1 $^{Cre/+}$; TomatoGFP/+ | 6.74 |

| | | | | |
|---|---|---|---|---|
| E17.5 | 11 | D767-81 | Isl1 $^{Cre/+}$; TomatoGFP/+ | 9.83 |
| E17.5 | 12 | D767-84 | Isl1 $^{Cre/+}$; TomatoGFP/+ | 7.6 |
| E17.5 | 13 | D767-85 | Isl1 $^{Cre/+}$; TomatoGFP/+ | 7.15 |
| E17.5 | 14 | D767-86 | Isl1 $^{Cre/+}$; TomatoGFP/+ | 6.83 |
| E17.5 | 15 | D772-94 | Isl1 $^{Cre/+}$; TomatoGFP/+ | 7.12 |
| E17.5 | 16 | D772-97 | Isl1 $^{Cre/+}$; TomatoGFP/+ | 6.62 |

**Table S11.** qRT-PCR data for *Isl1* in hearts analyzed in Fig. 4D. Strikethrough indicates that the value was not used for calculating the mean and standard deviation.

| | | | Isl1 Exon1-Exon2 | | | | Rpl4 | | | | |
|---|---|---|---|---|---|---|---|---|---|---|---|
| **Age** | **Genotype** | **Sample** | **Well** | **Ct** | **Mean Ct** | **SD Ct** | **Well** | **Ct** | **Mean Ct** | **SD Ct** | **dCt (Isl1 - Rpl4)** |
| E10.5 | *Isl1 +/+* | d795-1 | A01 | 30.24 | 30.20 | 0.14 | A01 | 20.15 | 19.98 | 0.15 | 10.22 |
| | | | A02 | 30.04 | | | A02 | 19.95 | | | |
| | | | A03 | 30.31 | | | A03 | 19.85 | | | |
| | | d795-2 | A04 | 27.12 | 27.20 | 0.08 | A04 | 19.50 | 19.48 | 0.07 | 7.72 |
| | | | A05 | 27.28 | | | A05 | 19.53 | | | |
| | | | A06 | 27.21 | | | A06 | 19.40 | | | |
| | | d796-2 | B04 | 27.10 | 27.18 | 0.09 | B04 | 19.34 | 19.37 | 0.04 | 7.81 |
| | | | B05 | 27.16 | | | B05 | 19.41 | | | |
| | | | B06 | 27.27 | | | B06 | 19.34 | | | |
| | | d796-6 | B07 | 26.63 | 26.60 | 0.03 | B07 | 19.47 | 19.38 | 0.09 | 7.22 |
| | | | B08 | 26.60 | | | B08 | 19.37 | | | |
| | | | B09 | 26.58 | | | B09 | 19.30 | | | |
| | | d796-9 | C04 | 27.76 | 27.89 | 0.12 | C04 | 19.29 | 19.22 | 0.06 | 8.67 |
| | | | C05 | 27.99 | | | C05 | 19.18 | | | |
| | | | C06 | 27.93 | | | C06 | 19.18 | | | |
| | | d808-2 | C10 | 27.93 | 28.04 | 0.12 | C10 | 19.31 | 19.42 | 0.10 | 8.62 |
| | | | C11 | 28.17 | | | C11 | 19.46 | | | |
| | | | C12 | 28.02 | | | C12 | 19.50 | | | |
| | | d808-3 | D01 | 27.68 | 27.76 | 0.10 | D01 | 19.62 | 19.48 | 0.13 | 8.28 |
| | | | D02 | 27.87 | | | D02 | 19.40 | | | |
| | | | D03 | 27.73 | | | D03 | 19.41 | | | |
| | | d808-5 | D04 | 27.48 | 27.57 | 0.09 | D04 | 18.96 | 18.95 | 0.04 | 8.62 |
| | | | D05 | 27.66 | | | D05 | 18.99 | | | |
| | | | D06 | 27.58 | | | D06 | 18.90 | | | |
| | | d808-8 | E01 | 27.82 | 27.65 | 0.17 | E01 | 19.23 | 19.16 | 0.07 | 8.49 |

| | | | | | | | | | | | |
|---|---|---|---|---|---|---|---|---|---|---|---|
| | | | E02 | 27.65 | | | E02 | 19.13 | | | |
| | | | E03 | 27.47 | | | E03 | 19.10 | | | |
| | | | A07 | 29.08 | | | A07 | 19.58 | | | |
| | | d795-3 | A08 | ~~28.74~~ | 29.05 | 0.05 | A08 | 19.53 | 19.53 | 0.06 | 9.52 |
| | | | A09 | 29.01 | | | A09 | 19.46 | | | |
| | | | A10 | 30.98 | | | A10 | 19.97 | | | |
| | | d795-9 | A11 | 31.39 | 31.41 | 0.29 | A11 | 19.83 | 19.95 | 0.11 | 11.46 |
| | | | A12 | ~~31.84~~ | | | A12 | 20.05 | | | |
| | | | B01 | 30.62 | | | B01 | 19.40 | | | |
| | | d796-1 | B02 | 30.57 | 30.89 | 0.04 | B02 | 19.29 | 19.30 | 0.10 | 11.59 |
| | | | B03 | ~~31.15~~ | | | B03 | 19.20 | | | |
| | | | B10 | 29.12 | | | B10 | 19.41 | | | |
| | | d796-7 | B11 | 29.15 | 29.16 | 0.05 | B11 | 19.49 | 19.48 | 0.07 | 9.68 |
| | | | B12 | 29.21 | | | B12 | 19.54 | | | |
| | | | C01 | 30.26 | | | C01 | 19.20 | | | |
| | *Isl1* $^{Cre/+}$ | d796-8 | C02 | ~~30.49~~ | 30.26 | 0.01 | C02 | 19.16 | 19.15 | 0.06 | 11.11 |
| | | | C03 | 30.25 | | | C03 | 19.08 | | | |
| | | | C07 | ~~28.23~~ | | | C07 | 19.31 | | | |
| | | d808-1 | C08 | 28.59 | 28.66 | 0.09 | C08 | 19.29 | 19.29 | 0.02 | 9.37 |
| | | | C09 | 28.72 | | | C09 | 19.27 | | | |
| | | | D07 | ~~28.61~~ | | | D07 | ~~19.47~~ | | | |
| | | d808-6 | D08 | 29.00 | 28.99 | 0.01 | D08 | 19.20 | 19.16 | 0.06 | 9.84 |
| | | | D09 | 28.98 | | | D09 | 19.11 | | | |
| | | | D10 | 28.43 | | | D10 | 19.17 | | | |
| | | d808-7 | D11 | 28.68 | 28.56 | 0.18 | D11 | 19.15 | 19.16 | 0.01 | 9.40 |
| | | | D12 | ~~29.11~~ | | | D12 | 19.15 | | | |
| | | | E04 | 28.81 | | | E04 | 19.05 | | | |
| | | d808-9 | E05 | 29.01 | 28.89 | 0.11 | E05 | 19.04 | 19.00 | 0.08 | 9.89 |
| | | | E06 | 28.85 | | | E06 | 18.91 | | | |

**Table S12.** Comparative Ct analysis of Isl1 in hearts shown in Fig. 4D. SEM = Standard error of the mean.

| | Mean dCt | ddCt | $2^{-ddCt}$ | SEM | Range Lower SEM | Range Upper SEM |
|---|---|---|---|---|---|---|
| Isl1 +/+ | 8.41 | 0.00 | 1.00 | 0.28 | 1.22 | 0.82 |
| Isl1 Cre/+ | 10.20 | 1.80 | 0.29 | 0.30 | 0.35 | 0.23 |

**Table S13.** Sample identifiers of embryos shown in Fig. 5B.

| Age | Panel | Sample | Genotype |
|---|---|---|---|
| E17.5 | 1 | d737-3 | *Nipbl* $^{+/+}$; *Isl1* $^{+/+}$ |
| E17.5 | 2 | d737-6 | *Isl1* $^{Cre/+}$ |
| E17.5 | 3 | d737-1 | *Nipbl* $^{+/-}$ |
| E17.5 | 4 | d737-2 | *Nipbl* $^{+/-}$; *Isl1* $^{Cre/+}$ |

**Table S14.** Sample identifiers of embryos shown in Fig. 5C.

| Age | Panel | Sample | Genotype |
|---|---|---|---|
| E15.5 | 1 | EZW16-5 | *Nipbl* $^{+/+}$; *Isl1* $^{+/+}$ |
| E15.5 | 2 | EZW16-4 | *Isl1* $^{Cre/+}$ |
| E15.5 | 3 | EYP6-8 | *Nipbl* $^{+/-}$ |
| E15.5 | 4 | EYP6-9 | *Nipbl* $^{+/-}$; *Isl1* $^{Cre/+}$ |

**Table S15.** Signs of fetal demise observed in embryos analyzed in Fig. 5D.

| Age | No | Sample | Genotype | Sign(s) of Fetal Demise |
|---|---|---|---|---|
| E17.5 | 1 | d733-12 | *Nipbl* $^{+/+}$; *Isl1* $^{+/+}$ | None |
| E17.5 | 2 | d737-3 | *Nipbl* $^{+/+}$; *Isl1* $^{+/+}$ | None |
| E17.5 | 1 | d727-2 | *Isl1* $^{Cre/+}$ | None |
| E17.5 | 2 | d727-3 | *Isl1* $^{Cre/+}$ | None |
| E17.5 | 3 | d727-4 | *Isl1* $^{Cre/+}$ | None |
| E17.5 | 4 | d727-5 | *Isl1* $^{Cre/+}$ | None |
| E17.5 | 5 | d727-6 | *Isl1* $^{Cre/+}$ | None |
| E17.5 | 6 | d733-8 | *Isl1* $^{Cre/+}$ | None |
| E17.5 | 7 | d733-14 | *Isl1* $^{Cre/+}$ | None |
| E17.5 | 8 | d737-6 | *Isl1* $^{Cre/+}$ | None |
| E17.5 | 9 | d737-7 | *Isl1* $^{Cre/+}$ | None |
| E17.5 | 1 | d733-7 | *Nipbl* $^{+/-}$ | None |
| E17.5 | 2 | d733-11 | *Nipbl* $^{+/-}$ | None |
| E17.5 | 3 | d737-1 | *Nipbl* $^{+/-}$ | None |
| E17.5 | 4 | d737-4 | *Nipbl* $^{+/-}$ | None |
| E17.5 | 5 | d737-5 | *Nipbl* $^{+/-}$ | None |
| E17.5 | 1 | d727-1 | *Nipbl* $^{+/-}$; *Isl1* $^{Cre/+}$ | None |
| E17.5 | 2 | d733-9 | *Nipbl* $^{+/-}$; *Isl1* $^{Cre/+}$ | undergoing resorption |
| E17.5 | 3 | d733-10 | *Nipbl* $^{+/-}$; *Isl1* $^{Cre/+}$ | undergoing resorption |
| E17.5 | 4 | d733-13 | *Nipbl* $^{+/-}$; *Isl1* $^{Cre/+}$ | undergoing resorption |
| E17.5 | 5 | d737-2 | *Nipbl* $^{+/-}$; *Isl1* $^{Cre/+}$ | hemorrhaging, severe edema |
| E17.5 | 6 | d737-8 | *Nipbl* $^{+/-}$; *Isl1* $^{Cre/+}$ | undergoing resorption |

**Table S16.** Crown rump lengths of embryos analyzed in Fig. 5E.

| Age | No | Sample | Genotype | Crown Rump Length (mm) |
|---|---|---|---|---|
| E15.5 | 1 | EUG7-3 | *Nipbl* $^{+/+}$; *Isl1* $^{+/+}$ | 16.48 |
| E15.5 | 2 | EUH7-1 | *Nipbl* $^{+/+}$; *Isl1* $^{+/+}$ | 15.21 |
| E15.5 | 3 | EUH7-2 | *Nipbl* $^{+/+}$; *Isl1* $^{+/+}$ | 15.24 |
| E15.5 | 4 | EUH7-3 | *Nipbl* $^{+/+}$; *Isl1* $^{+/+}$ | 14.54 |
| E15.5 | 5 | EUH7-8 | *Nipbl* $^{+/+}$; *Isl1* $^{+/+}$ | 14.62 |
| E15.5 | 6 | EZW16-5 | *Nipbl* $^{+/+}$; *Isl1* $^{+/+}$ | 15.18 |
| E15.5 | 7 | FEY7-1 | *Nipbl* $^{+/+}$; *Isl1* $^{+/+}$ | 14.96 |
| E15.5 | 8 | EWH5-1 | *Nipbl* $^{+/+}$; *Isl1* $^{+/+}$ | 14.51 |
| E15.5 | 9 | EWH5-6 | *Nipbl* $^{+/+}$; *Isl1* $^{+/+}$ | 14.48 |
| E15.5 | 10 | EYP6-1 | *Nipbl* $^{+/+}$; *Isl1* $^{+/+}$ | 15.24 |
| E15.5 | 11 | EYP6-3 | *Nipbl* $^{+/+}$; *Isl1* $^{+/+}$ | 15.09 |
| E15.5 | 12 | EYP6-4 | *Nipbl* $^{+/+}$; *Isl1* $^{+/+}$ | 14.64 |
| E15.5 | 13 | EYP6-5 | *Nipbl* $^{+/+}$; *Isl1* $^{+/+}$ | 14.81 |
| E15.5 | 14 | FFA16-2 | *Nipbl* $^{+/+}$; *Isl1* $^{+/+}$ | 14.95 |
| E15.5 | 15 | FFA16-6 | *Nipbl* $^{+/+}$; *Isl1* $^{+/+}$ | 15.22 |
| E15.5 | 16 | FFA16-7 | *Nipbl* $^{+/+}$; *Isl1* $^{+/+}$ | 14.88 |
| E15.5 | 17 | FFA16-9 | *Nipbl* $^{+/+}$; *Isl1* $^{+/+}$ | 15.65 |
| E15.5 | 18 | FJP5-1 | *Nipbl* $^{+/+}$; *Isl1* $^{+/+}$ | 16.46 |
| E15.5 | 19 | FJP5-7 | *Nipbl* $^{+/+}$; *Isl1* $^{+/+}$ | 15.97 |
| E15.5 | 1 | EUG7-1 | *Isl1* $^{Cre/+}$ | 15.62 |
| E15.5 | 2 | EUG7-2 | *Isl1* $^{Cre/+}$ | 14.74 |
| E15.5 | 3 | EZW16-1 | *Isl1* $^{Cre/+}$ | 15.80 |
| E15.5 | 4 | EZW16-4 | *Isl1* $^{Cre/+}$ | 14.90 |
| E15.5 | 5 | EZW16-7 | *Isl1* $^{Cre/+}$ | 14.89 |
| E15.5 | 6 | FEY7-3 | *Isl1* $^{Cre/+}$ | 14.99 |
| E15.5 | 7 | FEY7-5 | *Isl1* $^{Cre/+}$ | 14.41 |
| E15.5 | 8 | EWH5-3 | *Isl1* $^{Cre/+}$ | 13.00 |

| | | | | |
|---|---|---|---|---|
| E15.5 | 9 | EWH5-4 | *Isl1* $^{Cre/+}$ | 14.69 |
| E15.5 | 10 | EXU5-4 | *Isl1* $^{Cre/+}$ | 13.44 |
| E15.5 | 11 | EXU5-6 | *Isl1* $^{Cre/+}$ | 13.12 |
| E15.5 | 12 | EXU5-7 | *Isl1* $^{Cre/+}$ | 12.76 |
| E15.5 | 13 | EYP6-6 | *Isl1* $^{Cre/+}$ | 14.83 |
| E15.5 | 14 | EQU8-1 | *Isl1* $^{Cre/+}$ | 15.96 |
| E15.5 | 15 | EQU8-3 | *Isl1* $^{Cre/+}$ | 15.34 |
| E15.5 | 16 | EQU8-4 | *Isl1* $^{Cre/+}$ | 15.76 |
| E15.5 | 17 | EQU8-7 | *Isl1* $^{Cre/+}$ | 14.97 |
| E15.5 | 18 | ERV5-1 | *Isl1* $^{Cre/+}$ | 15.80 |
| E15.5 | 19 | ERV5-5 | *Isl1* $^{Cre/+}$ | 16.08 |
| E15.5 | 20 | FFA16-3 | *Isl1* $^{Cre/+}$ | 14.86 |
| E15.5 | 21 | FFA16-8 | *Isl1* $^{Cre/+}$ | 14.69 |
| E15.5 | 1 | EUG7-5 | *Nipbl* $^{+/-}$ | 14.46 |
| E15.5 | 2 | EUH7-5 | *Nipbl* $^{+/-}$ | 12.49 |
| E15.5 | 3 | EUH7-6 | *Nipbl* $^{+/-}$ | 11.66 |
| E15.5 | 4 | EUH7-7 | *Nipbl* $^{+/-}$ | 11.87 |
| E15.5 | 5 | EZW16-2 | *Nipbl* $^{+/-}$ | 14.79 |
| E15.5 | 6 | FEY7-4 | *Nipbl* $^{+/-}$ | 13.22 |
| E15.5 | 7 | EWH5-2 | *Nipbl* $^{+/-}$ | 13.28 |
| E15.5 | 8 | EWH5-5 | *Nipbl* $^{+/-}$ | 12.97 |
| E15.5 | 9 | EYP6-8 | *Nipbl* $^{+/-}$ | 13.08 |
| E15.5 | 10 | EQU8-6 | *Nipbl* $^{+/-}$ | 14.20 |
| E15.5 | 11 | ERV5-2 | *Nipbl* $^{+/-}$ | 14.48 |
| E15.5 | 12 | FFA16-5 | *Nipbl* $^{+/-}$ | 14.10 |
| E15.5 | 13 | FJP5-4 | *Nipbl* $^{+/-}$ | 14.14 |
| E15.5 | 14 | FJP5-5 | *Nipbl* $^{+/-}$ | 13.45 |
| E15.5 | 1 | EUG7-4 | *Nipbl* $^{+/-}$; *Isl1* $^{Cre/+}$ | 13.82 |
| E15.5 | 2 | EZW16-3 | *Nipbl* $^{+/-}$; *Isl1* $^{Cre/+}$ | 13.62 |

| | | | | |
|---|---|---|---|---|
| E15.5 | 3 | EZW16-6 | *Nipbl* $^{+/-}$; *Isl1* $^{Cre/+}$ | 13.20 |
| E15.5 | 4 | FEY7-2 | *Nipbl* $^{+/-}$; *Isl1* $^{Cre/+}$ | 12.47 |
| E15.5 | 5 | FEY7-6 | *Nipbl* $^{+/-}$; *Isl1* $^{Cre/+}$ | 12.96 |
| E15.5 | 6 | FEY7-7 | *Nipbl* $^{+/-}$; *Isl1* $^{Cre/+}$ | 12.27 |
| E15.5 | 7 | FEY7-8 | *Nipbl* $^{+/-}$; *Isl1* $^{Cre/+}$ | 12.44 |
| E15.5 | 8 | EYP6-2 | *Nipbl* $^{+/-}$; *Isl1* $^{Cre/+}$ | 13.35 |
| E15.5 | 9 | EYP6-7 | *Nipbl* $^{+/-}$; *Isl1* $^{Cre/+}$ | 14.62 |
| E15.5 | 10 | EYP6-9 | *Nipbl* $^{+/-}$; *Isl1* $^{Cre/+}$ | 12.97 |
| E15.5 | 11 | EQU8-2 | *Nipbl* $^{+/-}$; *Isl1* $^{Cre/+}$ | 13.51 |
| E15.5 | 12 | EQU8-5 | *Nipbl* $^{+/-}$; *Isl1* $^{Cre/+}$ | 14.53 |
| E15.5 | 13 | ERV5-3 | *Nipbl* $^{+/-}$; *Isl1* $^{Cre/+}$ | 14.15 |
| E15.5 | 14 | ERV5-4 | *Nipbl* $^{+/-}$; *Isl1* $^{Cre/+}$ | 13.84 |
| E15.5 | 15 | FFA16-1 | *Nipbl* $^{+/-}$; *Isl1* $^{Cre/+}$ | 13.33 |
| E15.5 | 16 | EUH7-4 | *Nipbl* $^{+/-}$; *Isl1* $^{Cre/+}$ | 12.96 |
| E15.5 | 17 | FJP5-2 | *Nipbl* $^{+/-}$; *Isl1* $^{Cre/+}$ | 14.58 |
| E15.5 | 18 | FJP5-3 | *Nipbl* $^{+/-}$; *Isl1* $^{Cre/+}$ | 13.32 |
| E15.5 | 19 | FJP5-6 | *Nipbl* $^{+/-}$; *Isl1* $^{Cre/+}$ | 13.32 |

**Table S17.** Defects observed in hearts analyzed in Fig. 6B. ASD OP = Atrial septal defect ostium primum type, ASD OS = Atrial septal defect ostium secundum type, DORV = Double outlet right ventricle. OA = Overriding aorta, TGA = Transposition of the great arteries, PTA = Persistent truncus arteriosis, VSD = Ventricular septal defect.

| Age | No | Sample | Genotype | Outcome |
|---|---|---|---|---|
| E15.5 | 1 | EUG7-3 | *Nipbl* $^{+/+}$; *Isl1* $^{+/+}$ | NO DEFECT |
| E15.5 | 2 | EUH7-1 | *Nipbl* $^{+/+}$; *Isl1* $^{+/+}$ | NO DEFECT |
| E15.5 | 3 | EUH7-2 | *Nipbl* $^{+/+}$; *Isl1* $^{+/+}$ | NO DEFECT |
| E15.5 | 4 | EUH7-3 | *Nipbl* $^{+/+}$; *Isl1* $^{+/+}$ | NO DEFECT |
| E15.5 | 5 | EUH7-8 | *Nipbl* $^{+/+}$; *Isl1* $^{+/+}$ | NO DEFECT |
| E15.5 | 6 | EZW16-5 | *Nipbl* $^{+/+}$; *Isl1* $^{+/+}$ | NO DEFECT |
| E15.5 | 7 | EYP6-1 | *Nipbl* $^{+/+}$; *Isl1* $^{+/+}$ | NO DEFECT |
| E15.5 | 8 | EYP6-3 | *Nipbl* $^{+/+}$; *Isl1* $^{+/+}$ | NO DEFECT |
| E15.5 | 9 | EYP6-4 | *Nipbl* $^{+/+}$; *Isl1* $^{+/+}$ | NO DEFECT |
| E15.5 | 10 | EYP6-5 | *Nipbl* $^{+/+}$; *Isl1* $^{+/+}$ | NO DEFECT |
| E15.5 | 11 | EWH5-1 | *Nipbl* $^{+/+}$; *Isl1* $^{+/+}$ | NO DEFECT |
| E15.5 | 12 | EWH5-6 | *Nipbl* $^{+/+}$; *Isl1* $^{+/+}$ | NO DEFECT |
| E15.5 | 13 | FEY7-1 | *Nipbl* $^{+/+}$; *Isl1* $^{+/+}$ | NO DEFECT |
| E15.5 | 14 | FFA16-2 | *Nipbl* $^{+/+}$; *Isl1* $^{+/+}$ | NO DEFECT |
| E15.5 | 15 | FFA16-6 | *Nipbl* $^{+/+}$; *Isl1* $^{+/+}$ | NO DEFECT |
| E15.5 | 16 | FFA16-7 | *Nipbl* $^{+/+}$; *Isl1* $^{+/+}$ | NO DEFECT |
| E15.5 | 17 | FFA16-9 | *Nipbl* $^{+/+}$; *Isl1* $^{+/+}$ | NO DEFECT |
| E15.5 | 18 | FJP5-1 | *Nipbl* $^{+/+}$; *Isl1* $^{+/+}$ | NO DEFECT |
| E15.5 | 19 | FJP5-7 | *Nipbl* $^{+/+}$; *Isl1* $^{+/+}$ | NO DEFECT |
| E15.5 | 20 | FKX6-5 | *Nipbl* $^{+/+}$; *Isl1* $^{+/+}$ | NO DEFECT |
| E15.5 | 21 | FKX6-7 | *Nipbl* $^{+/+}$; *Isl1* $^{+/+}$ | NO DEFECT |
| E15.5 | 22 | FKI7-3 | *Nipbl* $^{+/+}$; *Isl1* $^{+/+}$ | NO DEFECT |
| E15.5 | 23 | FKI7-5 | *Nipbl* $^{+/+}$; *Isl1* $^{+/+}$ | NO DEFECT |
| E15.5 | 1 | EQU8-1 | *Isl1* $^{Cre/+}$ | NO DEFECT |

| | | | | |
|---|---|---|---|---|
| E15.5 | 2 | EQU8-3 | *Isl1* $^{Cre/+}$ | ASD OS |
| E15.5 | 3 | EQU8-4 | *Isl1* $^{Cre/+}$ | NO DEFECT |
| E15.5 | 4 | EQU8-7 | *Isl1* $^{Cre/+}$ | NO DEFECT |
| E15.5 | 5 | ERV5-1 | *Isl1* $^{Cre/+}$ | NO DEFECT |
| E15.5 | 6 | ERV5-5 | *Isl1* $^{Cre/+}$ | NO DEFECT |
| E15.5 | 7 | EUG7-1 | *Isl1* $^{Cre/+}$ | NO DEFECT |
| E15.5 | 8 | EUG7-2 | *Isl1* $^{Cre/+}$ | NO DEFECT |
| E15.5 | 9 | EWH5-3 | *Isl1* $^{Cre/+}$ | NO DEFECT |
| E15.5 | 10 | EWH5-4 | *Isl1* $^{Cre/+}$ | NO DEFECT |
| E15.5 | 11 | EXU5-4 | *Isl1* $^{Cre/+}$ | VSD |
| E15.5 | 12 | EXU5-6 | *Isl1* $^{Cre/+}$ | NO DEFECT |
| E15.5 | 13 | EXU5-7 | *Isl1* $^{Cre/+}$ | NO DEFECT |
| E15.5 | 14 | EYP6-6 | *Isl1* $^{Cre/+}$ | NO DEFECT |
| E15.5 | 15 | EZW16-1 | *Isl1* $^{Cre/+}$ | NO DEFECT |
| E15.5 | 16 | EZW16-4 | *Isl1* $^{Cre/+}$ | NO DEFECT |
| E15.5 | 17 | EZW16-7 | *Isl1* $^{Cre/+}$ | NO DEFECT |
| E15.5 | 18 | FEY7-3 | *Isl1* $^{Cre/+}$ | NO DEFECT |
| E15.5 | 19 | FEY7-5 | *Isl1* $^{Cre/+}$ | NO DEFECT |
| E15.5 | 20 | FFA16-3 | *Isl1* $^{Cre/+}$ | NO DEFECT |
| E15.5 | 21 | FFA16-8 | *Isl1* $^{Cre/+}$ | NO DEFECT |
| E15.5 | 22 | FKI7-2 | *Isl1* $^{Cre/+}$ | NO DEFECT |
| E15.5 | 1 | EQU8-6 | *Nipbl* $^{+/-}$ | NO DEFECT |
| E15.5 | 2 | ERV5-2 | *Nipbl* $^{+/-}$ | NO DEFECT |
| E15.5 | 3 | EUG7-5 | *Nipbl* $^{+/-}$ | NO DEFECT |
| E15.5 | 4 | EUH7-5 | *Nipbl* $^{+/-}$ | NO DEFECT |
| E15.5 | 5 | EUH7-6 | *Nipbl* $^{+/-}$ | NO DEFECT |
| E15.5 | 6 | EUH7-7 | *Nipbl* $^{+/-}$ | ASD OS |
| E15.5 | 7 | EZW16-2 | *Nipbl* $^{+/-}$ | NO DEFECT |
| E15.5 | 8 | EYP6-8 | *Nipbl* $^{+/-}$ | NO DEFECT |

| | | | | |
|---|---|---|---|---|
| E15.5 | 9 | EWH5-2 | *Nipbl* $^{+/-}$ | NO DEFECT |
| E15.5 | 10 | EWH5-5 | *Nipbl* $^{+/-}$ | NO DEFECT |
| E15.5 | 11 | FEY7-4 | *Nipbl* $^{+/-}$ | ASD OP |
| E15.5 | 12 | FFA16-5 | *Nipbl* $^{+/-}$ | NO DEFECT |
| E15.5 | 13 | FJP5-4 | *Nipbl* $^{+/-}$ | NO DEFECT |
| E15.5 | 14 | FJP5-5 | *Nipbl* $^{+/-}$ | NO DEFECT |
| E15.5 | 15 | FKX6-1 | *Nipbl* $^{+/-}$ | NO DEFECT |
| E15.5 | 16 | FKX6-2 | *Nipbl* $^{+/-}$ | ASD OS; TGA |
| E15.5 | 17 | FKX6-3 | *Nipbl* $^{+/-}$ | NO DEFECT |
| E15.5 | 18 | FKX6-6 | *Nipbl* $^{+/-}$ | NO DEFECT |
| E15.5 | 19 | FKI7-4 | *Nipbl* $^{+/-}$ | NO DEFECT |
| E15.5 | 20 | FKI7-6 | *Nipbl* $^{+/-}$ | ASD-OS; VSD |
| E15.5 | 1 | EQU8-2 | *Nipbl* $^{+/-}$; *Isl1* $^{Cre/+}$ | ASD OP; VSD; DORV |
| E15.5 | 2 | EQU8-5 | *Nipbl* $^{+/-}$; *Isl1* $^{Cre/+}$ | VSD; DORV |
| E15.5 | 3 | ERV5-3 | *Nipbl* $^{+/-}$; *Isl1* $^{Cre/+}$ | ASD OS |
| E15.5 | 4 | ERV5-4 | *Nipbl* $^{+/-}$; *Isl1* $^{Cre/+}$ | NO DEFECT |
| E15.5 | 5 | EUG7-4 | *Nipbl* $^{+/-}$; *Isl1* $^{Cre/+}$ | ASD OS |
| E15.5 | 6 | EUH7-4 | *Nipbl* $^{+/-}$; *Isl1* $^{Cre/+}$ | VSD; PTA |
| E15.5 | 7 | EZW16-3 | *Nipbl* $^{+/-}$; *Isl1* $^{Cre/+}$ | NO DEFECT |
| E15.5 | 8 | EZW16-6 | *Nipbl* $^{+/-}$; *Isl1* $^{Cre/+}$ | ASD OS |
| E15.5 | 9 | EYP6-2 | *Nipbl* $^{+/-}$; *Isl1* $^{Cre/+}$ | NO DEFECT |
| E15.5 | 10 | EYP6-7 | *Nipbl* $^{+/-}$; *Isl1* $^{Cre/+}$ | VSD; ASD OP |
| E15.5 | 11 | EYP6-9 | *Nipbl* $^{+/-}$; *Isl1* $^{Cre/+}$ | VSD; PTA |
| E15.5 | 12 | FEY7-2 | *Nipbl* $^{+/-}$; *Isl1* $^{Cre/+}$ | PTA,VSD |
| E15.5 | 13 | FEY7-6 | *Nipbl* $^{+/-}$; *Isl1* $^{Cre/+}$ | PTA,VSD |
| E15.5 | 14 | FEY7-7 | *Nipbl* $^{+/-}$; *Isl1* $^{Cre/+}$ | NO DEFECT |
| E15.5 | 15 | FEY7-8 | *Nipbl* $^{+/-}$; *Isl1* $^{Cre/+}$ | PTA, VSD, ASD OP |
| E15.5 | 16 | FFA16-1 | *Nipbl* $^{+/-}$; *Isl1* $^{Cre/+}$ | ASD OS |
| E15.5 | 17 | FJP5-2 | *Nipbl* $^{+/-}$; *Isl1* $^{Cre/+}$ | NO DEFECT |

| | | | | |
|---|---|---|---|---|
| E15.5 | 18 | FJP5-3 | *Nipbl* $^{+/-}$; *Isl1* $^{Cre/+}$ | ASD-OS |
| E15.5 | 19 | FJP5-6 | *Nipbl* $^{+/-}$; *Isl1* $^{Cre/+}$ | NO DEFECT |
| E15.5 | 20 | FKI7-1 | *Nipbl* $^{+/-}$; *Isl1* $^{Cre/+}$ | ASD-OP |
| E15.5 | 21 | FKI7-7 | *Nipbl* $^{+/-}$; *Isl1* $^{Cre/+}$ | NO DEFECT |

**Table S18.** Sample identifiers of hearts subjected to RNA sequencing in Fig. 7.

| Age | No | Sample | Genotype |
|---|---|---|---|
| E10.5 | 1 | IVF1-9 | Nipbl +/+; Isl1 +/+ |
| E10.5 | 2 | IVF1-11 | Nipbl +/+; Isl1 +/+ |
| E10.5 | 3 | IVF1-14 | Nipbl +/+; Isl1 +/+ |
| E10.5 | 4 | IVF5-15 | Nipbl +/+; Isl1 +/+ |
| E10.5 | 5 | IVF5-6 | Nipbl +/+; Isl1 +/+ |
| E10.5 | 6 | IVF6-8 | Nipbl +/+; Isl1 +/+ |
| E10.5 | 7 | IVF2-13 | Nipbl +/+; Isl1 +/+ |
| E10.5 | 8 | IVF6-5 | Nipbl +/+; Isl1 +/+ |
| E10.5 | 9 | IVF2-6 | Nipbl +/+; Isl1 +/+ |
| E10.5 | 10 | IVF2-18 | Nipbl +/+; Isl1 +/+ |
| E10.5 | 1 | IVF1-1 | Isl1 +/- |
| E10.5 | 2 | IVF1-6 | Isl1 +/- |
| E10.5 | 3 | IVF1-10 | Isl1 +/- |
| E10.5 | 4 | IVF2-4 | Isl1 +/- |
| E10.5 | 5 | IVF2-12 | Isl1 +/- |
| E10.5 | 6 | IVF5-1 | Isl1 +/- |
| E10.5 | 7 | IVF5-9 | Isl1 +/- |
| E10.5 | 8 | IVF5-14 | Isl1 +/- |
| E10.5 | 9 | IVF6-10 | Isl1 +/- |
| E10.5 | 1 | IVF5-7 | Nipbl +/- |
| E10.5 | 2 | IVF5-18 | Nipbl +/- |
| E10.5 | 3 | IVF2-10 | Nipbl +/- |
| E10.5 | 4 | IVF5-8 | Nipbl +/- |
| E10.5 | 5 | IVF5-2 | Nipbl +/- |
| E10.5 | 6 | IVF6-1 | Nipbl +/- |
| E10.5 | 7 | IVF6-2 | Nipbl +/- |
| E10.5 | 8 | IVF6-6 | Nipbl +/- |

| | | | |
|---|---|---|---|
| E10.5 | 9 | IVF2-9 | Nipbl +/- |
| E10.5 | 1 | IVF5-13 | Nipbl +/-; Isl1 +/- |
| E10.5 | 2 | IVF7-7 | Nipbl +/-; Isl1 +/- |
| E10.5 | 3 | IVF5-16 | Nipbl +/-; Isl1 +/- |
| E10.5 | 4 | IVF5-3 | Nipbl +/-; Isl1 +/- |
| E10.5 | 5 | IVF5-11 | Nipbl +/-; Isl1 +/- |
| E10.5 | 6 | IVF2-17 | Nipbl +/-; Isl1 +/- |
| E10.5 | 7 | IVF6-7 | Nipbl +/-; Isl1 +/- |
| E10.5 | 8 | IVF2-3 | Nipbl +/-; Isl1 +/- |
| E10.5 | 9 | IVF5-10 | Nipbl +/-; Isl1 +/- |

**Video S1.** Video of LSFM images captured through a whole $Nipbl^{Flox/+}$ heart. RV, right ventricle; LV, left ventricle; Ao, aorta; RA, right atrium; LA, left atrium; S, atrial septum; PT, pulmonary trunk.

**Video S2.** Video of LSFM images captured through a whole $Nipbl^{FIN/+}$ heart. RV, right ventricle; LV, left ventricle; Ao, aorta; RA, right atrium; LA, left atrium; S, atrial septum; PT, pulmonary trunk. Three defects are labeled: atrial septal defect (ASD), ventricular septal defect (VSD), and double outlet right ventricle (DORV).
